# In Vivo Network-Level Cerebrovascular Mapping Reveals the Impact of Flow Topology on Capillary Stalls After Stroke

**DOI:** 10.1101/2025.07.28.667165

**Authors:** Lukas Glandorf, Jeanne Droux, Etienne Jessen, Bastian Wittmann, Daewoon Song, Bruno Weber, Susanne Wegener, Bjoern Menze, Rainer Leitgeb, Dominik Schillinger, Mohamad El Amki, Daniel Razansky

## Abstract

Cerebral microvasculature is essential for brain function, but how flow and large-scale connectivity contribute to its resilience or failure remains poorly understood. To address this, we developed OMNIMap, a framework that integrates extended-focus optical coherence microscopy, learning-based segmentation and vessel-graph optimization, linking capillary flow and stalls to broader network context. Imaging over 60,000 capillaries in the mouse cortex revealed heterogeneous vascular responses to stroke, with capillary stalls, flow increases and decreases within the same networks. Divergent-to-convergent capillary subtypes had the highest stall risk. This association was not fully explained by differences in vessel structure, velocity or network position, suggesting that additional factors contribute to their vulnerability. Despite changes in individual capillary assignments after stroke, the overall proportions of capillary subtype remained largely preserved. Longitudinal mapping further showed that capillary stalls may persist until at least 7 days post-stroke. Together, these findings identify local flow topology as a marker of capillary vulnerability after stroke.

## INTRODUCTION

Maintaining cerebral blood flow is essential for brain function, delivering oxygen and nutrients while clearing metabolic waste. This feat is enabled by a sophisticated vascular network including arteries, veins and a mesh of dense and highly interconnected capillaries^1^. Disruptions to this capillary network are increasingly being recognized to contribute to a wide range of neurological conditions, including diabetes^2^, Alzheimer’s disease^3,4^, vascular dementia^5^, and stroke^6–9^. In aging and chronic neurological disease^10^, capillary stalls have been associated with leukocyte adhesion, other cellular obstructions, and changes in capillary diameter ^2,11–13^. Stroke is a leading cause of long-term disability, where even brief flow interruptions can trigger irreversible neuronal loss^14^. Beyond the infarct core, capillary dysfunction also limits tissue recovery in the surrounding penumbra, contributing to the “no-reflow” phenomenon that undermines the benefits of recanalization therapies^15,16^.

Despite its critical role, much remains unknown about how capillary connectivity and flow organization shape network resilience or failure. Why do certain capillaries stall while others remain resilient? Does capillary vulnerability arise from proximity to arterioles or venules, or is it governed by deeper principles of flow architecture? How well can the microvascular network compensate for input flow perturbations? Answering such questions requires tools capable of imaging the functional microvascular network at scale. Much of our understanding of microvascular failure relies on two-photon microscopy^17,18^ (2PM). 2PM provides high-resolution time traces of red blood cell flux in individual capillaries, but mapping flow across their wider connections is difficult when measurements sample selected vessels. Large-volume angiography reveals extensive vascular networks^19,20^, while recent light-field imaging can track blood cells across thousands of connected branches^21^. Conversely, mesoscale modalities^22–27^ access larger volumes but lack the resolution to resolve individual capillary connections and flow. These advances make network-scale measurements possible, but how local flow organization shapes capillary failure and recovery after stroke remains challenging and unresolved. Addressing this question requires longitudinal measurements of vascular connectivity, morphology and hemodynamics simultaneously in the same network.

To address this, we developed OMNIMap (Optical Coherence Microscopy for functional Microvascular Network Imaging and Mapping), a framework for *in vivo* volumetric mapping of cerebral microvascular networks. OMNIMap integrates extended-focus optical coherence microscopy with deep-learning segmentation and vessel graph optimization to quantify flow velocities and directions from pial vessels down to individual capillaries^28^ and resolve artery-vein identity and branching order. This enables us to relate capillary flow and stalling to the broader vascular architecture. Using OMNIMap we mapped flow topology, velocity changes and capillary stalls in the capillary network after stroke. We found that the most stall-prone capillary subtype is also the most frequent. Throughout the network, subtype proportions remain broadly similar although individual capillaries change subtype, while acute stalls predict later flow loss. Together, these findings connect local flow topology to microvascular vulnerability and offer a new lens through which to study microcirculatory failure.

## Results

### Extracting morphology and functional blood flow of microvascular networks

One major challenge in intravital cerebral blood flow imaging is attaining 3D quantitative vascular imaging with sufficient spatial resolution to enable visualization down to the smallest capillaries while rendering accurate microvascular connectivity maps over large brain areas. OMNIMap employs xfOCM for *in vivo* image acquisition, which integrates Bessel beam illumination and Gaussian beam detection with a dedicated data processing pipeline^28,29^. It can capture volumetric snapshots of the microvascular network at millimeter scale (1000 × 1000 × 360 μm^3^), thus enabling the study of highly interconnected microvascular network patches, including the feeding and draining pial vessel trees, penetrating arterioles and ascending venules, as well as the capillary bed (Fig. 1a). To extract hemodynamic information, we combined structural OCTA with deep learning-based segmentation (Supplementary Fig. 1) and functional DOCT (Fig. 1b: see Methods for details). The measured lateral resolution is 3 µm and the theoretical axial resolution approximately 4 µm in tissue. Repeated volumes are averaged over approximately 5 minutes for OCTA and 30 minutes for DOCT. Using OMNIMap, we obtained highly detailed structural reconstructions of the 3D microvascular networks along with their local flow velocity profiles (Fig. 1c). These data allow quantitative assessments of morphological parameters such as diameters (Fig. 1d,e) and blood flow velocities (Fig. 1f,g) for each blood vessel segment between branching points (acquisition cohorts in Supplementary Table 1). Both distributions are dominated by the high number of thin and slow-flowing capillaries, resulting in a mean of animal median diameter of 3.85 ± 0.10 µm consistent with previously described capillary diameters^30,31^ and mean of median blood flow velocities of 1.17 ± 0.36 mm·s^-1^. We provide a comparison of baseline values with previously published studies in Supplementary Table 2. Segmentation accuracy and measurement validation are summarized in Extended Data Fig. 1.

**Figure 1:**
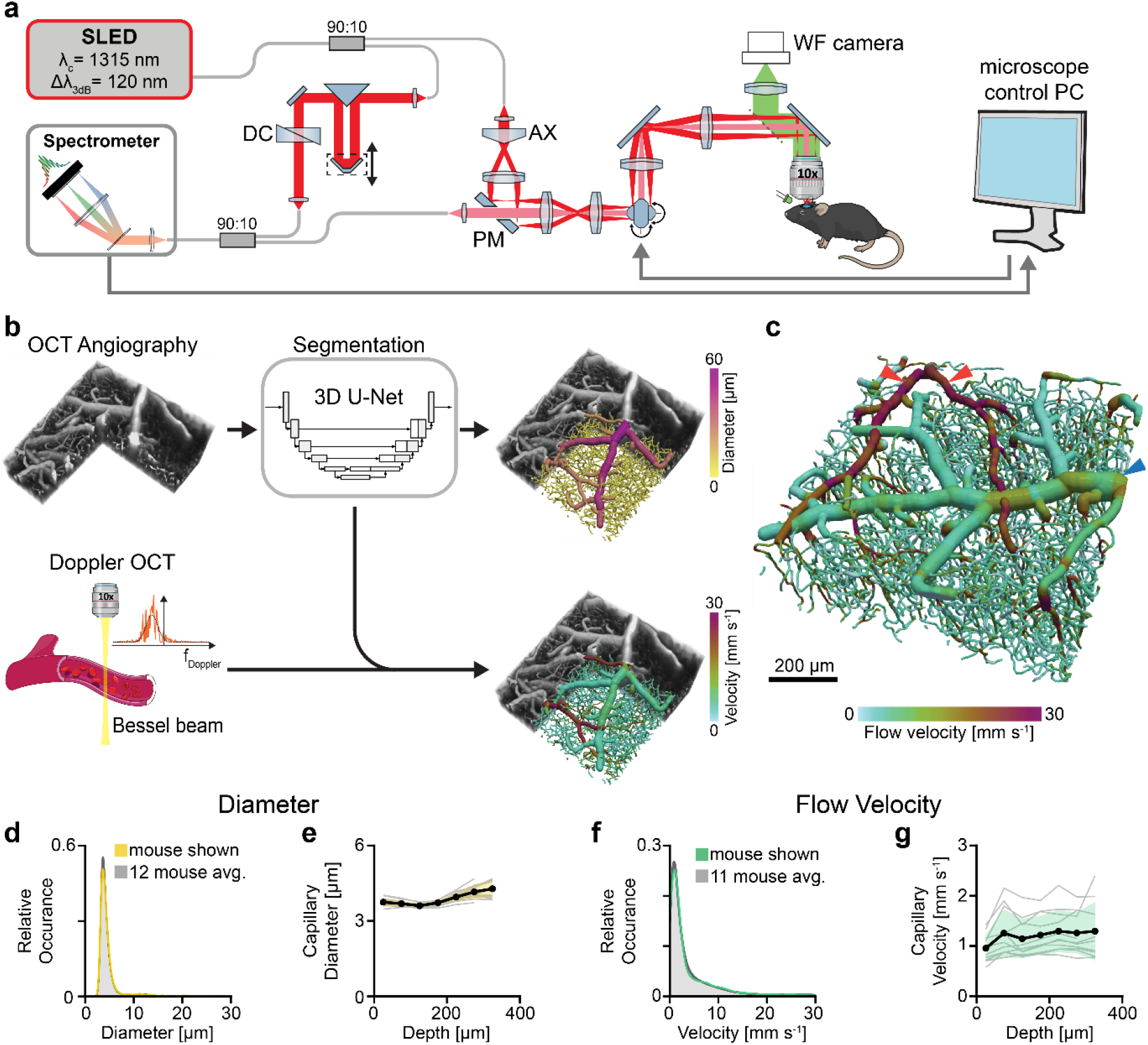
Acquisition and morphological/functional analysis of large-scale microvascular networks in vivo. (a) Schematic of the extended-focus optical coherence microscope (xfOCM) configured as a Mach-Zehnder interferometer (PM: pierced mirror, AX: axicon, DC: dispersion compensation glass, WF: wide-field camera). (b) Acquisition and processing workflow: optical coherence tomography angiography (OCTA) and Doppler OCT (DOCT) volumes are collected in separate scans, followed by a tailored 3D U-Net for blood vessel segmentation in the OCTA volume. A skeletonization step extracts vessel centerlines, enabling the calculation of vessel diameters and flow velocities for each segment. (c) 3D rendering of the imaged microvascular network with diameter and velocity distributions corresponding to the individual curves in (d) and (f). Downstream branches of the middle cerebral artery (MCA) can be identified by their high blood flow velocities and relatively narrow vessel structure (red arrows). Major pial vein tree segments in the volume tend to exhibit larger diameters but slower blood flow (blue arrows). Blood vessel connectivity is represented in an edge graph where each vessel bifurcation point is represented as a node and blood vessel segments between bifurcation points as connecting edges. Scale bar = 200 µm. (d) Vessel diameter distributions from a representative animal (yellow) and a 12-animal average (gray). Diameters exceeding 30 µm are truncated for clarity to emphasize capillary ranges. (e) Capillary diameter by cortical depth. (f) Flow velocity distributions measured across the same regions shown in (d), with green depicting a single dataset and gray showing the mean of 11 animals. (g) Capillary velocity by cortical depth. In e,g, gray lines show individual mice; black lines and shading show the mean ± SD of animal medians in 50-µm bins from 0 to 350 µm. Measurement validation is shown in Extended Data Fig. 1.

### Arterial-venous imbalance in automatically labeled cortical microvasculature

Next, we evaluated the ability of our method to specifically label arteries, arterioles, capillaries, venules and veins. Commonly employed manual artery-vein labeling becomes intractable for large or complex 3D data, particularly when considering vertically oriented penetrating arterioles and ascending venules, that gradually transition into the dense capillary bed. To address this, we developed a vessel graph-based algorithm that leverages blood flow direction extracted from the DOCT data (Supplementary Fig. 2) and known vessel connectivity to automatically classify each vessel as a pial artery, penetrating arteriole, capillary, ascending venule, or pial vein (Fig. 2; Supplementary Fig. 3). First, all pial vessels were automatically identified based on their diameter and near-horizontal orientation in the cortical plane (Fig. 2a). Some incorrect connectivity assignments at closely crossing pial vessels (Fig. 2a, green arrow) and erroneously flipped flow directions (Fig. 2a, magenta arrow) have been identified, correlating well with strongly increased or decreased normalized mass flow differences calculated at each bifurcation. To counteract this challenge that prevents direct graph traversal, we devised an algorithm that first identifies penetrating arterioles and ascending venules by their vertical orientation and flow direction as reliable information points (Fig. 2b). Then, the pial vessel network was optimized by minimizing the normalized mass balance through flow direction changes, edge deletions and small diameter corrections (Fig. 2c). Finally, we identified pial vessel trees that are dominated by divergent distribution of the blood towards smaller vessels (pial arteries) as well as primarily characterized by convergent collection of the blood (pial veins, Fig. 2d). The optimization substantially reduces normalized mass flow errors across animals, and four-way connections in the pial vasculature that occur at falsely identified interconnections (Extended Data Figure 2g).

**Figure 2:**
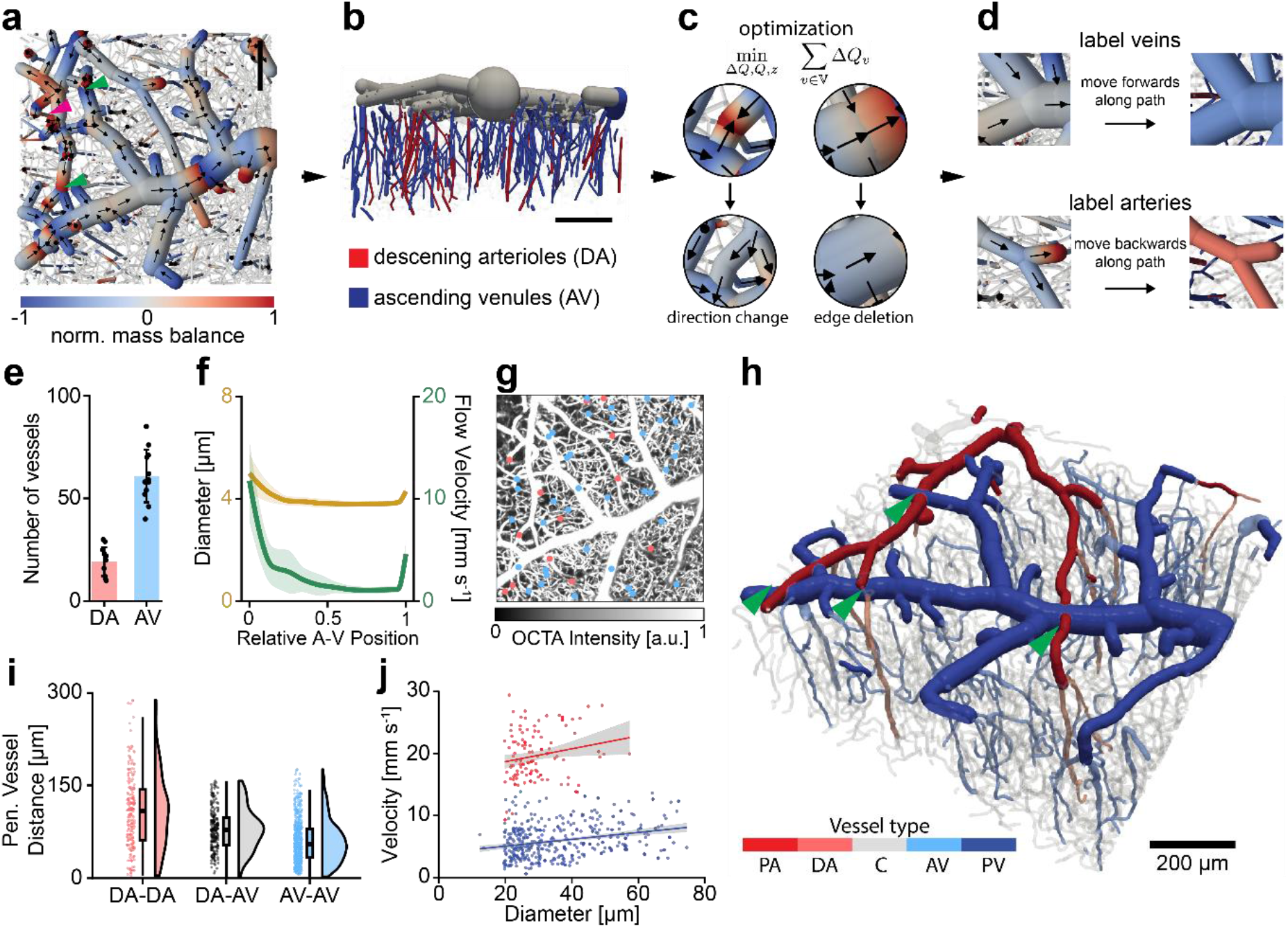
Automated classification of arterial and venous vessel trees in the vascular network. (a) Top-down view of the normalized mass balance for the pial vessel network, with each vessel segment simplified as a cylinder. The closer the mass balance is to zero, the more similar the in- and outflow at a branch point; values near ±1 denote highly unbalanced flows. Black arrows indicate the blood flow direction. (b) Side view highlighting penetrating arterioles (red) and ascending venules (blue) based on flow direction and angle relative to the pial surface; pial vessels are shown in gray, and capillaries are omitted for clarity. (c) Illustrative examples of flow direction changes and edge deletions implemented during the flow optimization procedure. (d) Representative converging branch point (top, blue) identified as a pial vein and a diverging branch point (bottom, red) labeled as a pial artery. (e) Counts of descending arterioles (DA) (19.25 ± 7.00) and ascending venules (AV) (60.92 ± 12.80) per mm² across twelve mice (N=12; mean ± SD). Counts are connected components within the imaged field; points show mice and bars show means ± SD. (f) Diameter and velocity along relative arterial-venous position. Lines and shading show means ± SD of animal medians (12 mice for diameter, 11 for velocity). (g) Maximum-intensity projection (MIP) of the OCTA volume, overlaid with the starting positions of penetrating arterioles (red) and ascending venules (blue). (h) Rendered microvascular network patch showing vessel-type labels (PA: pial artery, DA: descending arteriole, C: capillary, AV: ascending venule, PV: pial vein). Green arrows highlight crossing points where pial arteries and veins run in proximity but are correctly differentiated by the optimization algorithm. Scale bar: 200 µm. (i) Lateral distance distributions between starting points of penetrating vessels: arteriole-to-arteriole (red), arteriole-to-venule (gray), and venule-to-venule (blue). Each penetrating vessel’s distance to its nearest neighbor is included. (N=12). (j) Blood flow velocities plotted against vessel diameter for pial arteries (red) and pial veins (blue), with regression lines and 95% confidence intervals.

We found an average of 19 ± 7.00 descending arteriole (DA) and 60 ± 12.80 ascending venule (AV) trees per square millimeter, respectively (Fig. 2e). These densities and resulting 3.16 times higher number of ascending venules than penetrating arterioles align well with previously reported values^1,32^. Diameter and flow velocity decreased from DAs into capillary bed and increased toward drainage through AVs (Fig. 2f). Furthermore, their spatial arrangement echoes the characteristic column structure of one DA being surrounded by multiple AVs in the cortical surface (Fig. 2g).

The resulting microvascular networks exhibit clear partitions between arterial and venous trees, both in the pial layer and in their vertically oriented penetrating or ascending vessels (Fig. 2h). Even at critical crossing points where pial arteries and veins run in proximity, the optimized network depicts the correct connectivity and vessel-type classification (green arrows, Fig. 2h). Consistent with the higher number of AVs in the cortex, the mean of animal median distances between each DA and its closest AV was 79.30 ± 13.28 µm, compared with 56.48 ± 5.87 µm between each AV and its closest AV neighbor (Fig. 2i). In contrast, the mean of animal median distances between each DA and its closest DA neighbor was 110.95 ± 19.21 µm. Functionally, pial arteries exhibit higher flow velocities than similarly sized pial veins and manifest a trend of increasing velocities with larger vessel diameters (Fig. 2j).

### Morphological and flow-based metrics of stroke damage in the cortical penumbra

We then characterized the morphological and functional changes in the imaged microvascular network following the occlusion of the main supplying artery, the middle cerebral artery (MCA). We used a stroke model consisting of thrombin injection into the MCA, which leads to *in situ* clot formation (Fig. 3a). In each mouse, we imaged the same cortical volume through a cranial window one day prior and approximately 1 hour after the stroke induction (Fig. 3b). We then confirmed that the imaged area is situated within the penumbra using laser speckle contrast imaging^33^ (LSCI, Fig. 3c,d).

**Figure 3:**
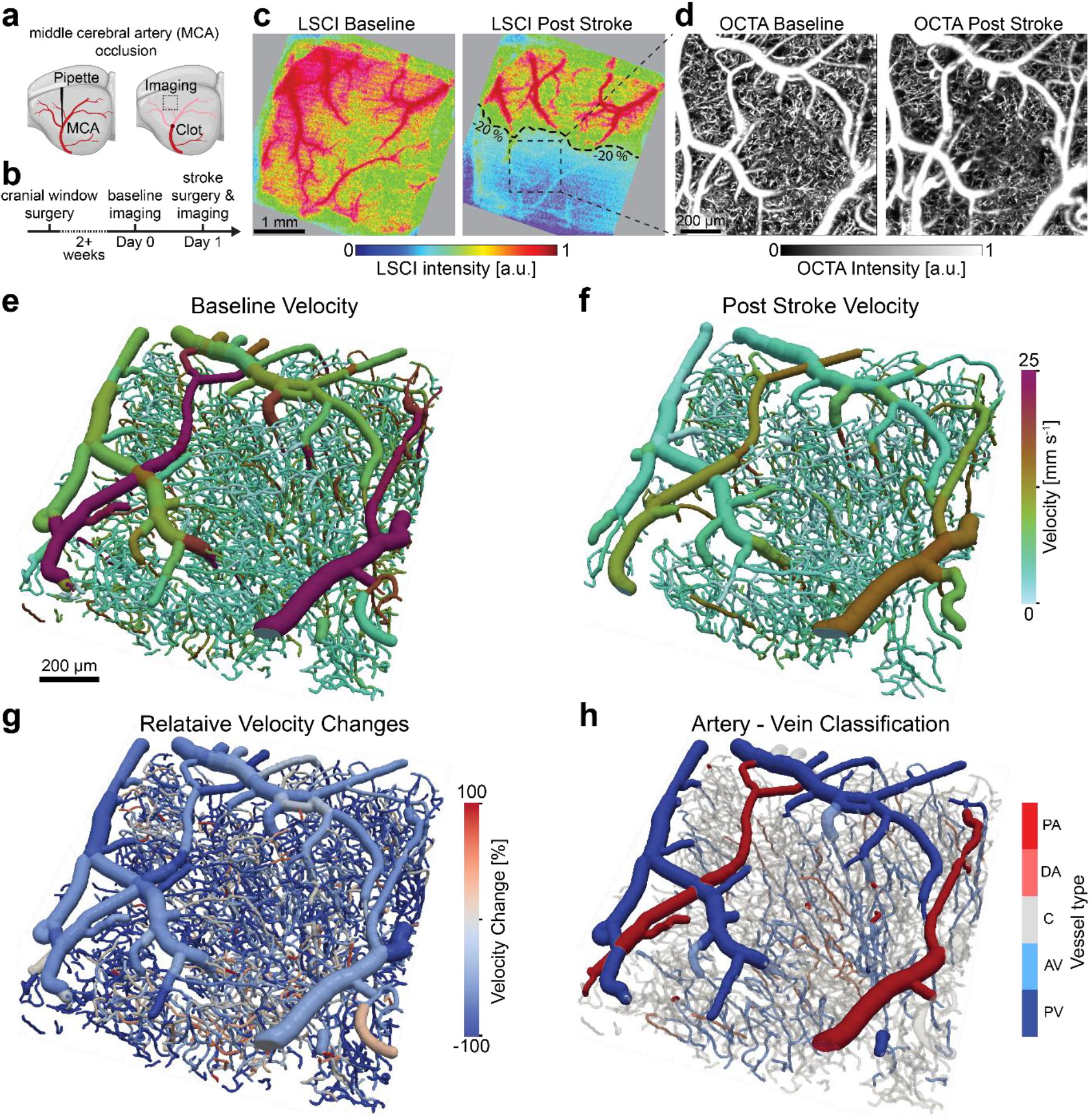
Quantitative in vivo imaging of the microvascular network response to stroke. (a) Schematic of the middle cerebral artery (MCA) occlusion procedure. Thrombin is injected into the M2 segment of the MCA to induce localized clot formation. (b) Experimental timeline, illustrating cranial window surgery, baseline imaging, stroke induction by Thrombin injection, and post-stroke imaging. (c) Laser speckle contrast imaging (LSCI) example at baseline and post-stroke, with the dashed square indicating the OCT imaging field of view. The contour marks a 20% perfusion reduction. Scale bar, 1 mm. (d) Maximum-intensity projection (MIP) of the OCTA volume at baseline and post-stroke from the same region shown in (c). Scale bar, 200 µm. (e, f) Representative microvascular network patches overlaid with color-coded flow velocities at baseline (e) and post-stroke (f). (g) Relative velocity changes in each vessels compared to baseline, with capillary stalls displayed at -100% for visualization. (h) Example of vessel-type labels (pial arteries in dark red, descending arterioles in light red, capillaries in gray, ascending venules in light blue, pial veins in dark blue) applied to the same microvascular network. Regional comparisons are shown in Extended Data Fig. 3.

We observed the impact of stroke when comparing the morphological and hemodynamic alterations before and after stroke (Fig. 3e,f). Detectable capillary length decreased by 27.6%. Flow velocities shifted toward lower values over much of the network, while the relative-change map also showed apparent local increases (Fig. 3g). The vessel-class rendering identifies pial arteries and veins, descending arterioles, ascending venules and capillaries in the same network (Fig. 3h).

Next, stroke-induced changes across vessel-type were conducted (Fig. 4a-f). On a morphological level, we observed that after MCA occlusion, pial arteries and veins in the penumbra showed no clear paired changes in diameter, nor did capillaries (Fig. 4a-c). Descending arterioles showed an estimated 7.2% diameter increase, while ascending venules showed no clear change (Extended Data Fig. 4a,b). Pial arteries exhibited a mean velocity drop of 31.1%, while pial veins, already characterized by slower flow, declined by 12.6% (Figs. 4d,e). Descending arterioles and ascending venules showed a similar pattern of decreased flow, albeit to a lesser degree than their pial counterparts (DA: -4.9 %, AV: -21.4%; Extended Data Fig. 4c,d). Among capillaries with measurable velocity at both imaging time points, the estimated change was -22.8% (Fig. 4f). Initially slow capillaries were more likely to accelerate after stroke, beyond the pattern seen in independent baseline-frame controls (Supplementary Fig. 4). Accelerations showed no consistent association with nearby stalls or bypass routes (Supplementary Fig. 5).

**Figure 4:**
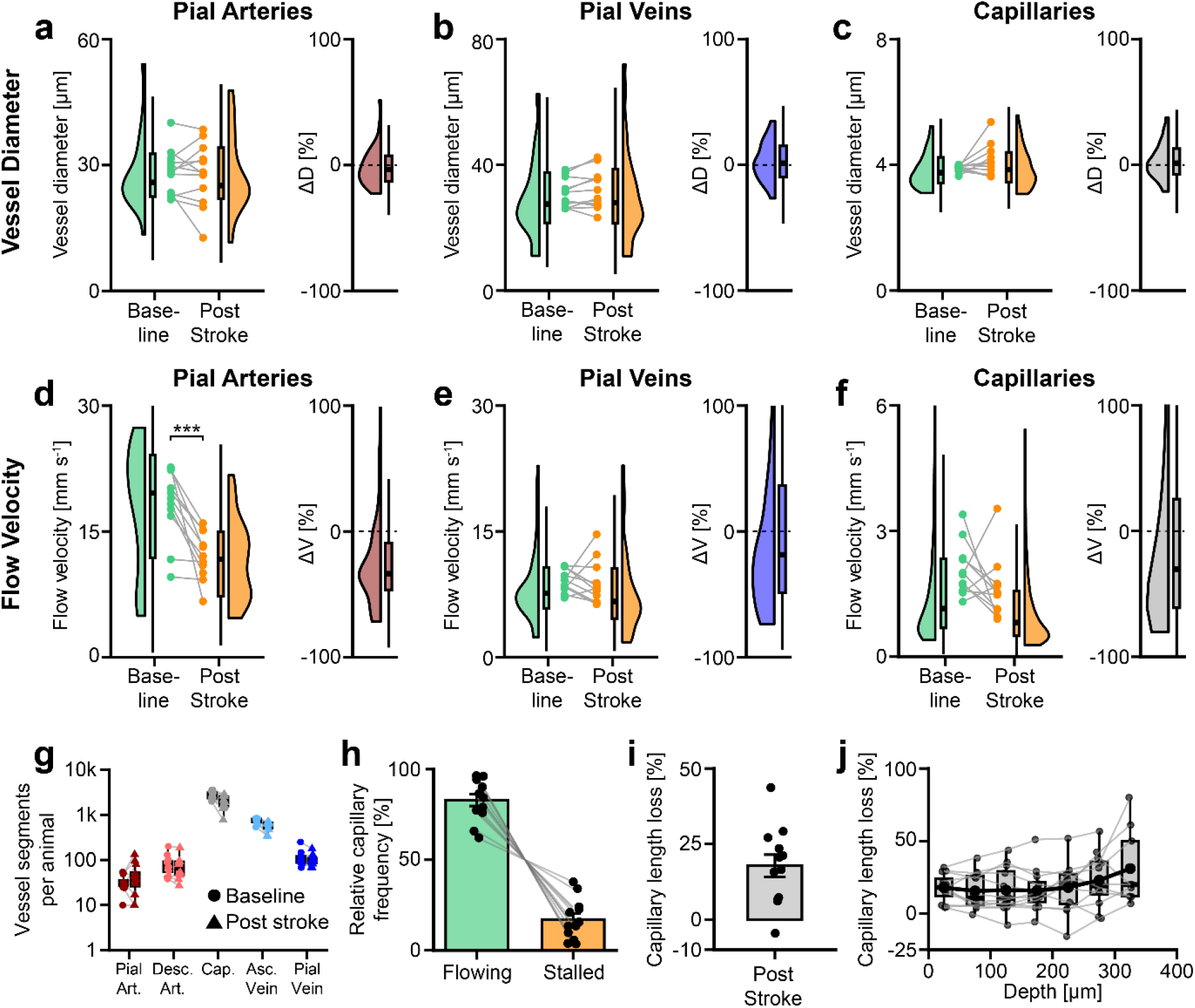
Vessel-type-specific responses to stroke across multiple animals. (a-c) Diameter measurements (baseline vs. post-stroke) and relative diameter changes for pial arteries (a), pial veins (b) and capillaries (c), showing no significant paired changes (N=12). (d-f) Flow velocities and relative velocity changes for pial arteries (d), pial veins (e) and capillaries (f). Paired changes were -31.1% (95% CI: -48.0 to -8.9%, P = 0.0141), -12.6% (−27.5 to +5.3%, P = 0.139) and -22.8% (−46.8 to +12.1%, P = 0.154), respectively (N=11). (g) Number of vessel segments, grouped by vessel type, at baseline and post-stroke (N=12). (h) Fraction of flowing (green) versus stalled (orange) capillaries among baseline vessels with resolved post-stroke fate (N=12). Mean stalled fraction, 17.1%; stalls denote loss of detectable OCTA perfusion. (i,j) Difference between the fraction no longer detected after stroke and the fraction detected only after stroke, using each direction’s assessable denominator, overall (i) and by depth (j). Mean difference, 17.8 percentage points (95% CI: 9.6-25.9). In a-f, measurements are paired across imaging time points; rainclouds show vessel distributions and connected points show animal means. Boxes show medians and IQRs, with 1.5 × IQR whiskers. Outer 5% tails are omitted from densities only; axis limits do not restrict tests. In h,i, bars show means ± SEM; in j, gray lines show mice and black their mean. Paired-change tests: two-sided t-tests of animal median log ratios. Distribution tests: whole-animal sign flips. P values are unadjusted; *** denotes distribution P < 0.001 in d (P = 0.000977). Penetrating vessels are shown in Extended Data Fig. 4.

The capillary bed accounts for most vessel segments in the cortex (Fig. 4g) and contributes substantially to cortical flow resistance^34^. During and after stroke, blood flow in many capillaries ceases through several mechanisms, including blockage by neutrophils, platelets, red blood cells or microthrombi, creating capillary stalls^15,35^. In this study, however, we use stall as an imaging-based operational classification rather than a mechanistic designation. Specifically, a capillary was classified as stalled when a capillary perfused and detectable by OCTA at baseline showed no detectable OCTA perfusion after stroke (Supplementary Fig. 6). This definition therefore captures acute loss of detectable capillary flow following arterial occlusion, irrespective of the underlying cellular mechanism. As a result, the capillary bed endures both decreased blood inflow and acute network consolidation from stalled capillaries. On average, 17.1% of capillaries stalled in the penumbra (Fig. 4h), consistent with previous smaller-scale findings^15,20^. Variations between animals likely stem from differences in angioarchitecture, stroke severity, and distance to the stroke core. As a separate control that does not require individual capillary matching, the fraction of capillaries no longer detected after stroke exceeded the fraction detected only after stroke by 17.8 percentage points on average (Fig. 4i). This net difference showed no clear monotonic trend with depth (Fig. 4j), while acute signal controls showed no progressively worsening deficit through 350 µm (Supplementary Fig. 7). Severely affected LSCI regions had a trend towards higher estimated stall fraction than moderately affected regions. Velocity and diameter changes showed no clear regional differences (Extended Data Fig. 3).

### Venous side capillary stalls are more numerous but not more likely

Capillary stalling occurs through multiple mechanisms, including changes in blood flow or tone^36,37^, increased adhesion proteins on endothelial cells and leucocytes^3,35^, as well as a dysfunction of the glycocalyx^5^. Earlier studies have begun exploring whether a capillary’s position within the microvascular network affects its susceptibility to stalling^38^. However, these investigations have been limited by the inability to map large-scale networks without involving labor-intensive manual tracing.

We compared the same vessels at baseline and post-stroke and defined stalls by loss of detectable OCTA perfusion along baseline capillaries (Fig. 5a), revealing capillary stalls throughout the entire microvascular network (Fig. 5b). Significant capillary loss was quantified through the cumulative perfused capillary length (Figs. 5c,d), exhibiting a drop of 28.4% that reflects an impaired delivery of oxygen and glucose throughout the network. This total-length change is distinct from the fraction of baseline capillaries classified as stalled (Fig. 4h). Segment counts can also change when the perfused graph is reconstructed: if a capillary side branch stalls, the two remaining segments at its former junction may merge into a single graph segment, contributing to an overall decline in the apparent number of capillary segments (Supplementary Fig. 8). Among stalled capillaries, median baseline segment length was 43.3% greater than in those that remained perfused (Supplementary Fig. 9).

**Figure 5:**
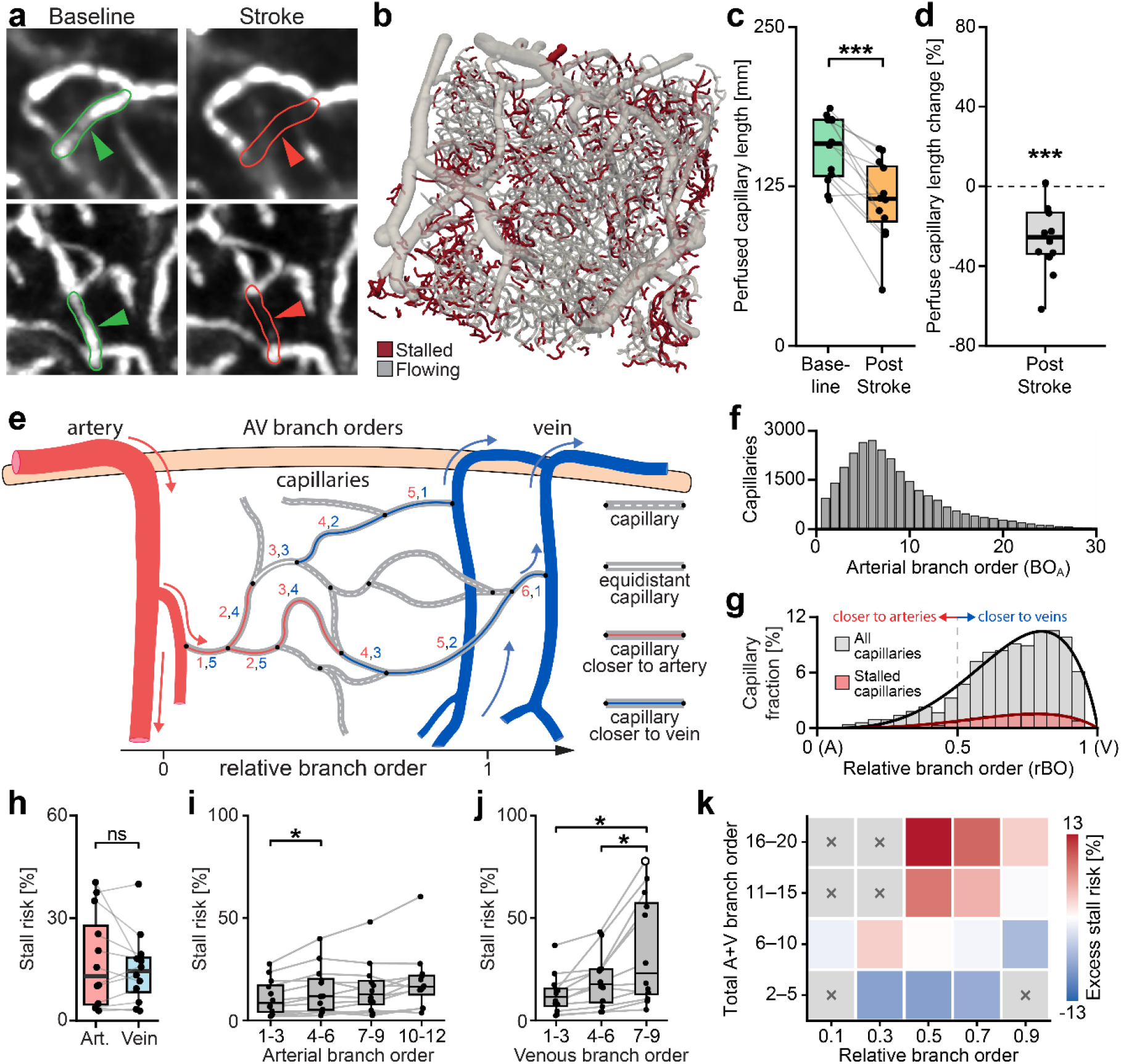
Mapping capillary stalls and branching order in the microvascular network. (a) Zoomed-in OCTA maximum-intensity projection (MIP) demonstrating capillary stall identification by comparing baseline (green outline/arrows) and stroke (red outline/arrows). Stalling capillaries are identified by loss of detectable OCTA perfusion. (b) Baseline microvascular network (MVN) patch highlighting capillaries that eventually stall (red) against all remaining vessels (gray). (c, d) Cumulative length of all perfused capillaries (c) and relative loss of perfused capillary length (d) from baseline to post-stroke (N=12). Mean loss, 28.4% (95% CI: 15.7-39.3%; two-sided paired log-ratio t-test, P = 0.000916, paired Hedges g_z = -1.21). (e) Schematic of arterial branching order (red) and reverse branching order from the venous side (blue), illustrating two shortest paths from an artery to separate veins. Relative branch order (rBO), arterial order divided by the sum of arterial and venous orders, spans (0,1), enabling comparisons across differently sized paths through the MVN. (f) Histogram of arterial capillary branching orders at baseline for 12 animals. (g) Relative capillary fraction histogram over rBO from artery (0) to vein (1). Baseline capillary fractions (gray) and fractions of stalled capillaries post-stroke (red) are compared. N=18,594 baseline capillaries with both orders and resolved fate. Both series use the baseline denominator and equal-animal averaging. (h) Stall incidence, calculated as the ratio of stalled to total capillaries, for arterial (rBO < 0.5, red) and venous (rBO > 0.5, blue) regions. Mean risks, 17.5% and 15.1%; venous-minus-arterial difference, -2.4 percentage points (95% CI: -7.4 to +2.6; two-sided paired t-test, P = 0.314, N=12). Equal-order capillaries are omitted. (i,j) Stall incidence by arterial (i) and venous (j) branch order. Tests compare paired animal risks with Holm correction across nine bin pairs. (k) Equal-animal excess stall risk, in percentage points, by relative position and total arterial-plus-venous order (2-5, 6-10, 11-15, 16-20). Gray cells lack ten capillaries per mouse in at least six mice. N=12. Boxes show medians and IQRs, with 1.5 × IQR whiskers. ns: P ≥ 0.05; *: P < 0.05; ***: P < 0.001.

Next, we classified each capillary with an arterial branching order based on its shortest connection to a DA (Figs. 5e, red numbers). Likewise, a reverse branching order quantifies the shortest distance from the venous tree (Fig. 5e, blue numbers). To compare paths of varying lengths, we introduce a relative branching order (rBO) on the interval (0,1), calculated by dividing a capillary’s arterial branching order with the sum of its arterial and venous branching orders. We found that arterial branching orders peak between 6 and 8 (Fig. 5f), aligning with previously reported *in vivo* results^37–39^. We quantified the accuracy of branch-order assignment within the experimental field-of-view using cropped reference vascular networks (Extended Data Fig. 2; Supplementary Fig. 10).

We next addressed the question of whether capillaries closer to veins (rBO > 0.5) are more prone to stalling (Fig. 5g). We observed that the fraction of stalled capillaries near the venous side is significantly higher than that near the arterial side (rBO < 0.5) across animals (83.2% versus 11.9%, respectively; Fig. 5g; Supplementary Fig. 11). However, the distribution of stalls follows the overall distribution of capillaries, which is skewed toward the venous tree (Fig. 5g, gray), due to the greater number of ascending venules compared to penetrating arterioles in the mouse brain (Fig. 2g). As a result, 82.6% of all pre-stroke capillaries lie closer to veins, 11.0% closer to arteries, and the remaining 6.4% have an rBO of exactly 0.5. When we computed stall incidence by taking the ratio of stalled to total capillaries in each category, there was no significant difference (p=0.314) between the arterial and venous sides (Fig. 5h). These data indicate that while there are more stalls near veins, we found no evidence of higher stall risk among capillaries closer to veins.

### Capillaries farther from penetrating vessels are more likely to stall

Relative branch order describes a capillary position along the arterial-to-venous axis but not its absolute branch distance from penetrating vessels. We therefore analyzed arterial and venous branch orders separately, grouping orders 1-3, 4-6 and 7-9 on both sides with an additional arterial group for orders 10-12 (Fig. 5i,j). Stall risk was higher at arterial branch orders 4-6 than at orders 1-3 (Fig. 5i). On the venous side, orders 7-9 had higher stall risk than orders 1-3 and 4-6, (Fig. 5j). Velocity decreased less at arterial branch orders 4-6 than at orders 1-3 (Extended Data Fig. 5a). Diameter changes also showed no clear differences between these groups on either side (Extended Data Fig. 5c,d). Finally, we combined relative arterial-to-venous position with total branch order (Fig. 5k). Within the central and venous portions of the arterial-to-venous axis, capillaries with total orders 11-20 had higher stall risk than those with orders 2-10.

### Topological capillary subtypes shaping microvascular connectivity and capillary stalls

To examine how the microvascular network consolidates after stroke, we leveraged our comprehensive data set that includes blood flow directions for all capillaries. We classified capillaries into four subtypes based on their distinct inflow-outflow patterns. Similar to previous *in silico* work, each capillary connecting to two other capillaries at its input and two at its output has been evaluated^40^. At each junction, capillaries can diverge (one inflow feeding two outflows) or converge (two inflows merging into one outflow). The central capillary linking two junctions was classified as divergent-to-divergent (D-to-D, Fig. 6a), divergent-to-convergent (D-to-C, Fig. 6b), convergent-to-divergent (C-to-D, Fig. 6c), or convergent-to-convergent (C-to-C, Fig. 6d).

**Figure 6:**
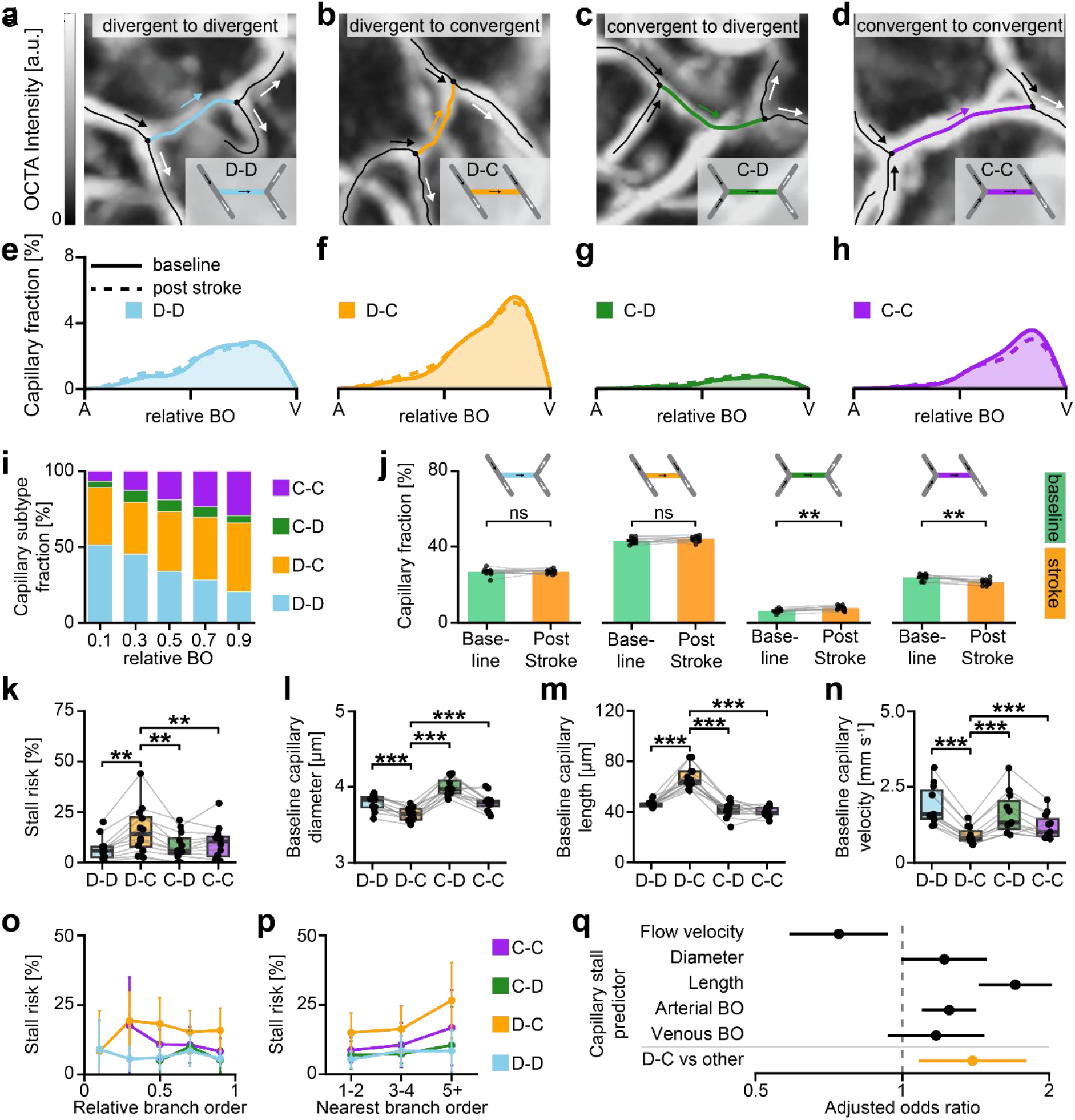
Topological capillary subtypes differ in morphology, flow and stall risk. (a–d) Partial OCTA maximum-intensity projections (MIPs) illustrating the four topological capillary subtypes. Each subtype is defined by the inflow (black arrows) and outflow (white arrows) patterns at its two junctions: Divergent-to-Divergent (D-D, blue), Divergent-to-Convergent (D-C, orange), Convergent-to-Divergent (C-D, green), Convergent-to-Convergent (C-C, purple). In a divergent connection, one inflow splits into two outflows; a convergent connection merges two inflows into one outflow. (e–h) Relative distributions of each topological subtype (D-D, D-C, C-D, C-C) across relative branch order (rBO). Solid lines represent baseline, dashed lines post-stroke. (i) Baseline subtype composition within relative-position bins. (j) Total fraction of each subtype per animal (N=12), before (green) and after (orange) stroke. Bars show means ± SEM. Paired t-tests show increases of 1.4 percentage points for C-D and decreases of 2.6 points for C-C (Holm-adjusted P = 0.002 each across four comparisons). D-D and D-C remain non-significant. (k) Subtype-specific stall incidence among 11,578 capillaries in 12 mice: mean risks, 6.8%, 16.1%, 8.2% and 9.9% for D-D, D-C, C-D and C-C. D-C exceeds each other subtype (two-sided paired t-tests, all Holm-adjusted P < 0.01 across six pairs). (l-n) Baseline diameter (l), segment length (m) and flow velocity (n) by subtype (N=12 for diameter/length, N=11 for velocity). D-C capillaries are narrower, longer and slower than each other subtype (two-sided paired log-ratio t-tests, all P < 0.001 after Holm correction across six pairs per metric). (o,p) Subtype-specific stall incidence by relative position and nearest penetrating-vessel order (1-2, 3-4, 5+). Points and error bars show equal-animal means and pointwise 95% CIs. (q) Adjusted stall odds ratios from animal-specific models combined across 10 mice. Models include all displayed predictors, tissue depth and boundary proximity. Points and bars show random-effects estimates and Knapp–Hartung 95% CIs. Continuous effects are per within-animal SD. Diameter, length and velocity are log-transformed. D-C versus other: odds ratio 1.39 (95% CI: 1.08-1.80; P = 0.028, Holm-adjusted across two focal coordinate-model tests). In k-n, points show mice and boxes show medians and IQRs, with 1.5 × IQR whiskers. All t-tests are two-sided. ns: P ≥ 0.05; **: P < 0.01; ***: P < 0.001.

Next, we mapped the distribution of each topological capillary subtype within the brain cortex (Figs. 6e - 6h). We observed that D-to-C is the most abundant capillary subtype (Fig. 6f) while C-to-D is the least common. These data show that diverging connections originate near the arterial tree and disperse throughout the network, though they diminish earlier toward the venous side (Fig. 6e). In contrast, converging connections are skewed toward the venous tree, where they consolidate the microvascular network (Fig. 6h). Notably, the distributions of the different capillary subtypes overlap substantially (Fig. 6i), suggesting that the brain microvascular network does not merely expand in the first few branching orders, then consolidating near the veins. Instead, diverging, static, and converging patterns intermix throughout the network.

Moreover, stroke did not substantially alter the distribution of capillary subtypes in the affected microvascular network (Fig. 6e-h). Subtype fractions also remain stable across animals (Fig. 6j). Although C-to-D increased and the C-to-C subtype decreased, their differences in mean capillary fraction remain below 3 percentage points.

We next tested whether capillary stalls risk differed among capillary subtypes. D-to-C capillaries had the highest stall risk, while D-to-D capillaries had the lowest and were relatively more common toward the arterial side (Fig. 6i,k). We therefore asked whether the higher risk in D-to-C capillaries could be explained by differences in their structural, hemodynamic, or network properties. At baseline, D-to-C capillaries were slower, smaller in diameter, and longer than the other subtypes (Fig. 6l-n).

After stroke, D-to-C capillaries remained longer and slower, with less consistent diameter differences and narrower separation of subtype median velocities (Extended Data Fig. 6a-f). D-to-C capillaries also had lower estimated shear stress. Across capillaries, longer baseline segments and lower baseline velocities were associated with higher stall risk, whereas diameter showed no clear association (Supplementary Table 3).

We next tested whether the higher stall risk of D-to-C capillaries depended on network position. D-to-C capillaries generally showed elevated stall risk across the arterial-to-venous axis (Fig. 6o). Stall risk also tended to increase with branch order from the nearest penetrating vessel. D-to-C capillaries had the highest mean stall risk in each displayed branch-order group, followed by C-to-C capillaries (Fig. 6p, compare Fig. 5i,j).

Finally, we asked whether the measured structural, hemodynamic and network properties explained the higher stall risk of D-to-C capillaries. After adjustment for baseline diameter, segment length, velocity, arterial and venous branch orders, tissue depth and proximity to the imaging boundary, D-to-C capillaries retained higher stall odds than the other subtypes combined (Fig. 6q; Supplementary Table 4). Thus, the association between D-to-C topology and stalling was not fully explained by the measured differences in morphology, hemodynamics, or network position.

### Acute capillary stalls predict long-term flow loss

We next asked whether capillary stalls at 1 h predicted the longer-term capillary perfusion. Tracking baseline capillaries at 1h and day 7 post-stroke revealed four perfusion fates: perfused at both time points, recovered stall, delayed stall, and persistent stall (Fig. 7a-g). Among 2,375 capillaries with resolved fate, 64.8% remained perfused at both time points (Fig. 7h). Of 307 stalled capillaries, 84 recovered by day 7, whereas 223 remained stalled at day 7. An additional 529 capillaries developed delayed stalls, increasing the overall stalled fraction to 31.7% at day7 (Fig. 7i, Extended Data Fig. 7a).

**Figure 7:**
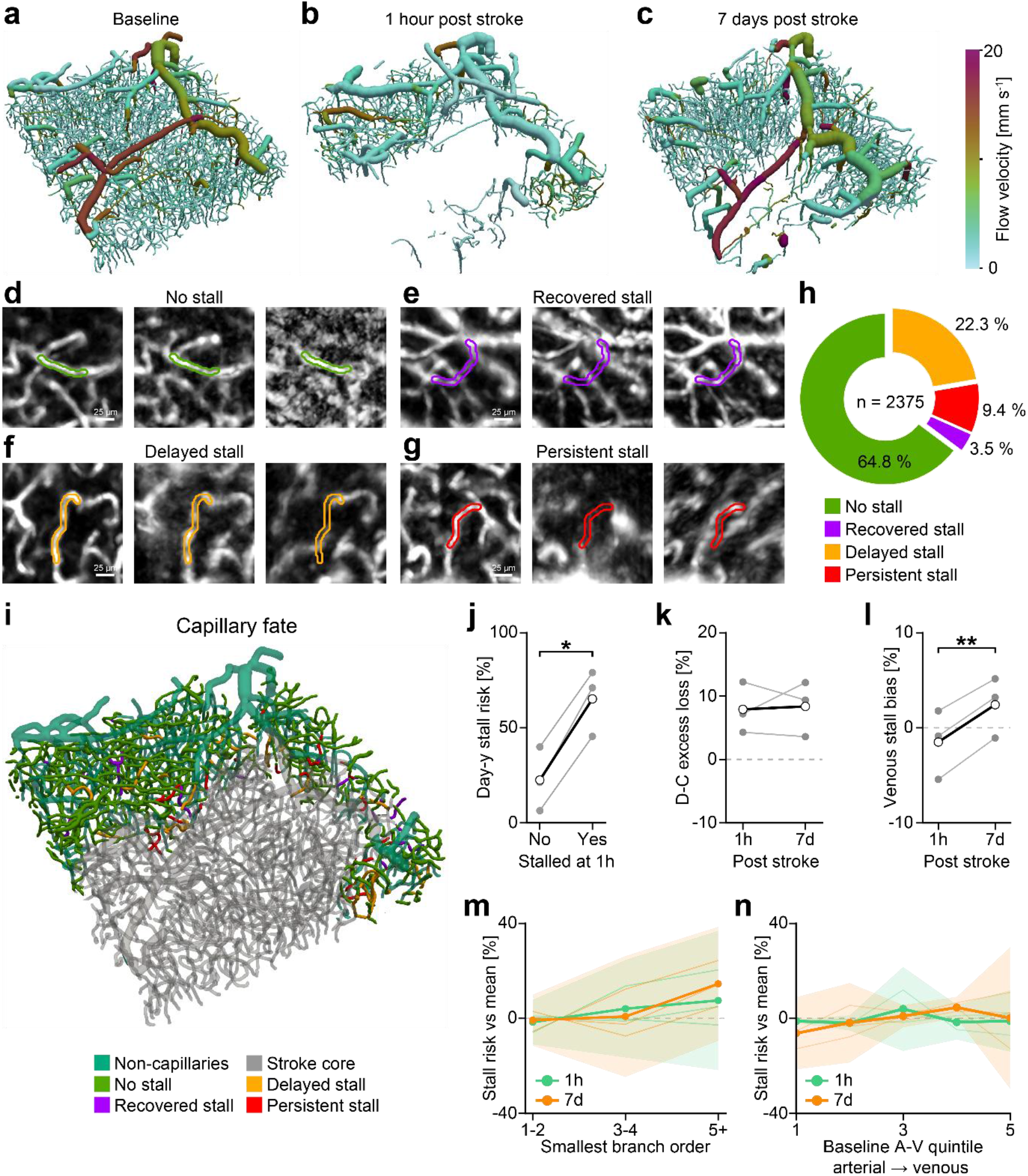
Acute capillary stalls predict later perfusion loss. (a-c) The same cortical network at baseline, 1 h and 7 days after stroke, colored by flow velocity. (d-g) Examples of capillaries with no observed stall, recovered stall, delayed stall and persistent stall, respectively, from baseline through 1 h to day 7. Scale bars, 25 µm. (h) Pooled fates among 2,375 baseline capillaries: 1,53S perfused at both imaging time points, 84 recovered, 52S delayed and 223 persistently stalled. (i) Longitudinal fates of capillaries. (j) Day-7 stall risk after acute perfusion or an acute stall: equal-animal means, 22.5% and c5.2%; difference, 42.7 percentage points (S5% CI: 27.4-58.0). (k) D-C excess risk over other classified baseline subtypes: 7.S percentage points at 1 h (S5% CI: -2.1 to 17.S) and 8.4 at day 7 (−2.5 to 1S.2) with paired change of 0.5 points (−S.5 to 10.4). (l) Venous-minus-arterial rank-half risk difference within baseline venous drainage territories: -1.5 points at 1 h and +2.4 at day 7 with paired increase of 3.S points (S5% CI: 2.7-5.2). (m,n) Stall risk relative to each animal’s overall risk by nearest penetrating-vessel order and baseline arterial-venous quintile. Thin lines show mice, bold curves and shading show equal-animal means and S5% CIs. In j-l, gray connected points show mice and black their mean. N=3 mice; two-sided paired t-tests. All quantitative panels use capillaries outside the 1 h OCTA-defined core and resolved fate at both follow-ups. *: P < 0.05; **: P < 0.01.

Acute stalling outside the acute OCTA-defined core consistently predicted stalls at day 7 (Fig. 7j). Across mice, 65.2% of capillaries stalled at 1 h were also stalled on day 7, compared with 22.5% of those perfused at 1 h.

We also asked whether the D-to-C vulnerability identified acutely persisted at day 7. Among 1,567 capillaries with a classified baseline subtype, D-to-C capillaries had higher stall risk than the other subtypes combined in each mouse, at both 1 hour and day 7 (Fig. 7k).

Finally, we tested whether the relationship between network-position and stall risk changed over time. Within venous drainage territories, stall risk shifted toward the venous side and the venous-to-arterial risk difference increased at day 7 (Fig. 7l). Capillaries at least five branch orders from the nearest penetrating vessel also had above-average stall risk at day 7 (Fig. 7m) and stall risk across arterial-venous position quintiles is shown for both visits in Fig. 7n.

## Discussion

Understanding how microvascular networks fail during stroke requires imaging tools that capture both capillary-scale flow and network-wide architecture, an unmet challenge *in vivo*. OMNIMap addresses this gap by combining extended-focus OCM with deep-learning segmentation and vessel graph analysis, enabling *in vivo* mapping of flow and connectivity across thousands of cortical vessels. By capturing volumetric data at capillary resolution and integrating DOCT to measure blood flow velocities and directions, we provide a holistic view of the cortical microvasculature, encompassing arteries, arterioles, capillaries, venules and veins. This approach overcomes longstanding limitations of manual vessel tracing and imaging techniques restricted to small fields-of-view, thus enabling quantitative assessments of vessel morphology and flow metrics on a large scale *in vivo*.

OMNIMap enables large-scale mapping of flow-resolved capillary connectivity, revealing several key features of network behavior after stroke. First, roughly one-sixth of capillaries stall after stroke, consistent with prior reports in the penumbra^15^. Although stalls are more frequent near venules, this reflects their higher density rather than intrinsic vulnerability. Capillaries separated from penetrating vessels by more branches were more likely to stall, even at similar relative arterial–venous positions. Beyond overall stalling patterns, we identify and quantify four topological capillary subtypes (D-to-D, D-to-C, C-to-D, C-to-C) that intermix throughout the network. Post-stroke, their distribution shifts only modestly. Despite this broadly stable composition, D-to-C capillaries were more likely to stall, even after accounting for their baseline morphology, velocity and network position and become a potential target for future therapeutic interventions.

Capturing the distribution of capillary flow subtypes *in vivo* has been challenging due to the need for large-scale imaging at capillary resolution, with artery-vein labeling and flow direction sensing. Prior studies have relied on *in silico* modeling^40^. Our in vivo data extend these simulations by measuring which subtypes are most likely to stall after arterial occlusion. Simulations predict that stalls in C-to-D capillaries produce the most pronounced downstream effects, whereas D-to-C stalls have more localized impact^40^. Importantly, we find D-to-C capillaries are most likely to stall, with mean stall risk exceeding that of the other subtypes. D-to-C capillaries are also by far the most prevalent in our dataset. Thus, both the likelihood of a connection failing and the consequences of its failure matter for the network. Incorporating measured subtype frequencies and stall risks into these models could improve predictions of how multiple capillary failures affect the surrounding network. Recent in vivo measurements also revealed predominantly diverging junctions near penetrating arterioles, converging junctions near venules, and mixed junction patterns within the intervening capillary network^21^. Our analysis extends this junction-level description by relating the flow patterns at both ends of a capillary to its susceptibility to stalling after stroke.

D-to-C capillaries remained at heightened risk after joint adjustment. Alternative connections at both ends could allow surrounding flow to continue during a stall, limiting pressure changes that might dislodge obstructing RBCs or neutrophils^15,35^. Pressure differences and cell displacement were not measured. D-to-C capillaries had lower derived shear-stress estimates but no clear pressure-drop difference. These estimates are derived from the measured vessel properties and do not independently establish this mechanism. Residual risk could also reflect unmeasured biological properties.

Pericytes could be potential contributors to subtype-specific stall risk. Previous studies demonstrated that pericyte dysfunction can impair capillary reperfusion and lead to capillary stalls in stroke and other neurological disorders^8,13,30,37^. Future studies investigating whether pericyte coverage, soma position or function is different in these capillary subtypes would be interesting for understanding if this vulnerability is driven by pericytes.

Capillary velocity responses varied after stroke, with both decreases and increases, consistent with flow redistribution after arterial occlusion. Initially slow capillaries preferentially increased their velocity beyond the variation observed in independent baseline measurements. However, neither stall adjacency nor route analyses identified a consistent bypass signature. Changes in upstream supply, local vascular tone and the balance of neighboring flows could all contribute. Collateral circulation may also shape the responses^41–43^, but its contribution was not investigated in this study.

Acute stalls predicted day-7 flow loss despite recovery and delayed stalling within the same networks, complementing longitudinal angiography^44^ and repeated stall measurements^45^. In contrast, the higher day-7 risk of D-to-C capillaries remained uncertain, and the shift toward the venous side was local rather than network-wide. Importantly, persistent OCTA non-detection does not establish anatomical pruning or uninterrupted arrest. More frequent imaging could distinguish sustained failure from repeated loss and recovery, while vessel-wall labeling could determine whether non-perfused capillaries are structurally retained or pruned^46^ .

While yielding novel insights into capillary flow topology, our method is not free of limitations. OMNIMap prioritizes volumetric coverage over temporal resolution, limiting the detection of rapid vasodynamics such as heart-rate oscillations or transient stalls. Future implementations using specialized acquisition modes or single-vessel scanning could address this, as demonstrated in previous xfOCM studies of hemodynamic changes^47^ and capillary stalls^16^. Flow velocities are also derived indirectly via Doppler shift and variance, reflecting a combination of RBC and plasma motion, which can reduce accuracy for very slow flows (<0.5 mm·s^-1^) or small changes^28^. Finally, fit-derived diameter measurements support population-level comparisons, although subtle changes in individual capillaries remain difficult to resolve. Depth, signal and cropping controls quantify measurement and field-of-view limitations.

While prior *in vivo* studies have explored the effects of single capillary occlusions^48^, none have considered the four flow topology subtypes identified here at scale. Building on prior simulations^40^, and our observation of stable subtype distributions post-stroke, future work should selectively occlude capillaries of each subtype while tracking downstream and upstream flow responses to clarify their local impact. This could distinguish susceptibility to stalling from the consequences of losing a particular connection. Combining subtype mapping with oxygen measurements would test whether the frequent stalling of the longer D-to-C capillaries creates distinct patterns of local oxygen loss. When a side branch loses flow, adjacent surviving segments can form a longer uninterrupted flow path without anatomical fusion or elongation. Whether this increases the later vulnerability of the surviving path to stalling remains an important question. Finally, more frequent longitudinal imaging and follow-up beyond day 7 may uncover how flow topology evolves during recovery and network remodeling.

OMNIMap presents a compelling opportunity to evaluate emerging stroke therapies targeting microvascular dysfunction. Capillary stalling, a key driver of the no-reflow phenomenon, underscores the need for adjunct treatments beyond thrombolysis. Candidate approaches include inhibiting neutrophil adhesion or, as shown in Alzheimer’s disease^13^ models, using vasodilators to improve perfusion and reduce immune cell stalling. A key question is whether these therapies can effectively reduce stalls, restore flow, and improve later network recovery. Our thrombin model could be combined with tPA treatment to study recanalization. Yet, one limitation of our study is that we cannot rule out the effect of anesthesia. Moreover, further cohorts with increased animal numbers would be required for clinical translation while considering species differences such as in the arterial-venous angioarchitecture. These differences include the reported predominance of penetrating arterioles in human cortex, compared with ascending venules in mouse cortex^49^. OMNIMap is uniquely suited for addressing these questions, complementing high-resolution and molecularly specific 2PM by offering a network-wide perspective on therapeutic efficacy.

Overall, OMNIMap enables functional imaging of the cortical microvasculature, capturing vessel connectivity, flow velocity, and direction, before and after stroke. This allows tracking stroke-induced changes in flow topology at the capillary network level, offering insight into how perfusion is preserved or lost. It also links capillary vulnerability to local topology and connects acute stalls to later perfusion fate. By bridging single-vessel resolution with mesoscale coverage, OMNIMap provides a scalable framework for investigating other small-vessel pathologies, including Alzheimer’s disease, age-related capillary decline, and hypertension.

## Methods

### xfOCM system

The imaging system and its performance have been described previously^28,29^. The current configuration^28^ incorporates an upgraded Exalos superluminescent diode (EBS300040-02, Exalos AG, Switzerland; central wavelength, 1315 nm; 3-dB bandwidth, 120 nm) and 90:10 rather than 50:50 beam splitters to increase illumination and sample-arm signal while maintaining sufficient reference-arm power. The theoretical axial resolution derived from the source spectrum is approximately 4 µm in tissue, assuming a refractive index of 1.33. Optical power at the sample was 15 mW, measured after the objective. Imaging was performed with a 10× infrared-optimized objective (NA 0.3), providing an experimentally measured lateral resolution of approximately 3 µm across the 360 µm depth-of-focus.

### xfOCM imaging

The 1 h post stroke (acute) analysis included 12 mice (BALB/c, aged 8-12 weeks, 6 female, 6 male) with valid baseline and post-stroke registration. One of these mice was excluded from velocity and velocity-derived analyses because of spectral-background contamination, leaving 11 velocity-eligible animals. Five mice underwent longitudinal imaging. Of these, three had accepted baseline, acute and day-7 comparisons for quantitative vessel tracking. Acquisition availability and analysis-specific exclusions are listed in Supplementary Table 1. Two additional mice we imaged but excluded from all results due to the stroke core covering the 1 h post stroke OMNIMap acquisition completely, leaving no vessels to be imaged. All animal experiments were performed in accordance with the Swiss Federal Act on Animal Protection and approved by the Cantonal Veterinary Office Zurich (animal licenses ZH030_2023, ZH165/19).

After cranial window implantation, mice were allowed to recover for two weeks before xfOCM measurements. During imaging, animals were head-fixed under anesthesia (details below). To enhance and homogenize the Doppler OCT signal in capillaries, a 100 µL bolus of 20% Intralipid (MedChem Express) was injected intravenously five minutes before data acquisition. A 1 mm × 1 mm field of view (FOV) was selected to capture downstream branches of the middle cerebral artery (MCA), downstream branches of the venous tree, and the intervening capillary bed. The final position of the motorized stages was recorded to enable repeated imaging at the same site. Additionally, a reference widefield image of the cranial window was acquired under green LED illumination.

xfOCM data collection proceeded in two steps, as described previously^28^: three repeated volumes (two for longitudinal imaging) were acquired with the Doppler OCT protocol (250 repeated A-scans per position at 22,978 Hz), followed by six (or eight) repeated OCTA volumes (four or eight repeated B-scans at approximately 76 Hz; A-scan rate, 45,956 Hz) at the same location. The reconstructed FOV spanned 1000 × 1000 × 360 µm³, with OCTA voxel spacing of approximately 1.95 µm laterally and 2.02 µm axially. Repeated OCTA volumes were averaged over several minutes. The same imaging protocol was repeated post-stroke. To ensure consistent positioning, the motorized stages were first returned to the recorded baseline coordinates. Live widefield camera images were then correlated with the baseline reference image and adjusted laterally until no pixel offset remained. Finally, axial alignment was refined by comparing real-time structural OCT scans with baseline reconstructions, focusing on matching the glass coverslip’s axial position.

### xfOCM reconstruction and registration

OCTA and DOCT data were reconstructed, registered, segmented, skeletonized, and processed to extract blood flow velocity vectors according to a previously described pipeline^28^. For brevity, only key definitions, modifications, and additions are presented hereafter. The underlying Doppler velocity method was previously characterized using flow phantoms and sequential in vivo comparison with two-photon microscopy^28^.

### Doppler reconstruction

For Doppler reconstruction, the complex OCM time series was high-pass filtered using a 250th-order equiripple filter with 270-Hz stopband and 400-Hz passband edges and 40-dB stopband attenuation. The signal was divided into non-overlapping 250-sample windows, Hann weighted, zero-padded, and transformed using a 4,096-point FFT to obtain the Doppler magnitude spectrum. Following spectral-background subtraction, each spectrum was fitted with a constrained least-squares single-Gaussian model. The fitted spectral center and width were converted using the system-specific optical model^50^ into axial v_z_ and transverse v_t_ velocity components, respectively, and total velocity was calculated as 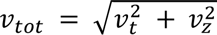 . The Doppler-derived angiogram-like volume (maxVol) was defined voxelwise as the greater peak magnitude in the positive- or negative-frequency half of the spectrum and used to register Doppler to OCTA volumes.

### Point-based velocity estimation

Each acquisition contributed two or three reconstructed Doppler frames to vessel velocity estimation. Frames were processed independently and registered to the first frame using the Doppler-derived angiogram-like volume (maxVol) as the anatomical reference. Native velocity measurements were treated as individual points, each retaining its measured value and fit quality. Their coordinates were transformed into alignment with OCTA without interpolating the velocity values onto a new image grid. Samples were then pooled across frames before vessel-level estimation. Capillary measurements required native-fit R² ≥ 0.5 and at least ten eligible sample memberships per segment before selecting the highest-R² 30% within each cross-section. Diameter-dependent within-section aggregation was followed by the median across sections. Pial and penetrating vessels used a separate finite-fit estimator without the capillary R² selection. The two estimators and velocity-floor correction are specified in Supplementary Note 1. Independent-frame controls for velocity increases are described in Supplementary Note 2 and spatial and alternative-route analyses in Supplementary Note 3.

### Volumetric registration

Between-visit registration aligned post-stroke OCTA volumes to baseline using initial depth alignment, constrained two-dimensional free-form registration of an OCTA maximum-intensity projection, axial refinement by translational image registration and multiscale three-dimensional deformable registration. We open-source our GPU-accelerated registration library that was developed for this project https://github.com/lukasglandorf/accel-deform-registration. The composed transformations aligned vessel coordinates for correspondence (Supplementary Fig. 12). Diameter, velocity and native graph measurements were retained in their original measurement reference frame. Registration quality was visually assessed to confirm if single-capillary comparisons remained valid in each dataset. Registration overlap and depth-dependent segmented-vessel support determined the tissue comparable between visits. Acute comparisons used pairwise support, whereas comparisons involving both acute and day-7 outcomes used joint support across all three visits. Registration stages, quality checks and the limited residual refinement are described in Supplementary Note 4. Depth-dependent signal and measurement-availability controls are described in Supplementary Note 5.

### Blood vessel segmentation

We employed an existing 3D U-Net–based approach (Supplementary Fig. 1) for volumetric OCTA segmentation, which was initially pre-trained on synthetic OCTA volumes ^51^ generated from real vascular graphs^52^ and subsequently fine-tuned with manually labeled xfOCM data. Unlike our earlier work^28^, we used all six labeled samples in a leave-one-out fine-tuning scheme: each model was trained on five samples and evaluated on the held-out sample. The mean held-out Dice score was 81.8% (Extended Data Fig. 1a). The six models were subsequently ensembled during inference.

The final trained model is publicly available on https://huggingface.co/bwittmann/octa-unetplusplus, with the inference pipeline hosted on https://github.com/bwittmann/syn-cerebral-octa-seg/tree/octa-unetplusplus.

### Vessel skeletonization and graphing

After segmentation and morphological cleaning, each volumetric blood vessel segmentation map was skeletonized using a specialized algorithm implemented in Voreen^53^. A corresponding vessel graph was then constructed, with branch points defined as nodes and the vessel segments between them as edges. Each edge stores specific characteristics of its vessel segment, such as centerline coordinates, diameter, and blood flow velocity, while the overall graph encodes the connectivity between vessels.

### Vessel diameter estimation

A segmentation-derived distance-field estimate for each vessel segment was used to define the local sampling window. At each interior point of the smoothed centerline, the native OCTA volume was sampled in a plane perpendicular to the local vessel direction. A transverse intensity profile was then fitted with a Gaussian-plus-background model:

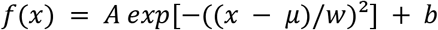

The fitted width *w* was averaged along the segment using only fits with R² > 0.8 and converted from pixels to micrometers to obtain the reported diameter. Capillary analyses used accepted direct OCTA fits. Unresolved values remained missing. Segmentation-derived estimates were retained as a fallback for larger pial and penetrating vessels. Paired changes required compatible measurement sources at both visits. Fit-coverage criteria and the split-half precision analysis are described in Supplementary Note 6. The derived shear-stress and pressure-drop estimates are defined in Supplementary Note 7.

### Graph representation of cortical microvascular network (MVN)

The MVN is treated as a directed graph T = (V, A) consisting of nodes *u* ∈ V and segments *a* ∈ A. Each segment *a* = *uv* is simplified as a straight, cylindrical tube with geometric locations *x_u_*, *x_v_* and given radius *r_a_* and (centerline) velocity *v_a_*. We assume blood to be an incompressible Newtonian fluid and flow to be laminar. The volumetric flow through a segment *a* follows with

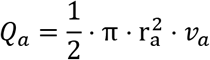

These assumptions for blood flow are shown to accurately describe the scaling relations of vessels^54,55^ and can be used to analyze flow phenomena inside the MVN^56^.

To make the problem of vessel identification and network optimization tractable, we assume vessels in the capillary bed to be correct and only allow changes to the topology (edges) of the pial vessels.

### DA / AV identification

DA and AV are identified by heuristically testing directed paths between different nodes. Only paths with either their start node *u* or end node *v* close to the pial surface are considered suitable candidates. Each candidate must have a penetration depth Δz of at least half the total depth of the MVN and its orientation must be aligned with the z-axis. The orientation is measured by the acute angle α spanned between the vector and the z-axis. Only paths with |α| < 30° are considered aligned. Lastly, a path must be straight with a predominant direction, which is checked by casting an oriented bounding box around the path and measuring the box size. The height of the box must be at least 2 times its length and width.

### Pial network optimization

Before the pial network is optimized, we check and resolve obvious artifacts from the initial segmentation, e.g., self-loops and unphysical radii between bigger vessels. Afterwards, all pial vessels, which directly connect to a DA or AV are included in the vector of fixed segments ̄A. To fix wrong flow direction, we minimize the balance of mass flow at each node *v*, which is defined by

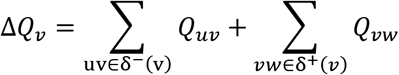

To differentiate between incoming and outgoing edges at node *v* we use:

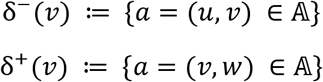

We introduce the decision variables *z_a_* for all segment *a* ∈ A to decide whether flow in *a* is part of the network (*z_a_* = 1), needs to be reversed (*z_a_* = −1) or deleted (*z_a_* = 0). The vector of optimization variables then becomes *y* = (*Q*, Δ*Q*, *z*) and our (discrete) optimization problem is defined with

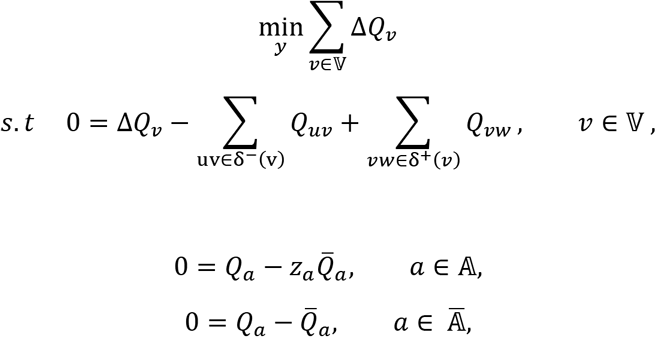

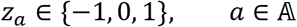

The first constraint defines the balance of mass flow, and the second constraint changes the flow according to the decision value *z_a_*. The third constraint fixes the flow of pial vessels, which are directly connected to a DA or AV.

### Pial vessel labeling

We label the (optimized) pial vessels by analyzing each path between two boundary nodes. A node is considered at the boundary of the pial network if it is either connected to a DA or AV or has only one edge. Along a path, we check each branching node to see if it is converging or diverging and determine if the path is primarily converging or diverging. Lastly, after traversing all paths, each edge is labeled according to the average of all individual paths containing it. In this way, the influence of remaining errors on the final labeling can be better controlled.

The code for the pial network optimization and artery-vein labeling is available as supplementary data.

### Distances between penetrating vessels

Descending arterioles or ascending venules that are directly connected in the network graph are grouped into a local penetrating vessel tree. Each tree’s “starting point” is defined at the node where it connects to the pial vasculature (Fig. 2g). To quantify spacing between penetrating vessels of the same or different types, we measure the lateral distance from this starting point to the closest starting point in another local vessel tree of the designated type (descending arteriole or ascending venule). Repeating this calculation for all trees produces a distribution of inter-tree distances, illustrating how penetrating vessels are arranged across the cortical surface (Fig. 2i).

### Blood vessel segment matching between baseline and stroke

To compare individual vessel segments between baseline and post-stroke volumes, it is necessary to find the correct correspondence. We compared centerline geometry in the registered coordinate system and retained accepted one-to-one, split and merge relationships. Each relation retained explicit baseline and follow-up identities. Follow-up measurements were combined once per baseline segment using a weighted geometric mean, with weights proportional to match overlap score multiplied by follow-up segment length. Velocity and diameter comparisons required eligible native measurements at both visits and the same final vessel class. Correspondence failure alone did not define a stall. Matching thresholds, aggregation rules and the restricted one-to-one analyses are specified in Supplementary Note 8.

### Capillary branching order

Each capillary segment’s arterial branch order (BO) is determined by finding the shortest path from either of its two nodes to any node within the arterial tree (pial artery or descending arteriole). Conversely, the venous branch order is computed by identifying the shortest path to a node in the venous tree (pial vein or ascending venule). Thus, each capillary segment is assigned both an arterial and a venous BO. If no path to either tree exists, the capillary is excluded from branch-order analyses. The relative branch order (rBO) provides a relative measure of a capillary’s position between these two trees:

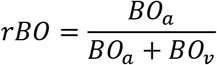

Where BO_a_ is the arterial and BO_v_ the venous branch order. Nearest-penetrating vessel order was defined as min(BO_a_, BO_v_). Physical-path alternatives used cumulative vessel length to each penetrating vessel (Supplementary Fig. 13). Total branch order was computed as tBO = BO_a_ + BO_v_. Topology was calculated on the complete native graph before selecting vessels for measurement or fate analyses (Supplementary Note 9). Arterial-venous capillary counts and shares are described in Supplementary Note 10. Finite-volume sensitivity was assessed using controlled crops of reference networks, including joint variation of lateral field width and imaging depth in surface-anchored crops (Supplementary Note 11).

### Capillary stall definition

A stall was defined as loss of detectable OCTA perfusion along a baseline capillary within tissue assessable at both visits. At least 10 µm of comparable centerline was required. Less than 40% of this length lacking local follow-up vessel evidence indicated present perfusion. 40% to less than 60% remained uncertain. At least 60% was a candidate loss, classified as a stall only after passing residual-signal and shadowing checks. Exact spatial and signal thresholds are given in Supplementary Note 12. Neither failed matching nor a Doppler-velocity cutoff defined a stall. Repeated OCTA acquisitions provided a visit-level perfusion state, without establishing the duration of a flow interruption or anatomical vessel loss.

### Capillary topology subtype

We identify capillary flow topology by classifying their input and output nodes as convergent or divergent, considering all three connected capillaries. If a capillary is not connected to two other capillaries at both nodes, it is not included. Subtype-specific stall analyses required a classified baseline subtype and assessable follow-up fate after stroke. Classification at both visits was required only for analyses of subtype transitions (Supplementary Fig. 14). These eligibility requirements and the vessel-class stabilization across time points are described in Supplementary Note 9.

### Changes in detectable capillary perfusion

Total detectable capillary length was measured independently at each visit within comparable tissue, without requiring individual vessel matches. This length change was kept separate from individual-vessel fate and the net loss of detectable capillaries, estimated by subtracting reverse from forward non-detection fractions within each animal. This comparison accounts for detectable vessels lost and gained between visits without changing individual stall classifications (Supplementary Note 12).

### Longitudinal vessel fate

Longitudinal analysis followed the same baseline capillaries at approximately 1 h and 7 days after stroke in three mice. The primary population used an acute OCTA-defined non-core (penumbra) region fixed for the later visit and required at least 80% of baseline centerline length within tissue assessable at all three visits. Vessels with resolved states at both follow-ups were classified as present at both visits, recovered by day 7, loss at both sampled visits, or loss first detected at day 7. Uncertain outcomes were evaluated separately (Supplementary Fig. 15). Baseline-subtype comparisons imposed an additional subtype-availability requirement. The regional definition, fate populations and LSCI-based core sensitivity are described in Supplementary Note 13.

### Animal anesthesia

For head-post attachment, anesthesia was induced with 4% isoflurane in oxygen at approximately 800 mL/min and maintained at 1.5-2.5% at approximately 300 mL/min. For cranial window implantation and stroke induction, mice first received local anesthetics at the incision site (1 mg/mL lidocaine, 5 mg/mL bupivacaine) and were anesthetized via subcutaneous injection of fentanyl (0.05 mg/kg, Sintenyl, Sintetica), midazolam (5 mg/kg, Dormicum, Roche), and medetomidine (0.5 mg/kg, Domitor, Orion Pharma). For xf-OCM imaging, mice received the same subcutaneous anesthetic regimen. During injectable anesthesia, the animals received a mixture of oxygen and air (30%/70%) at approximately 300 mL/min via a facemask and body temperature was kept at 37 °C using a homeothermic blanket (Harvard Apparatus). Mice were placed in a stereotaxic frame, and their eyes were protected with vitamin A ointment (Bausch C Lomb).

### Imaging head-post implantation

A bonding agent (Gluma Comfort Bond, Heraeus Kulzer) was applied to the cleaned skull and polymerized using a handheld blue light source (600 mW/cm^2^, Demetron LC). A custom aluminum head post was then connected to the skull with dental cement (EvoFlow, Ivoclar Vivadent AG) for stable head fixation during imaging. The incision was treated with antibiotic ointment (Neomycin, Cicatrex, Janssen-Cilag AG) and sealed with acrylic glue (Histoacryl, B. Braun). After surgery, animals were kept warm and analgesics were administered subcutaneously (buprenorphine 0.1 mg/kg and carprofen 10 mg/kg, Sintetica).

### Cranial window implantation

A 4 × 4 mm craniotomy was performed above the left somatosensory cortex (centered at -3 mm from Bregma and 3.5-4 mm lateral) using a dental drill (Bien-Air). The dura mater was exposed and covered with a 3 × 3 mm glass coverslip (UQG Optics), which was secured to the skull using dental cement.

### Thrombin stroke model

Focal cerebral ischemia was induced in the M2 segment of the middle cerebral artery (MCA). Briefly, mice were placed in a stereotactic frame, and a skin incision was made between the left eye and ear. The temporal muscle was retracted, followed by craniotomy and removal of the dura mater. A glass pipette (calibrated at 1 µl) was inserted into the MCA lumen, and 1 µL of purified human alpha-thrombin (1 IU; HCT-0020, Haematologic Technologies, USA) was injected to form a clot in situ. The pipette was removed 10 minutes after injection. Ischemia was deemed stable if cortical blood flow (CBF) fell to ≤50% of baseline and remained at that level for ≥30 minutes. Animals without stable ischemia were excluded from further experiments.

### Laser speckle contrast imaging

Cortical blood flow was monitored before and during ischemia using a laser speckle imaging system (FLPI, Moor Instruments, UK). The software (MoorFLPI) generates relative perfusion maps in arbitrary units using a 16-color palette, enabling the assessment of real-time changes in cortical perfusion. Acute perfusion maps were normalized to baseline and aligned with the OCTA field (Supplementary Fig. 16). Regions were defined on the full LSCI field before applying the OCTA footprint: core indicated at least 70% reduction, with severe and moderate hypoperfusion bands spanning 45% to less than 70% and 20% to less than 45% reduction, respectively. An OCTA example of severe perfusion loss in the stroke core is shown in Supplementary Fig. 17. Local perfusion change and distance to the core boundary were analyzed separately. These LSCI regions describe perfusion severity. The primary longitudinal OCTA regional definition and processing details are specified in Supplementary Note 13.

### Statistical analysis

Animals were the units of statistical inference. Vessel-level measurements were summarized within each animal before combining animal estimates and pooled vessel distributions were descriptive. Paired positive measurement changes were analyzed on the log-ratio scale, with two-sided animal-level tests and 95% confidence intervals. Whole-distribution comparisons used exact whole-animal baseline/follow-up reversals, distinct from tests of paired summary changes. Adjusted stall associations were estimated using animal-specific logistic models and combined by random-effects meta-analysis with Knapp-Hartung intervals. Baseline-property analyses are specified in Supplementary Note 14. Multiple comparisons used the declared analysis-specific Holm families. Animal-bootstrap and t intervals were identified for each analysis. Missing measurements were not imputed, and all eligible observations were retained for inference irrespective of display limits. Exact estimators, model formulas, comparison families and supporting validation analyses are given in Supplementary Note 15. Figure captions report the corresponding animal and vessel denominators.

### Software

The processing pipeline was mostly implemented using Matlab (R2022b, The MathWorks, USA). Additionally, we used some functions provided by Stefan et al.^57^. OCTA-OCTA and OCTA-DOCT registration was done using SimpleITK (v2.3, simpleITK.org) and Python, with the GPU-accelerated deformable registration library described above for between-time point alignment. The deep learning pipeline is built upon PyTorch (pytorch.org) and MONAI (monai.io). The vessel type classification algorithm is implemented in Julia. Volumetric renders were rendered using Paraview (5.8, Kitware Inc., USA). Plots were created using R (v4.3, r-project.org) and the ggplot2 package (v3.4, ggplot2.tidyverse.org). Statistics were computed using R (v4.3, r-project.org).

## Contributions

L.G., M.E.A. and D.R. conceived the study. L.G. developed the experimental system. L.G. and J.D. carried out animal experiments. L.G., E.J., B.Wi. and D.S. developed the processing pipeline and conducted the data analysis and visualization. J.D., M.E.A., R.L., and B.M. contributed to the interpretation of the results. B.We. and S.W. provided the animal model and guided the animal experiments. D.R, R.L., and M.E.A. were involved in the study design and planning and supervised the work. All authors contributed to writing and revising the manuscript.

---

**Extended Data Figure 1:**
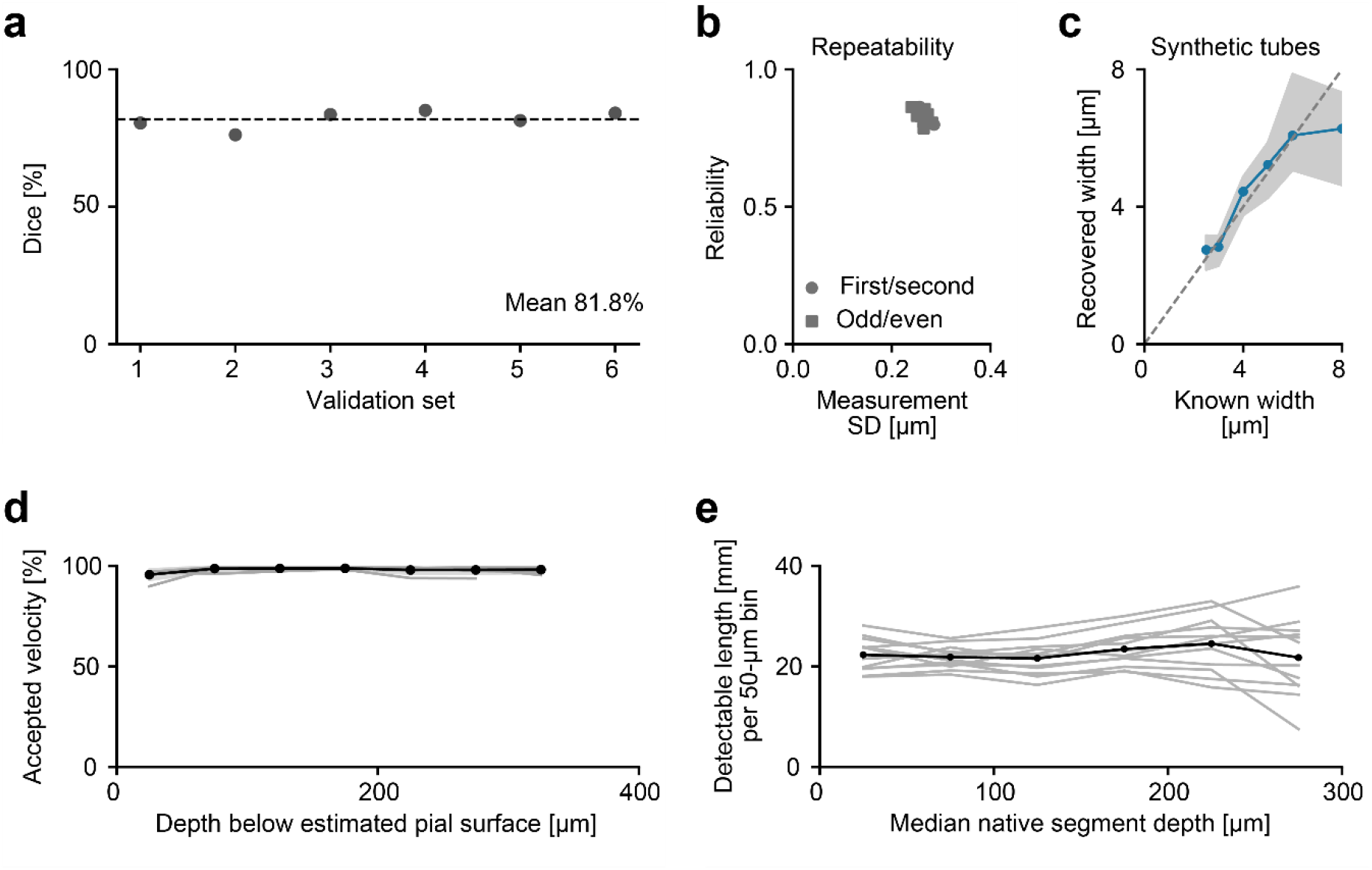
Measurement precision and depth-dependent availability. (a) Dice scores for six validation sets (81.8 ± 3.24%). (b) Repeatability across calibration datasets and visits, using contiguous and odd/even frame halves. Each point is a dataset/visit/split estimate of measurement SD and reliability. Pooled precision is SD 0.2c2 µm, MDCS5 0.72c µm and reliability 0.835. (c) Synthetic-tube recovery: Gaussian tube signals with known widths added to real OCTA backgrounds, varying orientation, signal strength, sampled segment support and junction contamination. Points and shading are medians and quartiles of all finite fits. (d) Velocity availability, relative to estimated pial depth. Thin lines, animals. Black line, mean. Shading, between-animal SD. (e) Detectable baseline length (n=12), assigned to 50-µm bins by median native segment depth.

**Extended Data Figure 2:**
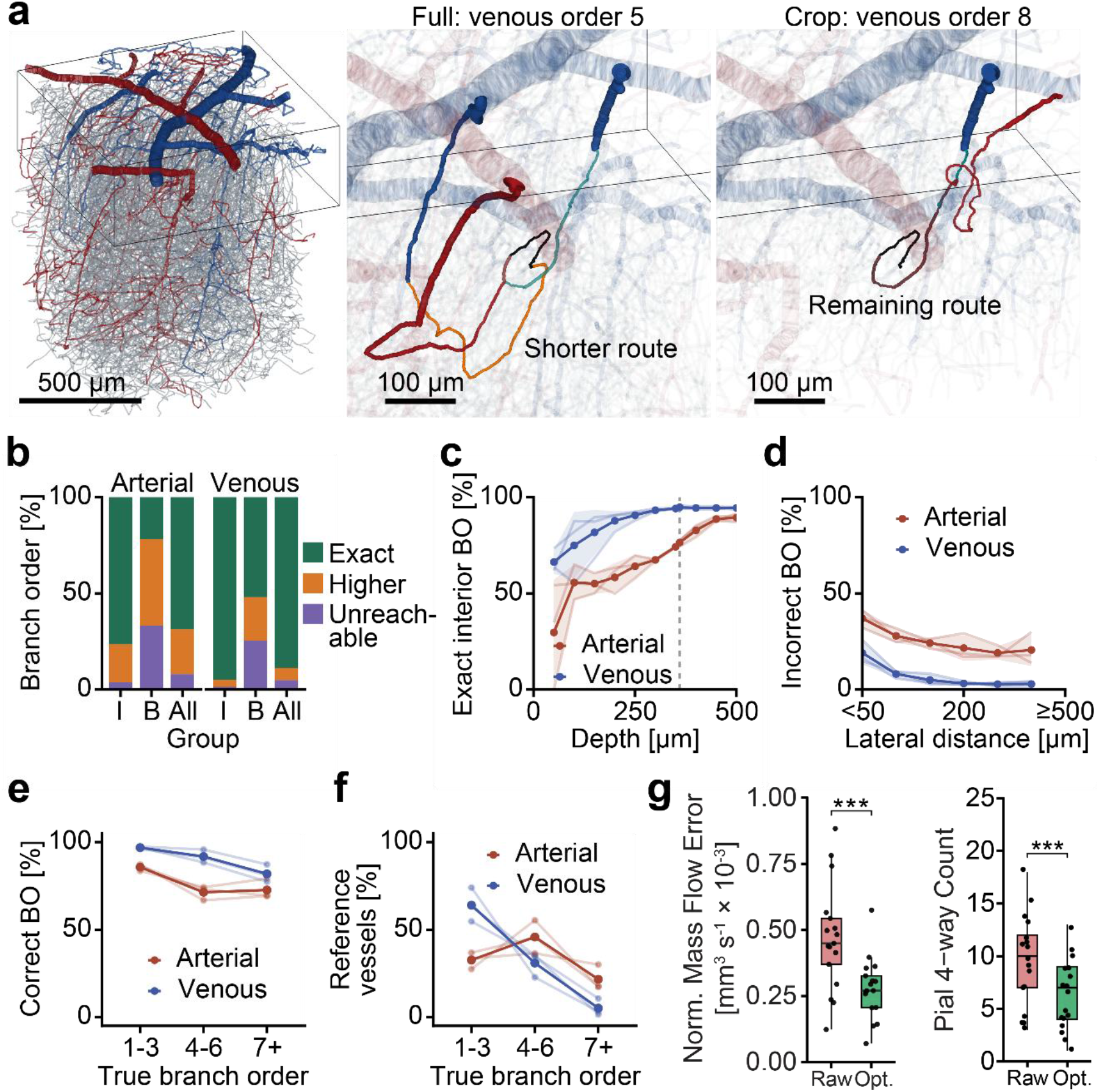
Branch-order recovery in surface-anchored volumes. (a) Reference graph with a pial-surface-anchored 1 × 1 × 0.3c-mm box^1^. Matched close-ups show example of venous order 5 becoming 8 when the shorter connection exits the lower face. (b) Exact, higher and unavailable orders: interior (I), boundary-crossing (B), all vessels. (c) Exact interior recovery versus depth. Dashed line, 3c0 µm, OMNIMap imaging depth. (d) Incorrect or unavailable orders versus nearest lateral-boundary distance. All 42 origins in three networks contribute. Crops are averaged within network, then networks equally. Lines show networks and their mean, shading between-network SD. (e) Accuracy within true-order bins. (f) Frequency of those bins: low values mean uncommon. (g) Normalized mass flow error before (raw) and after (opt.) flow optimization across the pial vasculature for eight mice, imaged at baseline and post-stroke (left, N=1c, p<0.001, paired). Number of four-way connections in the pial vasculature before and after optimization for the same data set (right, N=1c, p<0.001, paired).

**Extended Data Figure 3:**
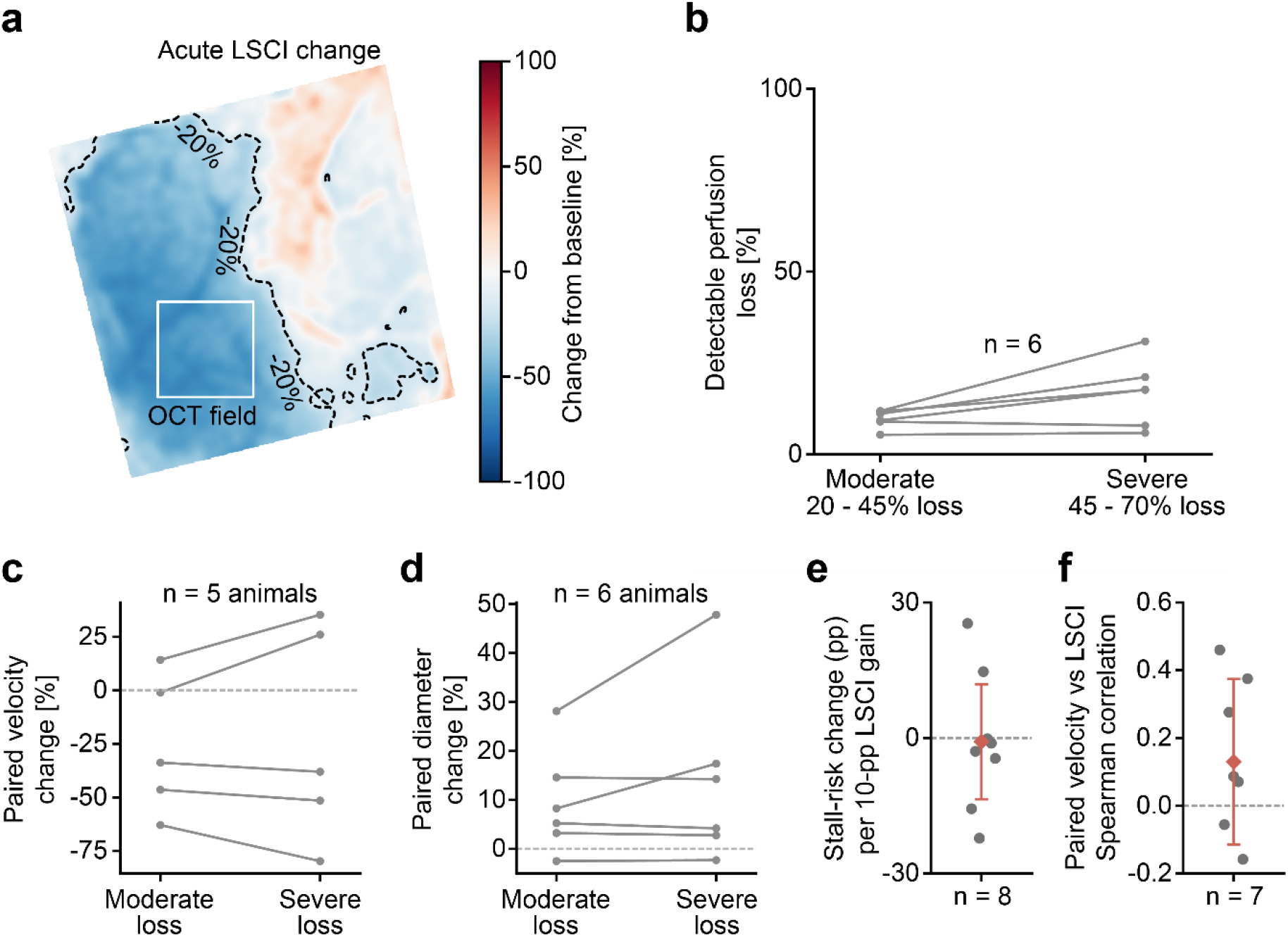
Capillary outcomes versus independently measured LSCI perfusion. (a) Representative LSCI change map with OCTA footprint. This animal has no region meeting the ≥70% reduction core threshold. (b-d) Stall fraction, paired velocity change and paired OCTA diameter change in moderate (20-45%) and severe (45-70%) LSCI-reduction regions. Severe-minus-moderate stall risk is +7.12 percentage points (S5% animal interval -0.51 to +14.75). Velocity and diameter severe/moderate response ratios are 0.S3 (0.c2-1.41) and 1.035 (0.ScS-1.10c). (e) Existing animal-specific change in stall risk per 10 percentage-point improvement in local LSCI (n=8). (f) Within-animal Spearman correlation between paired velocity change and local LSCI (n=7). Diamonds and intervals show equal-animal means and S5% t intervals.

**Extended Data Figure 4:**
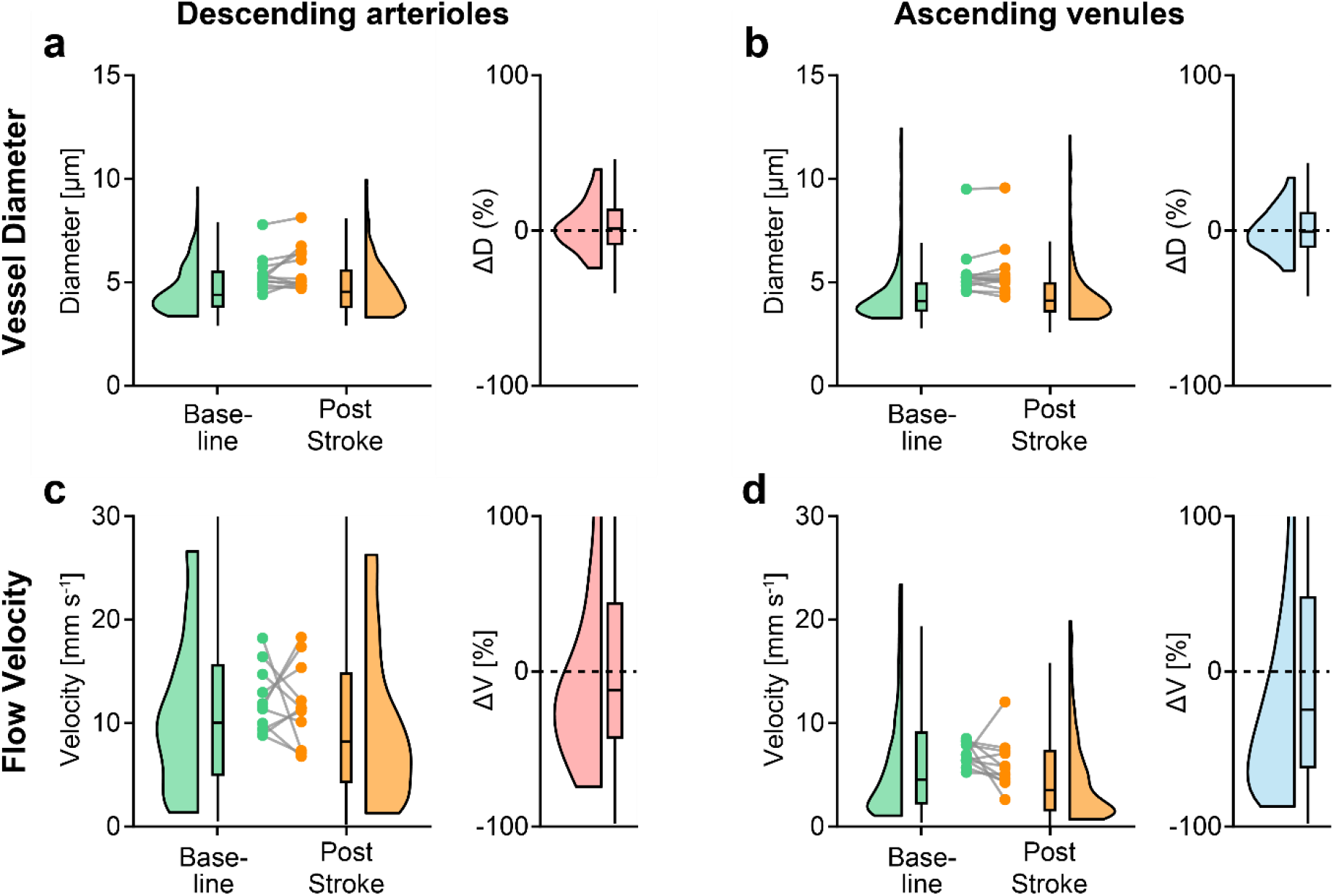
Penetrating-vessel diameter and velocity responses. (a, b) Baseline and 1 h diameter in descending arterioles and ascending venules with relative changes (n=12). (c,d) Corresponding velocity measurements (n=11). Violins show distributions, with density geometry displayed between the 5th and S5th percentiles. Boxes use all paired values and connected points show animal means.

**Extended Data Figure 5:**
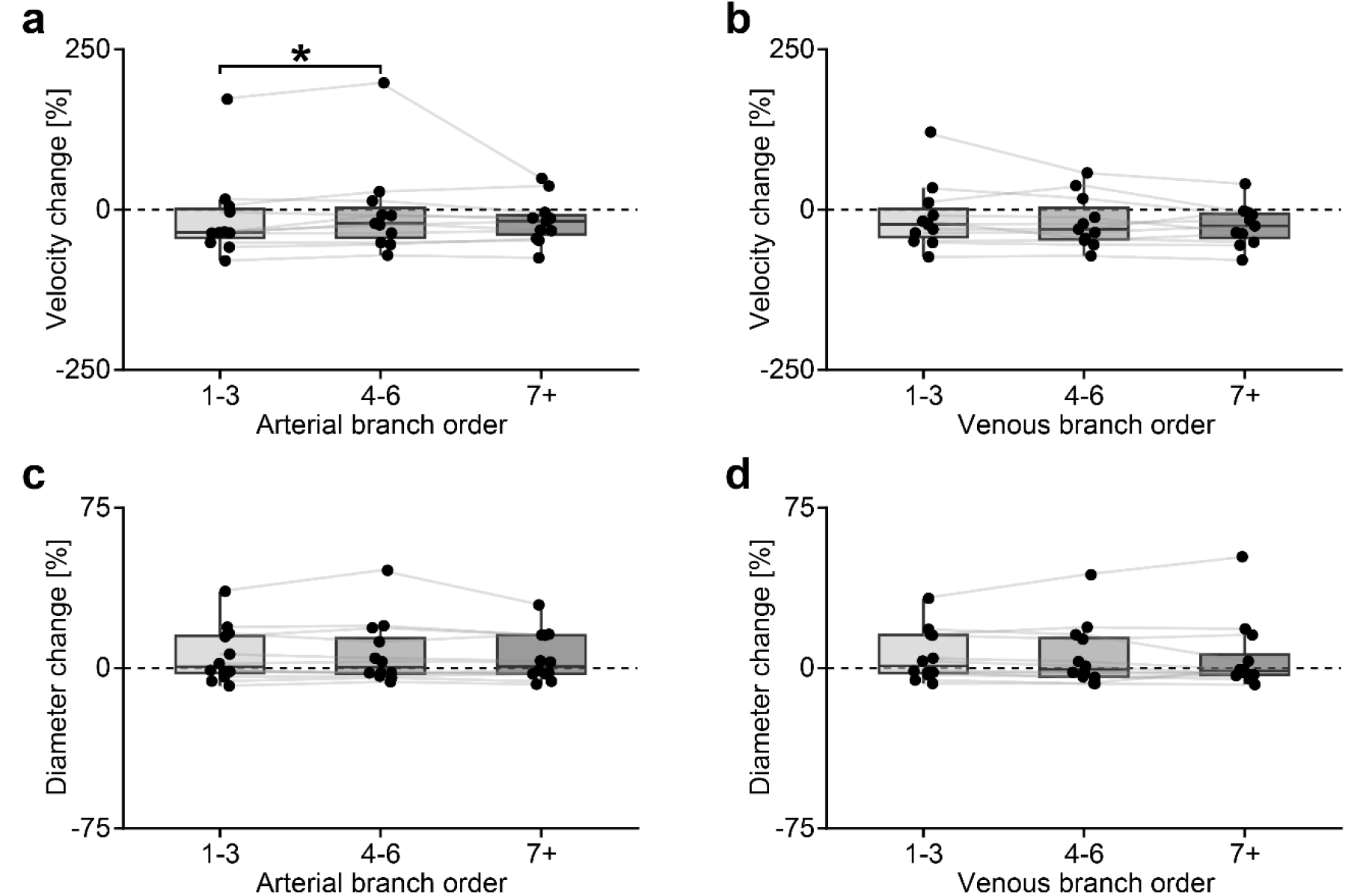
Paired diameter and velocity changes along the network. (a,b) Paired velocity change by arterial and venous branch order. (c,d) Corresponding OCTA diameter changes. Points are animal median paired log changes transformed to percent. Boxes show their median and interquartile range, with 1.5-IǪR whiskers. (n=12 diameter and n=11 velocity). The brackets use Holm-adjusted low-order comparison (*P < 0.05. Arterial velocity P = 0.0334).

**Extended Data Figure 6:**
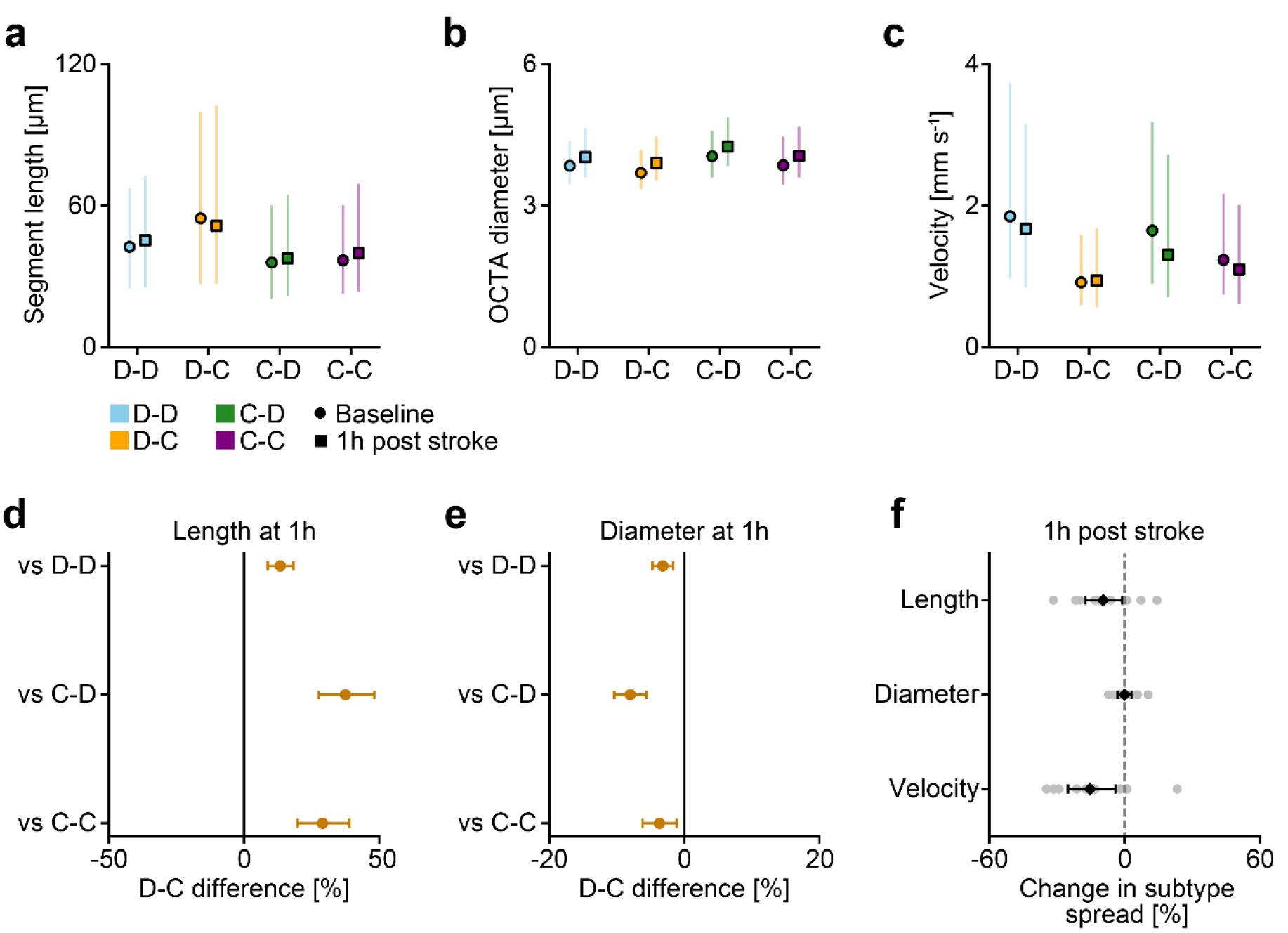
Subtype phenotype at 1h post stroke. (a-c), Length, OCTA diameter and measured velocity for separately classified baseline and 1h post stroke subtypes. Colors identify subtypes. Circles/squares identify time points. Points are equal-animal means of animal medians. Vertical ranges are mean animal quartiles. (n=12 for length and diameter, n=11 for velocity). (d,e) D-to-C contrasts against each other subtype at 1h post stroke, with S5% animal t intervals. (f) Change in the spread among all four subtypes, defined within each animal as the largest divided by the smallest subtype median. Gray points are animals. Diamonds and bars are log-scale means and S5% t intervals, transformed to percent. Spread changes are -S.5% for length (interval -17.4 to -0.S%), -0.04% for diameter (−3.1 to 3.1%) and -15.3% for velocity (−25.2 to -4.0%). Holm-adjusted P across the three metrics: 0.0c7S, 0.S77 and 0.0428.

**Extended Data Figure 7:**
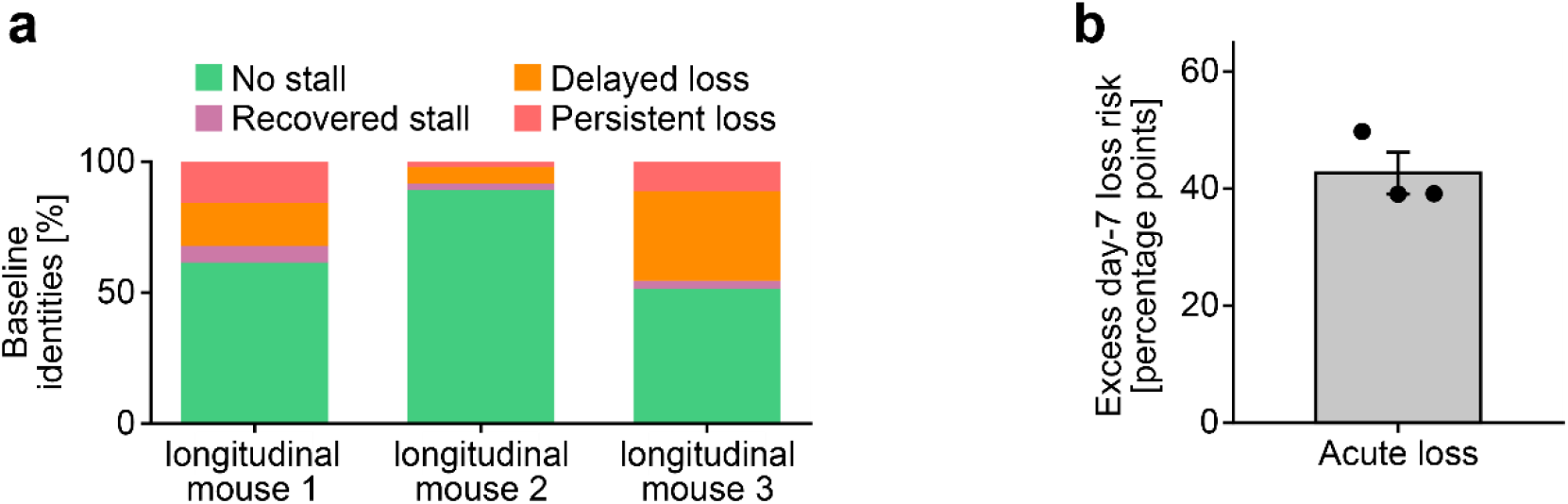
Longitudinal capillary fates across animals. (a) Fate composition among the 2,375 baseline identities with resolved acute and day-7 outcomes in three animals. Present both, recovered detectable perfusion, delayed loss and loss at both visits are distinguished. (b) Mean excess day-7 loss risk following acute loss compared with acute presence. The bar shows the equal-animal mean, the error bar shows SEM and points show the three animal risk differences.

