## Supplementary Material for "In Vivo Network-Level Cerebrovascular Mapping Reveals the Impact of Flow Topology on Capillary Stalls After Stroke"

Supplementary Notes (SN) 1 - 15, Supplementary Figures (SF) 1 - 17, Supplementary Tables 1 - 4

**SN 1: Doppler sampling and vessel-level velocity estimates**

For Doppler reconstruction, the complex OCM time series was high-pass filtered using a 250th-order equiripple filter with 270-Hz stopband and 400-Hz passband edges and 40-dB stopband attenuation. Non-overlapping 250-sample windows were Hann weighted and zero-padded for a 4,096-point Fourier transform. After spectral-background subtraction, the Doppler magnitude spectra were fitted with a Gaussian model. The fitted spectral center and width were used to estimate axial and transverse velocity components using the system-specific optical calibration, and total velocity was calculated as the square root of the sum of their squares.

Each Doppler frame was reconstructed independently. Registration placed each sample in the OCTA reference coordinates while retaining its measured velocity and spectral-fit R² by treating each voxel as a point measurement and not interpolating between points when applying transformations. Samples from repeated frames were collected around each vessel centerline point before estimating vessel velocity. The velocity values themselves were not interpolated into a registered image.

The sampling radius at each centerline point, in graph voxels, was:

$$r = max[round(D / 2), 1]$$

Here, $D$ is the diameter used to assign samples to the graph, also in graph voxels, and round denotes rounding to the nearest integer. Before calculating the radius, the graph diameter was rounded to an integer, even values were increased by one, and the minimum diameter was one voxel. D in the equation denotes this adjusted diameter; the 6-µm aggregation threshold below used the unrounded graph diameter. Sampling regions around neighboring points could overlap. A sample could therefore contribute to more than one cross-section. The reported sample count includes these repeated assignments and does not represent independent biological observations.

For capillaries, eligible samples had finite positive velocity and native-fit R² ≥ 0.50. At least ten eligible sample memberships across the vessel segment were required before quality-based selection. Within each cross-section, eligible samples were ranked by R². The highest 30% were retained, rounding the sample count upward and retaining all ties at the boundary. After this quality selection, the within-section estimate was the median of retained velocities for diameters ≤6 µm. For larger diameters, it was the median of the fastest 30% of retained samples, rounding the count to the nearest integer and retaining at least one sample. Segment velocity was the median across available cross-sections.

Pial arteries, descending arterioles, ascending venules and pial veins used finite native fits without a numerical R² floor and required at least ten eligible sample memberships per segment. They used the same diameter-dependent within-section aggregation followed by the median across sections, without the capillary top-30%-R² selection.

After calculating segment velocity, the correction was applied once:

$$v = \sqrt{max(u^{2} - c₀^{2}, 0)}$$

Here, $u$ is the uncorrected segment velocity and $v$ is the corrected velocity. The constant $c₀ = 0.49$ mm s⁻¹ represents the estimated flow-independent spectral broadening, expressed as an equivalent velocity. This accounts for broadening from the finite acquisition window, Brownian motion and system effects that would otherwise contribute to the velocity estimate.

The broadening estimate was obtained from the narrowest well-resolved Doppler peaks in one baseline dataset from each of the three acquisition series. Fitted spectral widths were pooled across B-scans, retaining fits with R² > 0.90, a spectral peak-to-median magnitude ratio >50, and absolute width <800 Hz. The 10th percentile of these widths was used to estimate the lower limit of spectral broadening. This width was converted to its transverse-velocity equivalent using the same optical calibration as the Doppler measurements, and a common $c₀$ of 0.49 mm·s⁻¹ was used.

A zero produced by the correction was distinct from a missing measurement. Instead, missing or rejected velocities and vessels without detectable perfusion were not assigned zero velocity.

Changes in individual vessels were calculated after combining their matched follow-up measurements (SN 8). Vessel measurements were first summarized within each animal, and animal summaries were then combined (SN 15). Comparing all detectable vessels at two time points does not necessarily compare the same vessels.

**SN 2: Independent-frame controls for velocity increases**

One baseline Doppler frame selected the slowest quarter of measurable capillaries. Other frames estimated same-time point and post-stroke changes. Frame roles were rotated, comparisons were first summarized within animal, and the difference between post-stroke and same-time point effects was evaluated across animals with sufficient independent baseline frames. This same-time point retest addresses selection and measurement noise but it does not provide a sham control over the full between-time point interval. The independent-frame control and paired velocity-change distributions are shown in Supplementary Fig. 4.

**SN 3: Spatial distribution of velocity increases and alternative routes**

Adjacency meant sharing a baseline graph junction with a capillary classified as losing detectable perfusion. Among survivors with accepted velocity at both time points, the probability of increased velocity and the adjusted change in velocity were compared for adjacent and other capillaries. Adjusted comparisons included baseline velocity, segment length, depth and boundary proximity. Distance measured by the number of connecting vessel segments was assessed separately. Directed arterial-to-venous routes were defined from connectivity before inspecting velocities. Unusually fast route fractions used a fixed within-animal baseline mean-plus-three-SD threshold, with time point-specific thresholds and independently reconstructed follow-up routes as separate sensitivities. Exact alternative-route searches around losses were restricted to the documented geometry and measurement support. Animal-specific route coverage and effects were retained. Spatial and route analyses are shown in Supplementary Fig. 5.

**SN 4: Registration and comparable tissue**

Repeated acquisitions were aligned using image-derived transformations, with baseline as the reference for between-time point comparisons. The combined transformation was applied to vessel coordinates to identify matches between time points. Velocity values and spectral-fit quality were retained from the original frame of reference, and diameter was measured in the original OCTA volume. Registration therefore aligned vessel positions without changing their measured velocity or diameter.

After initial depth alignment, a two-dimensional maximum-intensity projection from the upper third of the cropped angiogram supplied the in-plane free-form registration, which was propagated to three dimensions. An axial translation was then refined by image correlation. Slice-wise contrast-limited adaptive histogram equalization preceded four three-dimensional free-form deformation stages. The saved control-grid spacings were 40, 40, 20 and 20 voxels. Stages 1-3 used correlation similarity and stage 4 used local normalized cross-correlation. Smoothness weights were 0.04, 0.04, 0.04 and 0.02, and bending weights were 0.02, 0.01, 0.01 and 0.01, with an exponential Jacobian penalty to penalizes voxel folding. The projection-stage grid spacing was 100 voxels and the configured overlap-region margin was 20 voxels. Transformations were composed in their recorded order, using the inverse image mapping for point coordinates. The settings and transformation sequence for each comparison accompany the code.

Comparisons were restricted to tissue with valid registration and sufficient segmented-vessel signal at both time points. The usable depth was determined by the 15% segmentation criterion described below, independently of vessel fate, velocity or subtype. Acute (1 h) analyses used tissue assessable at baseline and the 1 h time point. Analyses of acute and day-7 outcomes used tissue assessable at all three time points. Longitudinal vessel comparisons required at least 80% of each baseline centerline's physical length within this shared tissue. The minimum 10-µm assessable length used for acute vessel-fate classification is a separate requirement (SN 12).

To determine usable depth, the fraction of segmented pixels and the number of separate segmented regions in each cross-section (minimum two pixels) were summarized in 20-µm slabs and smoothed with a three-bin running median. Each was compared with its own time point's 50-150 µm reference band. A slab passed if both quantities were at least 15% of their reference values, the segmented fraction was ≥0.001, there were at least five segmented regions per layer, and registration was valid over ≥80% of the geometry. Starting at the deepest slab and moving upward, the shared depth limit required three consecutive passing slabs at both time points. At least 80% of the remaining slabs toward the reference band also had to pass, with no two consecutive failures. The reference band itself was retained. The implementation also requires sufficient data in the reference band and checks the stability of the depth limit.

We open-source our developed 2D/3D GPU-accelerated deformable registration library on Github: [*https://github.com/lukasglandorf/accel-deform-registration*](https://github.com/lukasglandorf/accel-deform-registration)

**SN 5: Depth-dependent signal and measurement availability**

Vessel-to-background contrast and background noise were evaluated separately from clean OCTA images. The contrast-to-noise ratio was:

$$CNR = \frac{median(vessel intensity) - median(background intensity)}{1.4826 \times MAD(background intensity)}$$

Background was measured in the same depth slice. Background noise was 1.4826 times its median absolute deviation. CNR, contrast and noise were summarized in 50-µm depth bands, requiring at least 30 capillaries per animal, session and band. The accepted-cohort analysis extended to 350 µm and reported the number of contributing animals per band. Acute/baseline ratios were first summarized within each animal, then combined with equal weight per animal. Uncertainty was estimated with 20,000 bootstrap resamples of animals. To test whether changes differed with depth, log-transformed signal metrics were modeled using time point, depth and their interaction. Depth was centered at 150 µm and scaled in 50-µm steps. The model included an animal-specific intercept and depth slope and was fitted by maximum likelihood. The interval for the interaction was the fitted coefficient ±1.96 standard errors. Missing depth bands were not filled in. Changes in total detectable length within shared tissue were evaluated separately from individual-vessel fate. These controls are shown in Supplementary Fig. 7.

**SN 6: OCTA-derived diameter and its precision**

The distance field calculated from the segmentation defined the sampling window around the smoothed centerline. Intensity profiles across the vessel were taken from the original OCTA volume and fitted with a Gaussian:

$$f(x) = B + A exp[-((x - x₀) / w)^{2}]$$

Here, $x$ is position across the vessel, $B$ is background intensity, $A$ is amplitude, $x₀$ is the profile center and $w$ is the width parameter. Individual fits required R² > 0.8, and a segment required accepted widths at at least 30% of sampled positions. Accepted widths were averaged along the segment and expressed in micrometers. Capillary diameters that failed the measurement-source or fit-quality requirements remained missing. Large pial and penetrating vessels could use the segmentation-derived distance transform estimate when the Gaussian model was unsuitable. Paired diameter comparisons required compatible measurement sources at both time points.

The reported diameter is the fitted width $w$, not the Gaussian standard deviation or full width at half maximum. It describes the width of the OCTA signal rather than directly locating the anatomical lumen walls.

Optical resolution and measurement precision describe different properties. The approximately 3-µm lateral optical resolution describes the spatial detail resolved by the imaging system. It does not set a minimum uncertainty for estimating the width of a vessel's intensity profile.

A small change in width, $\delta w$, changes the intensity at sample $i$ by approximately:

$$\delta fᵢ \approx gᵢ \delta w$$

$$gᵢ = \frac{\partial f(xᵢ)}{\partial w} = \frac{2A(xᵢ - x₀)^{2}}{w^{3}} \cdot exp[-\left( \frac{(xᵢ - x₀)}{w} \right)^{2}]$$

Here, $gᵢ$ describes how strongly intensity changes with width at that position. For this simplified derivation, the other fit parameters are held fixed. With independent, zero-mean intensity errors $\varepsilonᵢ$ of variance $\sigma^{2}$, local least-squares fitting gives:

$$ŵ - w \approx\frac{\Sigmaᵢ gᵢ \varepsilonᵢ}{\Sigmaᵢ gᵢ^{2}}$$

$$Var(ŵ) \approx\frac{\sigma^{2}}{\Sigmaᵢ gᵢ^{2}}$$

Here, $ŵ$ is the estimated width. Small width changes can therefore be detected through continuous changes in the intensity profile, even when they are smaller than the optical resolution. Precision depends on signal sensitivity and noise. In practice, amplitude, center and background are also fitted, and their uncertainty contributes to width uncertainty. Averaging widths along a segment can reduce random error: for $M$ independent estimates with equal variance $s^{2}$, the variance of their mean is $\frac{s^{2}}{M}$. Correlation between measurements reduces this benefit, and interpolation does not create independent measurements.

We therefore measured repeatability using split-half reconstructions rather than assuming ideal noise averaging (Extended Data Fig. 1b). For each dataset, time point and split, the SD of paired half-reconstruction differences was divided by $\sqrt{2}$ and then by the empirical frame-scaling factor of $1.072$ to estimate the full-average measurement SD. These estimates were combined with weights proportional to the number of eligible vessel pairs. The measurement SD was 0.262 µm. Assuming equal, independent errors in two measurements, their 95% minimum detectable difference was:

$$MDC₉₅ = 1.96\cdot\sqrt{2}\cdot\sigmaₘ \approx0.726 \mu m$$

Here, $\sigmaₘ$ is the measurement SD. These estimates describe within- time point repeatability of OCTA-derived diameter. They do not include all systematic or between- time point errors and do not establish anatomical diameter accuracy below 3 µm. Smaller average differences between groups can still be estimated, with uncertainty assessed across animals rather than treating capillaries as independent biological replicates.

**SN 7: Derived hemodynamic estimates of shear and pressure**

Exploratory indices of wall shear stress and pressure drop along a segment were calculated as:

$$\tau=\frac{4\mu v̄}{r}$$

$$\Delta P=\frac{8\mu Lv̄}{r^{2}}$$

Here, $\mu= 0.0035 Pas$ is the assumed viscosity, $L$ is segment length, and $r$ is radius, using measurements that passed the source and quality checks. The cross-sectional mean velocity was approximated as $v̄ = v / 1.5$. This scaling was confined to these algebraic indices and did not alter the reported direct Doppler velocity. Resistance proportional to $L/r⁴$ described relative ranking, with the adopted diameter uncertainty propagated. These indices were not independent measurements of pressure, shear or a cellular mechanism.

**SN 8: Vessel matching and measurements across split or merged segments**

Baseline vessel segments were matched to segments in the registered and aligned follow-up graph. Matching allowed one-to-one relations, splits, mergers and combinations of these, while retaining the original segment identifiers. Velocity and diameter were then retrieved from the original images and graphs. A segment split or merger did not by itself indicate vessel loss, and each baseline segment remained a single observation.

Centerline overlap was measured in both directions: from baseline to follow-up and from follow-up to baseline, using a 4-voxel (approximately 7.8-µm) nearest-point search radius. Candidate relations required at least 0.15 overlap in each direction, a geometric mean of the two directional fractions of at least 0.10, and a relative score of at least 0.50. The relative score was the larger directional overlap fraction and supplied the match score used below. All qualifying relations were retained.

When a baseline segment matched several follow-up segments, their measurements were combined using the match score multiplied by follow-up segment length as the weight. Each baseline segment contributed once to the paired comparison. Velocity and diameter comparisons required valid measurements at both time points and diameter additionally required compatible measurement sources. Measurements were compared within the same final vessel class. Missing measurements were not replaced with zero. A strict one-to-one subset was used only for analyses explicitly requiring unique segment identity, including the designated topology-transition comparison.

The combined follow-up measurement was the weighted geometric mean:

$$wᵢⱼ = sᵢⱼ \times Lⱼ$$

$$y'ᵢ = exp[(\Sigmaⱼ wᵢⱼ ln(yⱼ)) / (\Sigmaⱼ wᵢⱼ)]$$

And

$\Deltaᵢ = ln(y'ᵢ / yᵢ)$.

Here, $i$ identifies the baseline segment, $j$ identifies an eligible follow-up segment, $sᵢⱼ$ is the match score, $Lⱼ$ is the follow-up segment length, $yⱼ$ is its positive post-stroke measurement, $yᵢ$ is the positive baseline measurement, and $\Deltaᵢ$ is the paired log change. The sums include only segments passing the measurement-quality, source and vessel-class requirements, so weights are renormalized after these restrictions. Capillary comparisons additionally required an assessable centerline fraction of at least 0.50. Other vessel classes used accepted matches without this additional 0.50 centerline-support requirement. These measurement requirements were separate from stall classification.

**SN 9: Network position and vessel classification**

Network connectivity and branch order were calculated on each complete original graph before selecting vessels with available measurements or perfusion classifications. Baseline predictors used the baseline graph. Network comparisons between time points used independently defined shared tissue and vessel matching was required only for comparisons following individual vessels.

Relative arterial–venous position: $rBO = BOₐ / (BOₐ + BOᵥ)$

Nearest-penetrating-vessel order: $min(BOₐ, BOᵥ)$

Total branch order: $tBO = BOₐ + BOᵥ$

Here, $BOₐ$ and $BOᵥ$ are arterial and venous branch order. Physical-distance alternatives used the distance along connected vessels to the arterial and venous interfaces, rather than straight-line distance. Sensitivity to the imaged field was assessed with the reference-network crops described in SN 11.

Subtype-specific stall risk required a classified baseline subtype and assessable subsequent fate, without requiring a classified follow-up subtype. A transition between subtypes required classification at both time points and the correspondence restriction specified for that analysis. Unclassified vessels remained available for outcomes not requiring subtype.

Vessel-class labels were reconciled across time points using all accepted matches, weighted by match score multiplied by the source segment length. A segment originally labeled as a capillary could receive a specific pial or penetrating-vessel label only when that class accounted for at least 80% of the total weighted evidence and evidence for the opposite arterial or venous class was at most 10%. The denominator included capillary and unclassified source classes. Existing non-capillary labels were retained. If different time points suggested conflicting changes to a baseline label, the original label was retained. These label weights used source segment length, whereas the measurement weights in SN 8 used follow-up segment length. This analysis labeling did not change the graph-model or pial-optimization procedure. When no matching evidence was available, the original label was retained.

**SN 10: Arterial-venous capillary counts and shares**

Baseline capillaries and subsequent stalls were counted separately on the arterial (rBO <0.5) and venous (rBO >0.5) sides within each animal, using the same both-connected, resolved-fate population as Figure 5h. Percentages used all eligible baseline capillaries or all stalls, respectively, including vessels at $rBO=0.5$ in the denominators. Counts described sampled networks and were not converted to densities. The four paired side comparisons used two-sided animal-level t-tests with Holm correction across the four tests (Supplementary Fig. 11).

**SN 11: Field-of-view validation of branch order**

Three reference vascular networks were cropped at 42 overlapping lateral placements, keeping the upper plane fixed at the minimum centerline depth of each full network. Square lateral widths of 250, 500, 750 and 1,000 µm were crossed with depths of 50, 100, 200, 300, 360, 400 and 500 µm. Smaller fields retained the same lateral center. Additional depths of 150, 250, 350 and 450 µm at 1,000-µm width reproduced the original axial series. The same graph-processing and branch-order procedures were used across field sizes. For each arterial and venous order, crop-intersecting capillaries with finite full-network order were classified by whether the cropped order was exact, overestimated or unavailable. Results were separated for interior vessels, vessels crossing a crop face and all vessels. Fractions were averaged across eligible placements within each network and then equally across the three networks. Between-network SD described variability. Overlapping placements were not treated as independent biological samples. Surface-anchored results are shown in Extended Data Fig. 2 and Supplementary Fig. 10. Freely positioned crops were separate stress tests (Supplementary Fig. 10).

**SN 12: Changes in detectable capillary perfusion**

Vessel perfusion was assessed along baseline centerlines in tissue assessable at the time points being compared. At least 10 µm of physical centerline length had to be assessable. Follow-up perfusion evidence was present at a sampled position when the target segmentation was within 2 voxels or a target centerline was within 4 voxels. On the 1.95-µm analysis grid, these distances were approximately 3.91 and 7.81 µm, respectively. The fraction lacking follow-up evidence was calculated from physical centerline length, rather than the number of segments. Less than 40% of comparable arc lacking both forms of evidence indicated present perfusion, 40% to less than 60% remained uncertain and at least 60% was a candidate loss.

Candidate losses were checked for residual local OCTA signal and overlying-vessel shadowing. Residual signal was evaluated along the longest assessable centerline stretch lacking nearby follow-up segmentation or a centerline. To allow for small alignment errors, the centerline was shifted by each integer-voxel displacement within a radius of 4 voxels. Background samples were taken 3 voxels away in the six coordinate directions and pooled over the sampled stretch for each displacement. Centerline intensity was standardized as:

$$z = \frac{I - median(B)}{max\left[ 1.4826 \times MAD(B), 10⁻¹^{2} \right]}$$

Here, $I$ is centerline intensity, $B$ contains the background samples, and MAD is their median absolute deviation. For each displacement, the 25th percentile of centerline $z$ was calculated and the largest of these values was retained. A value >0.30 made the loss candidate uncertain. For the shadow check, the maximum of the baseline and target 90th-percentile overlying segmentation radii, assessed over tissue at and above the centerline within a neighborhood extending 6 voxels in each lateral direction, had to be ≤8 voxels. Candidates exceeding this threshold remained uncertain. A high-confidence loss required at least 60% of assessable length to lack follow-up evidence and both uncertainty checks to pass. Vessels with insufficient assessable length were labeled not assessable.

These time point-level states describe loss of detectable OCTA perfusion. Repeated acquisitions average the perfusion signal over minutes and do not measure the duration of an individual flow interruption, establish complete red-cell arrest or distinguish persistent non-perfusion from anatomical vessel disappearance. Neither failure to obtain a vessel match nor a Doppler-velocity cutoff defined a stall (Supplementary Fig. 6). In general we differentiated between local stalls and full perfusion loss by examining if the surrounding vasculature was still present.

Individual-vessel analyses compared high-confidence loss with present perfusion. Uncertain states were retained when calculating sensitivity bounds. Net loss of detectable vessels was calculated separately: the same structural non-detection rule was applied from baseline to follow-up and then in the reverse direction. Each direction used its own assessable starting-vessel population as the denominator. Within each animal as: Net difference in detectable-vessel fractions = forward non-detection fraction − reverse non-detection fraction. Versions weighted by vessel count and by physical centerline length were reported separately. Because the two fractions use different starting populations, their difference is not the fraction of baseline vessels that stalled. This comparison did not change any individual vessel's classification. Total detectable network length was also measured independently at each time point within the shared tissue, without requiring vessel matches.

**SN 13: Regional perfusion and longitudinal comparisons**

LSCI perfusion change was normalized to each animal's baseline after registration. Regions were defined on the full smoothed LSCI field before applying the OCTA footprint. Perfusion core was a reduction of at least 70%. Outside core, hypoperfusion was divided into severe (at least 45% but less than 70% reduction) and moderate (at least 20% but less than 45% reduction) bands. Tissue with less than 20% reduction and hyperperfusion above 20% increase were identified separately. Signed distance to the core boundary described position relative to the core. Distance analyses required a measurable core in the full LSCI field, which could lie outside the OCTA footprint. No core distance was assigned when no core was identified. LSCI image coverage and tissue assessable by OCTA at both time points were checked separately (Supplementary Fig. 16).

Stall fractions and paired diameter and velocity changes were summarized within each animal and severity band before comparing animals. Continuous local LSCI change was assessed separately from the band comparisons and distance to the core. Velocity comparisons included only capillaries measurable at both time points (Extended Data Fig. 3). Animal-mean LSCI change and OCTA-detectable capillary-length change were compared across the eight animals with registered measurements (Supplementary Fig. 16).

The primary longitudinal comparison used the 1 h OCTA-defined region, fixed for the later time point. This core was derived from acute/baseline capillary-length retention using a 100-µm smoothing radius, with retention ≤20% defining core. Baseline vessels with at least 50% core membership were excluded from the non-core analysis. The LSCI-defined sensitivity used its own acute core definition, also fixed for day 7. These populations were analyzed separately (Supplementary Fig. 15).

Longitudinal analysis included three animals with accepted registration at baseline, the acute time point and day 7. Each included vessel had at least 80% of its baseline centerline length inside tissue assessable at all three time points, and a resolved perfusion state at both follow-ups. Eligible baseline vessels were classified as present at both follow-ups, loss followed by recovery, loss at both sampled follow-ups, or loss first detected at day 7. Subtype and network-position comparisons additionally required the corresponding baseline information. The association between acute and day-7 status was assessed by comparing the day-7 loss risk between the two acute-state groups within each animal, then combining the three animal risk differences. Uncertain outcomes were retained in the separate bounding analysis.

Baseline D-to-C stall risk was compared with the other classified baseline subtypes within each animal and time point. Local venous bias compared stall risk between the more venous and more arterial rank halves within baseline venous drainage groups. This was distinct from a network-wide comparison using a relative branch-order threshold of 0.5. Broader positional profiles used baseline nearest-penetrating-vessel branch order and fixed baseline arterial-venous rank quintiles, with risk centered on each animal's overall risk in the eligible topology population. Animal-level contrasts and uncertainty are specified in SN 15.

**SN 14: Baseline vessel properties and stall risk**

Within each animal, median baseline segment length was compared between capillaries subsequently classified as stalled and those remaining perfused, using vessels with both arterial and venous branch orders, resolved acute fate and finite positive length. Animal log ratios of these medians were combined and back-transformed. The separate marginal length association used an animal-specific logistic model with within-animal standardized log length. Baseline diameter, velocity and length associations without subtype and the jointly adjusted subtype analysis retained their own populations and model definitions (SN15; Supplementary Fig. 9; Supplementary Tables 3 and 4). Table 3 contains separate predictors. Table 4 contains the joint model.

**SN 15: Statistical analysis**

For the Figure 4 distribution tests, the baseline and follow-up empirical cumulative distributions were calculated separately within each animal. Their differences were averaged with equal animal weight, squared, and integrated over the pooled measurement distribution with equal animal weighting. The null distribution was obtained by evaluating all $2ⁿ$ reversals of baseline and follow-up labels at the whole-animal level, where $n$ is the number of animals. These tests assessed the full distributions separately from the paired summary-change tests.

Baseline-predictor models without subtype evaluated diameter, velocity and length separately. The diameter and velocity models included the focal log-transformed measurement, depth, log segment length, relative arterial-venous position and boundary proximity. The length model included log segment length, depth, relative arterial-venous position and boundary proximity. Continuous predictors were standardized within each animal. These models were distinct from the marginal length-only analysis in SN 14 and the joint subtype model below.

The joint subtype model included a binary D-to-C indicator, log diameter, log segment length, log velocity, arterial branch order, venous branch order, depth and boundary proximity. Continuous predictors were standardized within each animal. The physical-path alternative replaced the two branch-order terms with their corresponding path distances. Comparisons with and without velocity used the same velocity-complete vessels. Eligible animal fits required at least ten stalls and ten perfused vessels, at least twenty D-to-C and twenty other-subtype vessels, a full-rank design matrix and convergence. Fits with nonfinite coefficients or standard errors, absolute coefficients greater than ten or standard errors greater than five were excluded (Supplementary Table 4).

Holm correction was applied separately to the six Figure 4 distribution comparisons, the six subtype-pair comparisons within each phenotype or stall-risk family, the four subtype-fraction changes, and the nine arterial and venous branch-order risk comparisons. The two focal D-to-C tests from the branch-order and physical-path models formed a separate family from the exploratory correction across all eight joint-model coefficients. The four counts-and-shares comparisons used the family specified in SN 10. Other analysis-specific families, including the baseline-predictor screens and branch-order measurement changes, are identified with the corresponding tests in Supplementary Tables 3 and 4. Raw and adjusted P values are distinguished.

Confidence intervals for longitudinal mean risk differences and paired changes used a t distribution with n − 1 degrees of freedom. Where standardized paired continuous effects were reported, the effect was calculated as:

$$g_{z} = J(n - 1) \times mean(change) / SD(change)$$

Here, $J$ is the small-sample correction computed using the gamma function. Confidence intervals for standardized effects were obtained by inverting the noncentral t distribution. Ordinary 95% intervals accompanying Holm-adjusted P values were pointwise intervals, not simultaneous intervals.

**Supplementary Figure 1**

Segmentation 3D U-Net architecture

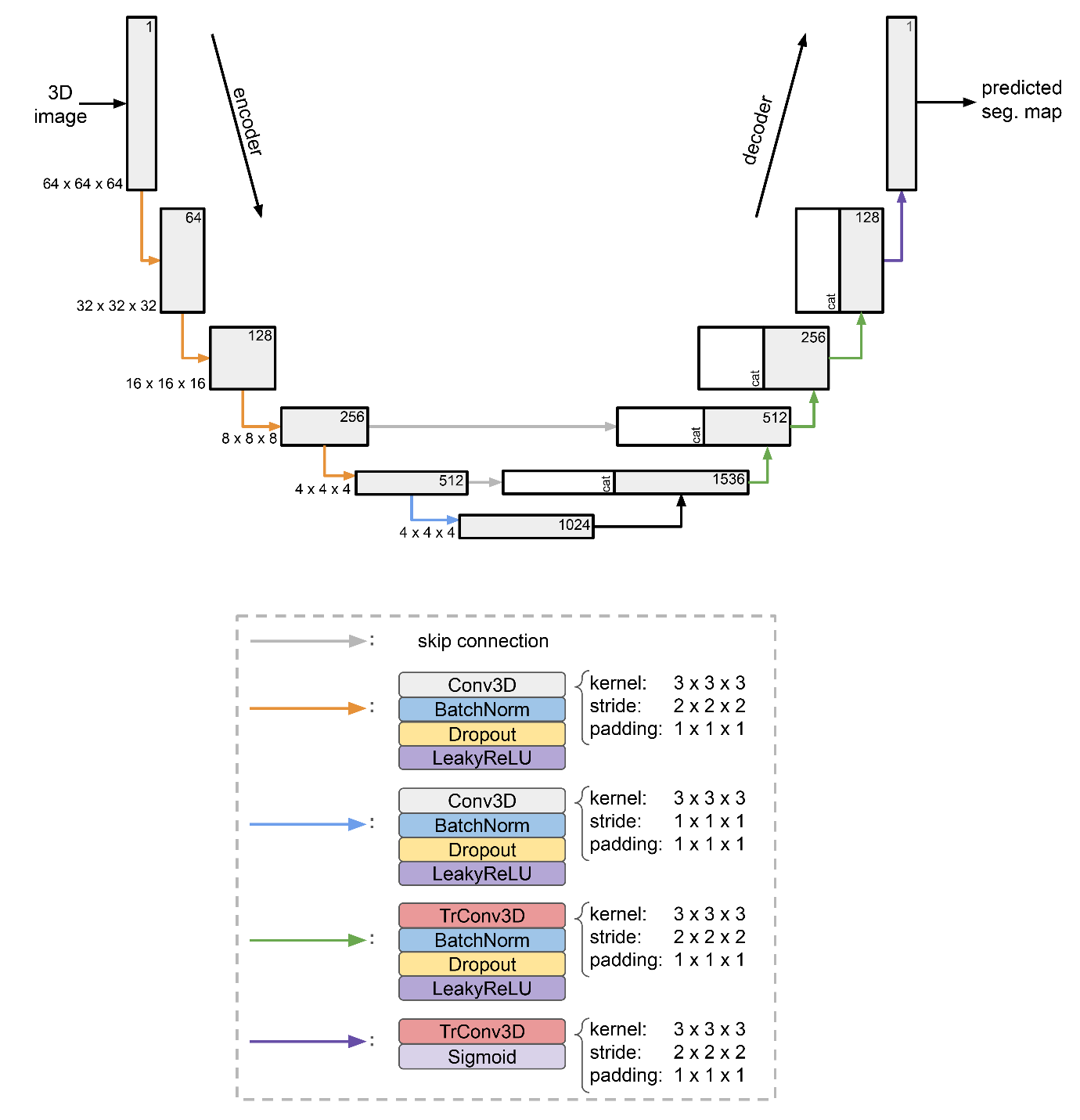

Figure S 1: Architecture of the employed 3D U-Net for segmentation.

**Supplementary Figure 2**

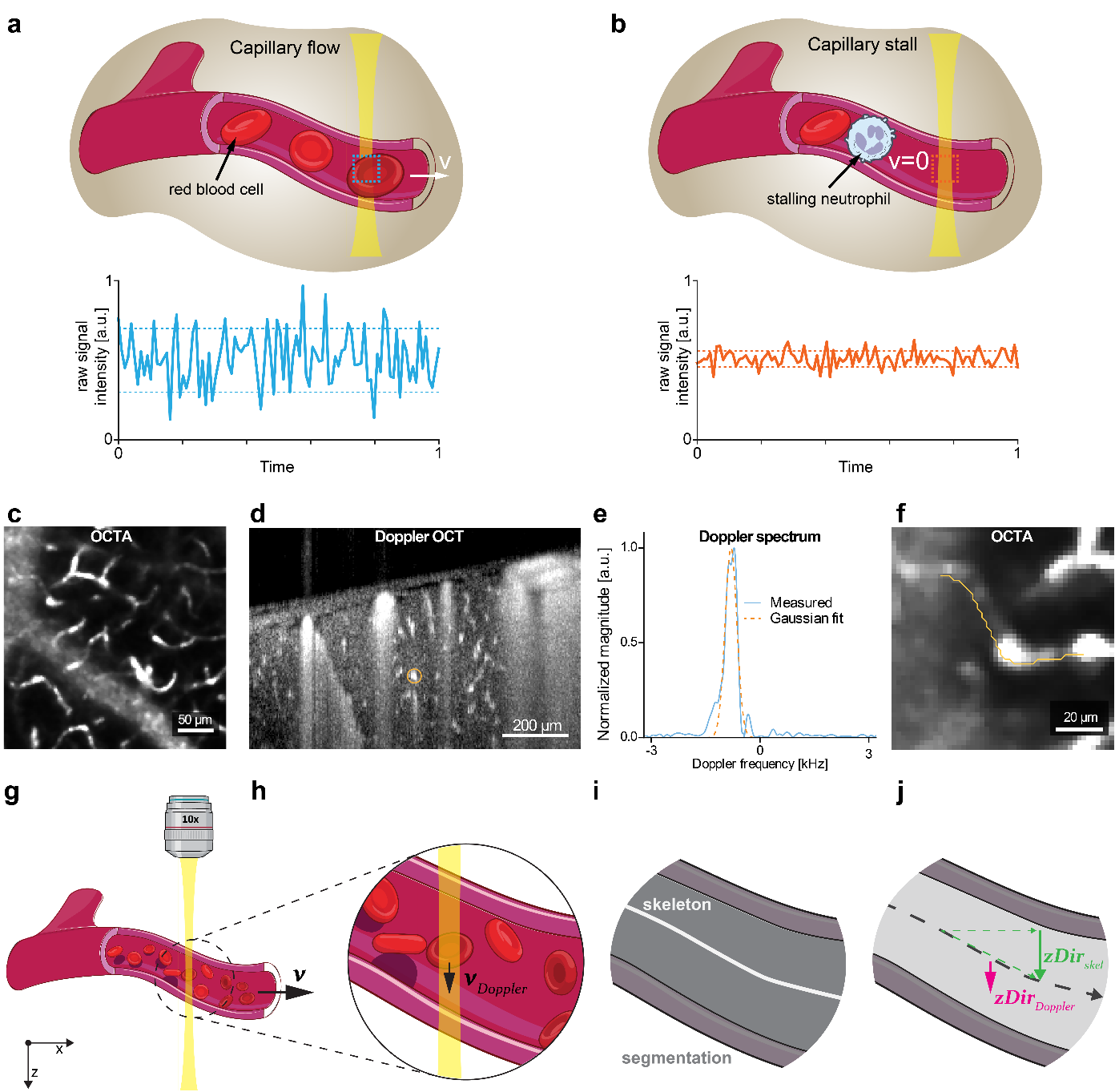

Figure S 2: Acquisition and flow direction. (a,b) Schematic of the OCTA signal. Moving red blood cells (RBCs) generate a time-varying signal. If a cell, such as a neutrophil, is stuck in a capillary, flow stops. Accordingly, the OCT signal varies less over time. The difference in time variations generate the OCTA contrast. (c) OCTA zoom-in. (d) Side-view of the Doppler intensity signal. The orange circle highlight the voxel displayed in (e). (e) The recorded Doppler spectrum and its Gaussian fit at the voxel highlighted in (d). (f) OCTA and skeleton of one capillary, describing the capillary sampling path. (g) Schematic side view of a vessel with flow predominantly along the x–z axes. Blood flow velocity (**v**) runs along the vessel’s centerline in one of two directions, designated “forward” or “backward.” (h) Magnified view of the xfOCM focal region. Doppler OCT (DOCT) measures only the axial component of blood flow (**v**_Doppler_), corresponding to the z-axis. (i) The vessel lumen is segmented using a custom 3D U-Net, producing a 3D map (gray). A skeletonization step yields centerlines (white), representing each vessel’s midpoint. (j) From this skeleton, local vessel orientation vectors are computed, reflecting how each segment is spatially constrained. By comparing the axial component of these orientation vectors (zDirD_skel_) to the DOCT-derived flow component (zDir_Doppler_), one determines whether the “forward” direction set for each vessel agrees with the true blood flow direction. This is done by ensemble averaging along all skeleton voxels of a blood vessel and additional weighting by the Doppler velocity magnitude, preferring regions with strong signal. If the two do not match, the flow is oriented opposite the vessel’s initially assigned direction.

**Supplementary Figure 3**

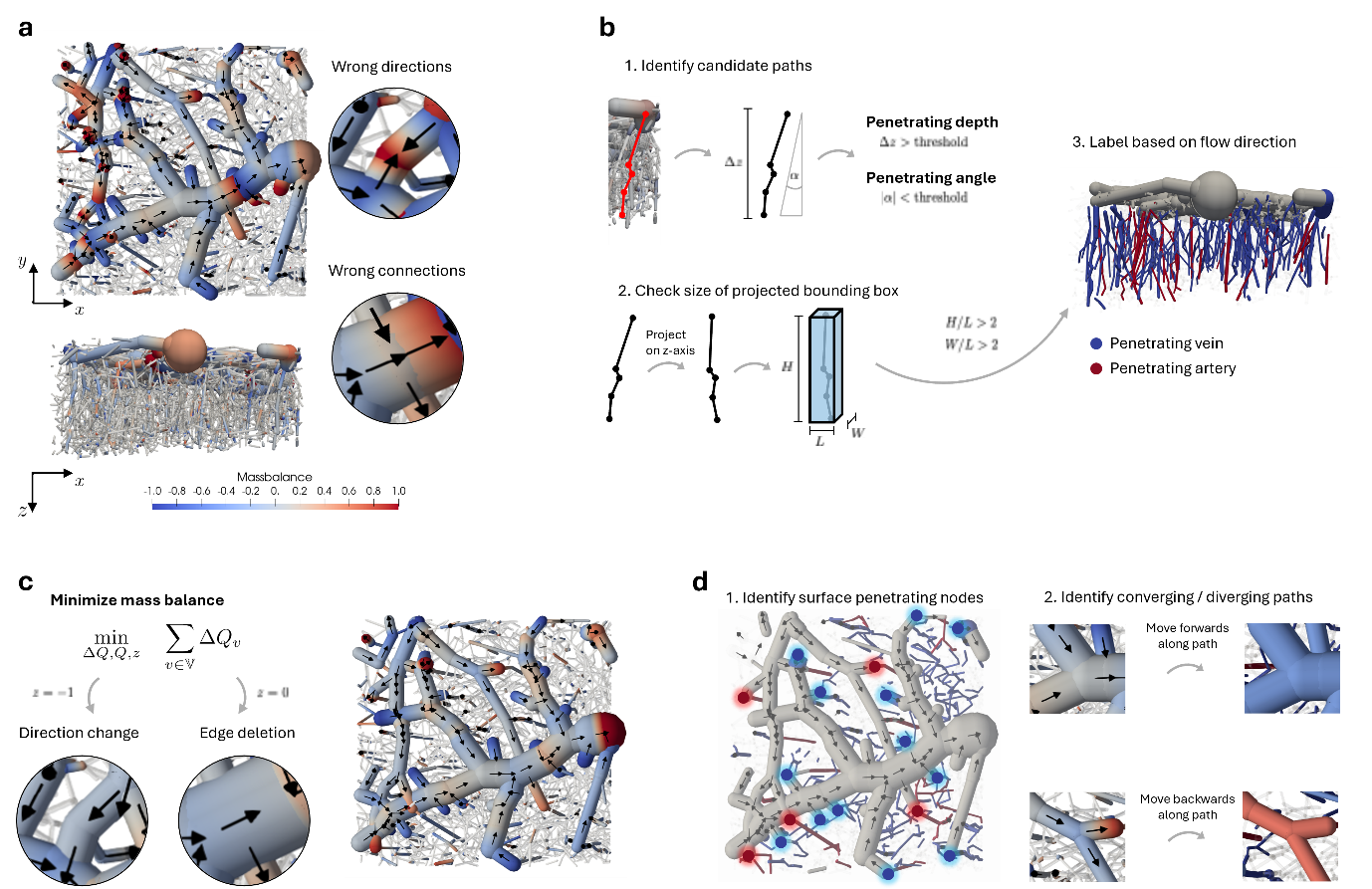
Graph-based AV-labeling algorithm

Figure S 3: Overview of the automatic labeling procedure of pial and penetrating vessels. (a) Top and side view of initial segmented data. Black arrows highlight the flow direction inside the pial vasculature. Wrong connections lead to significant errors in mass balance. (b) Schematic of identifying penetrating veins and arteries. Candidate segments are first identified by their global angle to the network and their penetration depth. Afterward, the overall geometry of a path is analyzed by checking if the projected bounding box is predominately vertical. Based on their flow direction, the remaining paths are labeled as either penetrating veins or penetrating arteries. (c) Optimization problem to fix errors in pial connections, consisting of the massflow error $\Delta Q$, mass flow $Q$ and decision variable $z$. Based on the result, connection stay intact ($z=1$), are reversed ($z=-1$) or deleted ($z=0$). Resulting optimized connections are shown on the right. (d) Overview of final pial vessel labeling. Surface penetrating nodes are highlighted in red and blue and are seeds to generate paths to (connected) boundary node. In each generated path, branching nodes are identified as either converging or diverging. Predominantly converging paths are labeled pial veins and diverging paths are pial arteries.

**Supplementary Figure 4**

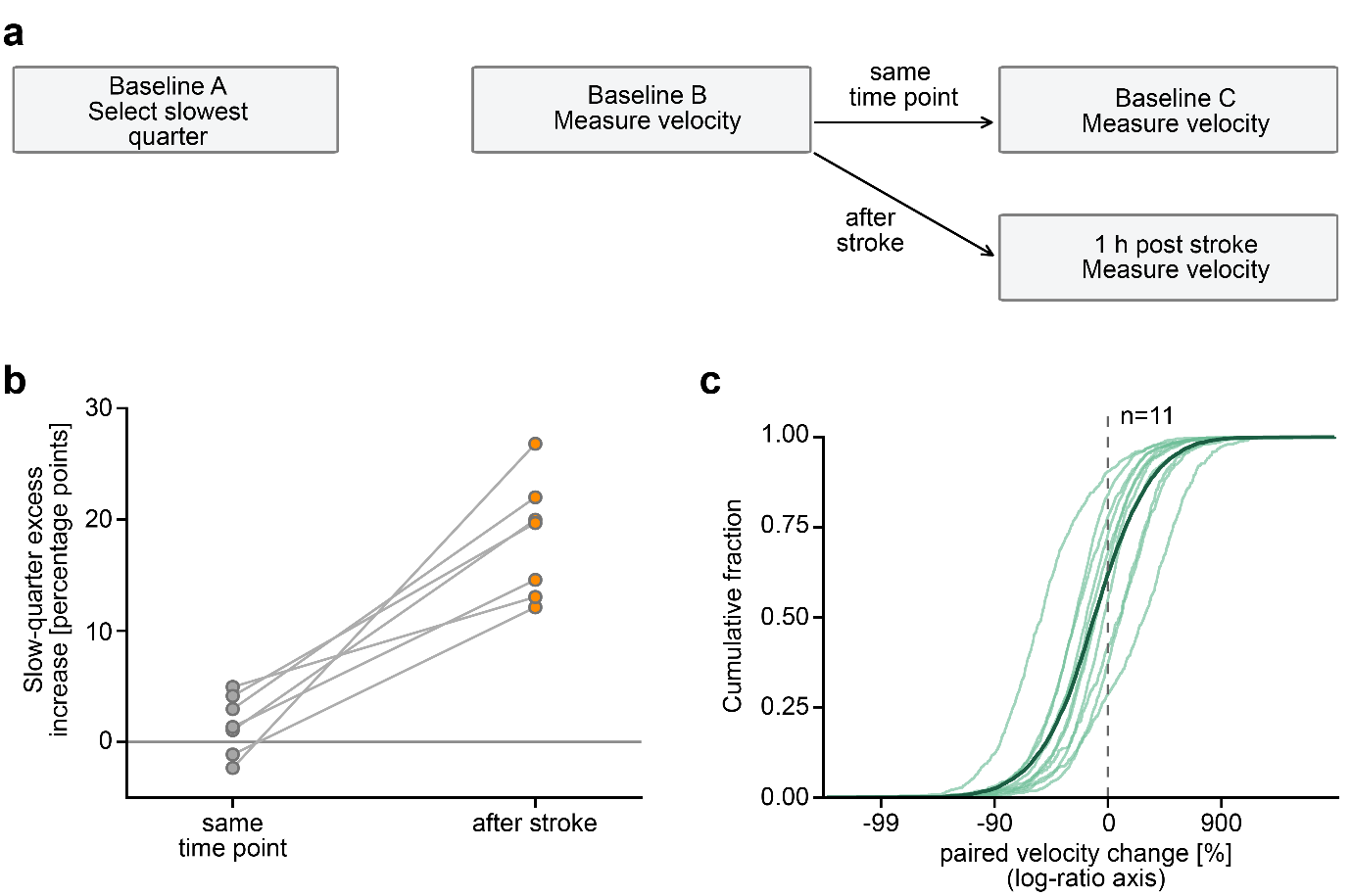

Figure S 4: Velocity increases and independent-frame controls. (a) Independent-frame design for selecting initially slow capillaries and measuring same-time point and post-stroke changes. Frame roles are rotated within animals. (b) Excess fraction accelerating in the slowest quarter versus the remaining capillaries, among 4,319 frame-complete, one-to-one capillaries in seven mice. The excess was 16.7 percentage points greater after stroke than in the same-time-point control (95% animal-level t interval, 10.6-22.9). (c) Cumulative distributions of paired relative velocity change among 13,027 mapped capillaries in 11 mice. Pale lines show animals and the dark line their equal-weight mean. Log-ratio spacing displays the full tails with percentage tick labels. Split/merge targets are combined once per baseline identity.

**Supplementary Figure 5**

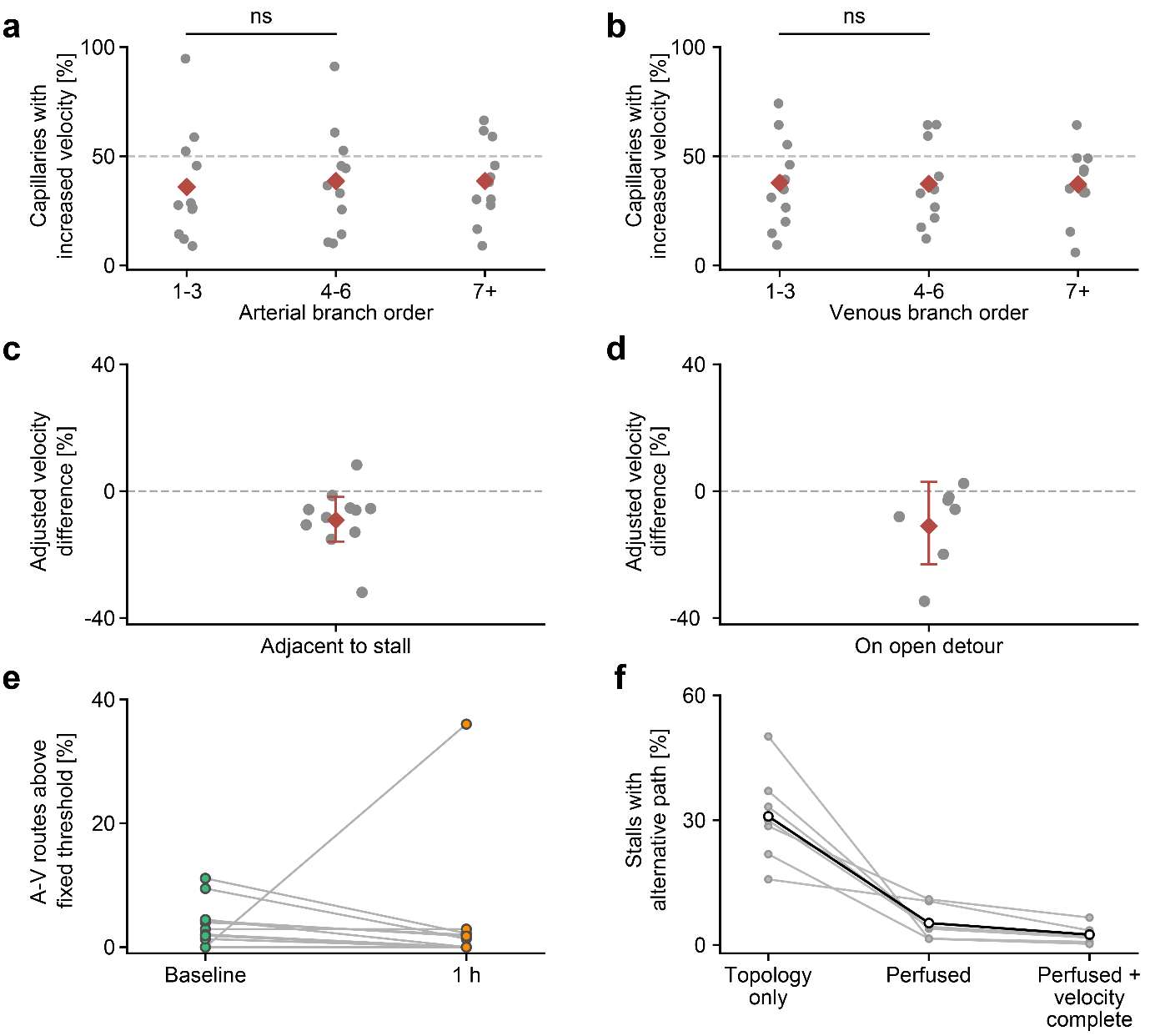

Figure S 5: Spatial distribution of velocity increases and alternative routes. (a,b) Fractions of measurable capillaries with increased velocity by baseline arterial (a) and venous (b) branch order in 11 mice. Gray points show animals and red diamonds their means. Brackets compare orders 4-6 with 1-3 using Holm correction; ns, not significant. (c,d) Adjusted velocity differences associated with direct adjacency to a stall (c; 11 mice) or an open alternative path around a stalled segment (d; seven mice). Gray points show animal estimates, red diamonds and bars show equal-animal estimates and 95% confidence intervals. (e) Fractions of baseline-defined, velocity-complete artery-to-vein routes exceeding each animal’s fixed baseline mean-plus-three-SD velocity threshold before and after stroke (f) Fractions of stalled segments with a topologically available, perfused or perfused-and-velocity-complete alternative path in seven mice. Gray lines show animals and black circles their means.

**Supplementary Figure 6**

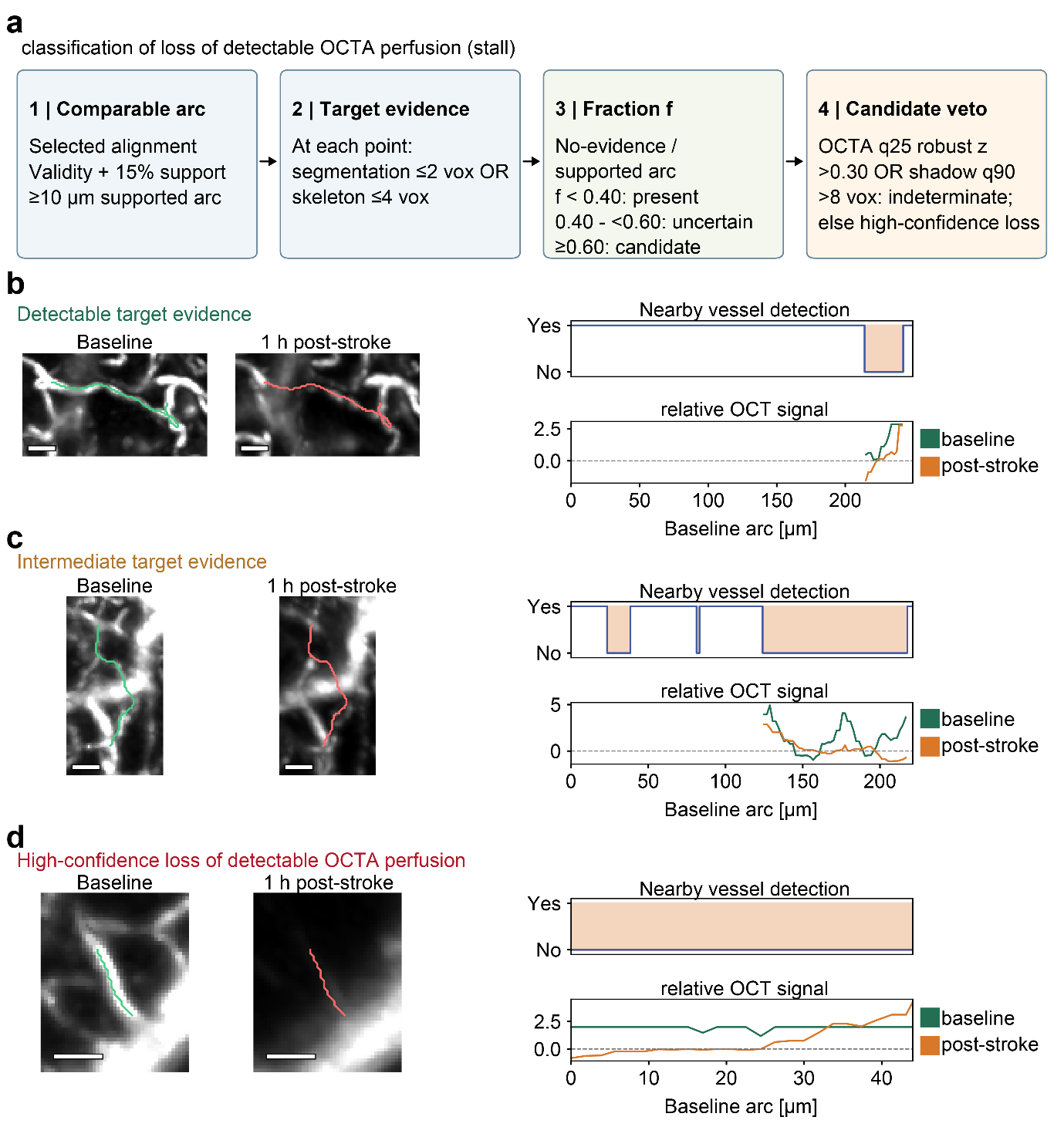

Figure S 6: Operational stall definition and examples. Workflow and three measured examples for classifying loss of detectable OCTA perfusion. Baseline and acute projections use the same contrast within each pair. Green and red lines mark the baseline capillary skeleton. The examples show detectable, indeterminate and high-confidence loss outcomes. Lower traces show baseline (green) and acute (orange) local OCTA signal along the longest interval lacking follow-up evidence. Scale bars, 25 µm.

**Supplementary Figure 7**

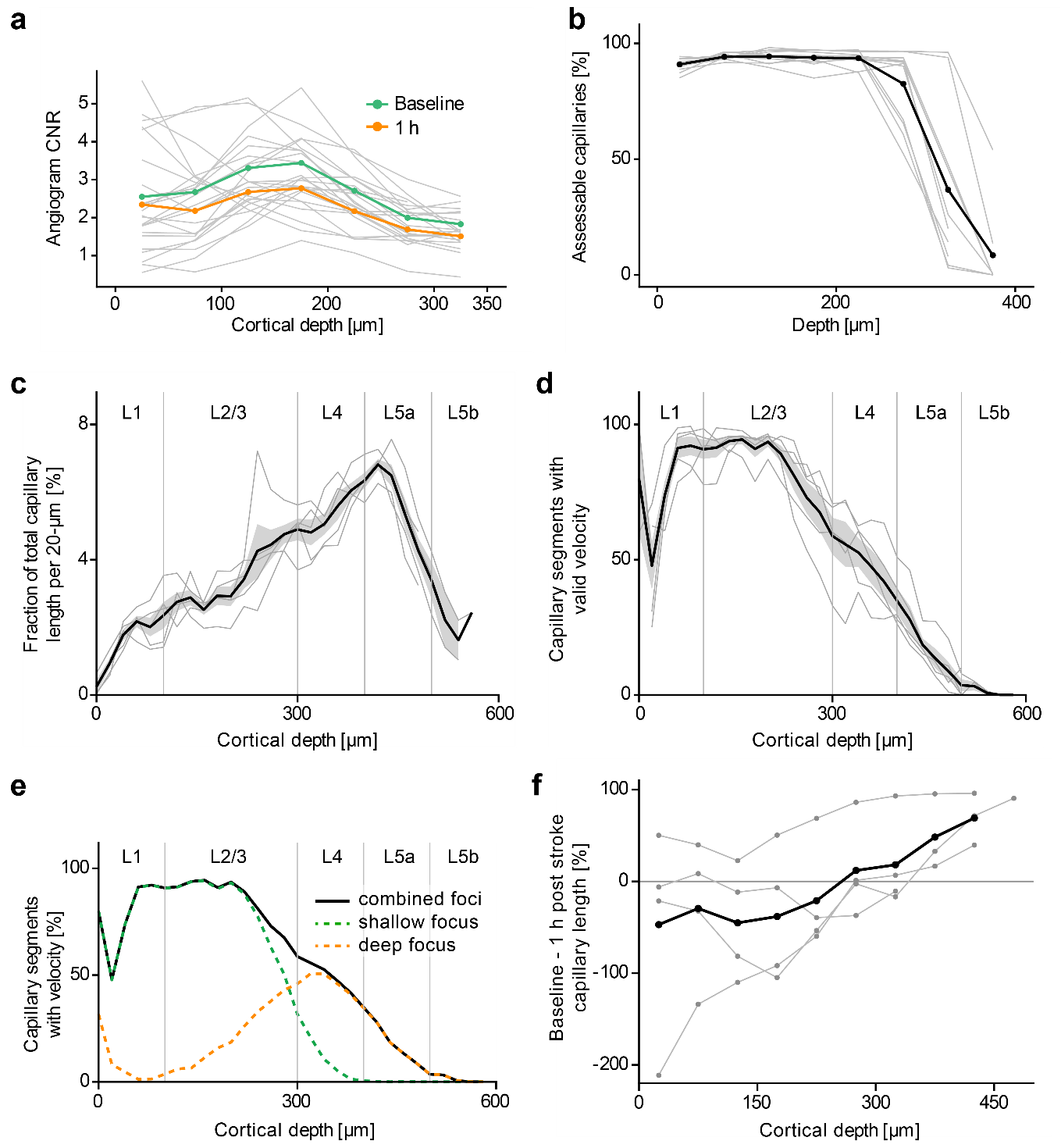

Figure S 7: Depth controls. (a) Baseline and 1-h contrast-to-noise ratio (CNR) in 50-µm depth bands. Gray lines show animals and colored lines show their means, with 12 animals below 300 µm and nine paired animals at 300-350 µm. (b) Tissue assessability by depth in 12 animals, with the equal-animal mean. (c) Detectable capillary length by depth in five stitched shallow/deep baseline acquisitions. Gray lines show animals and the black line and band show their mean ± SEM. Cortical-layer guides are approximate. (d,e) Velocity availability among detectable capillaries in the depth stitched acquisitions. (f) Baseline-minus-1-h detectable capillary length in four registered acute pairs, measured in shared 50-µm slabs without individual-vessel matching. Gray lines show animals and black shows their mean.

**Supplementary Figure 8**

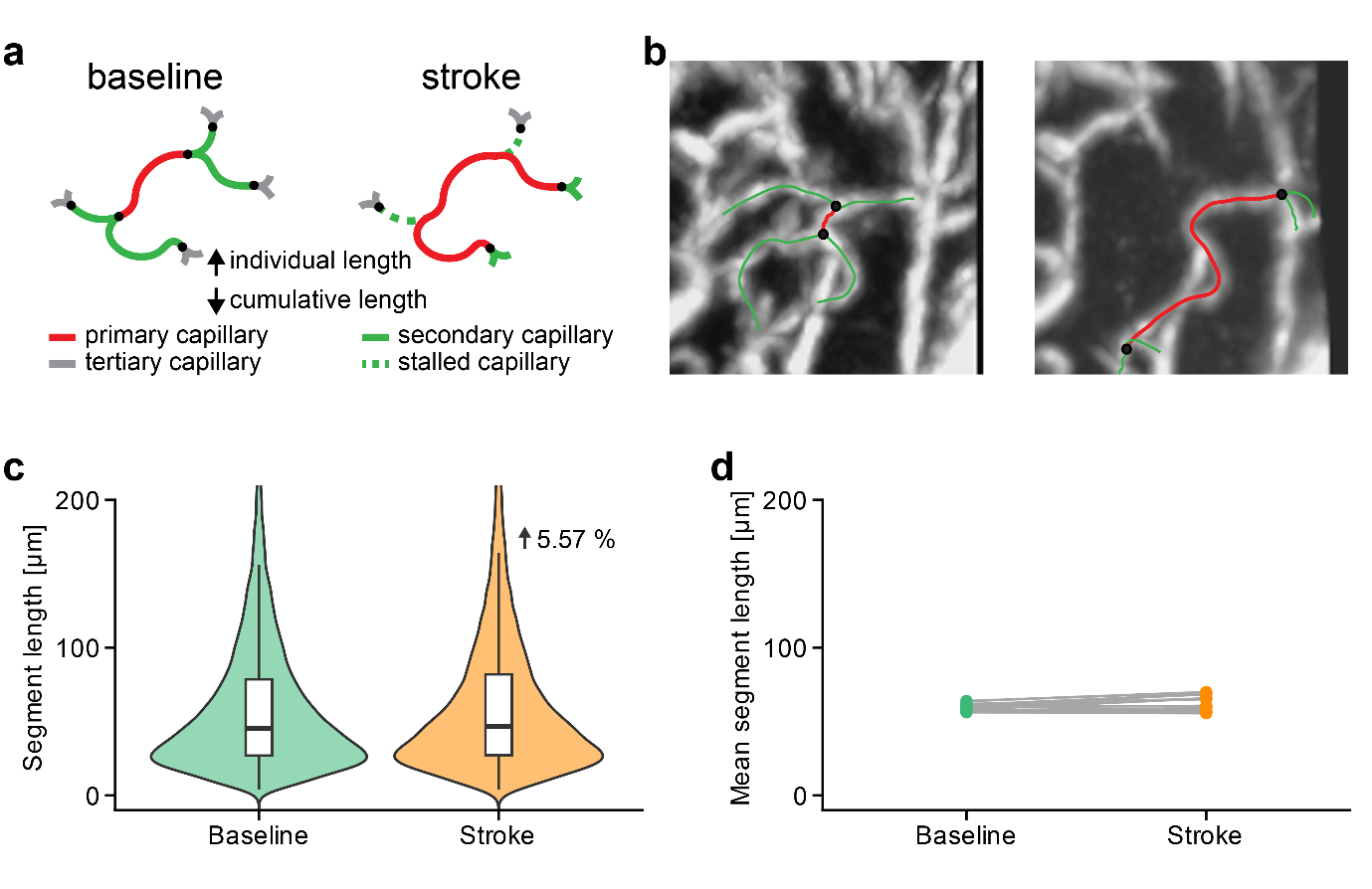

Figure S 8: Capillary graph-segment length after stroke. (a) Schematic of effective vessel segment fusion after adjacent stalls. (b) OCTA examples illustrate changes in detected segment organization. (c) Violin plots show baseline and 1-h post-stroke segment-length distributions. Across 12 mice, mean segment length increased from 59.36 to 62.71 µm (+3.35 µm), with a mean within-animal increase of 5.57%. (d) mean segment length across mice before (baseline) and after stroke (stroke).

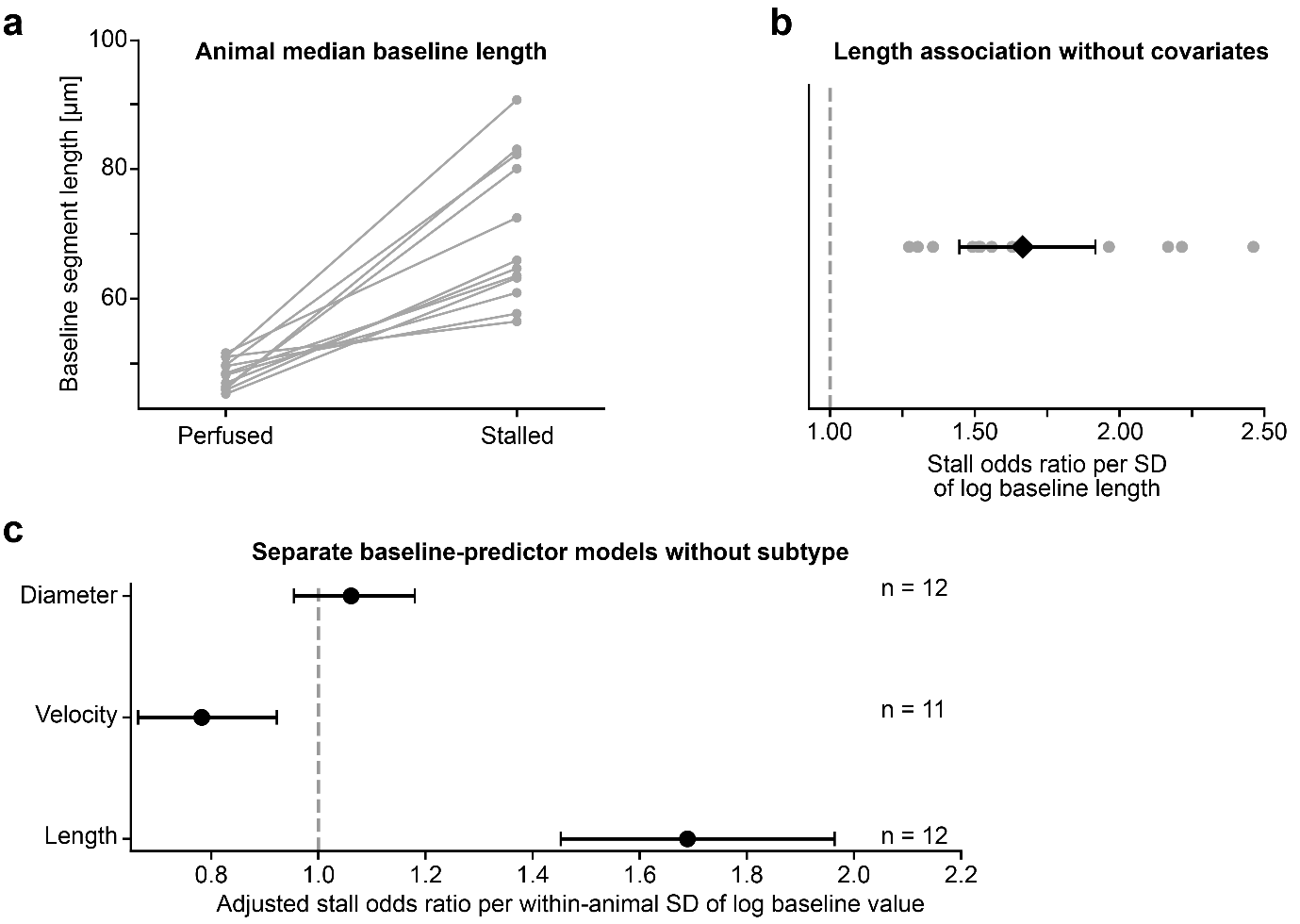
**Supplementary Figure 9**

Figure S 9: Baseline capillary properties and subsequent stall risk. (a) Baseline median segment length of capillaries remaining perfused or classified as stalled at 1 h in 12 mice. (b) Unadjusted association between baseline segment length and stall risk. (c) Separate associations of baseline diameter, velocity and length with stall risk, adjusted for depth, relative arterial-venous position and boundary proximity, without subtype. Diameter and velocity models additionally adjust for length. Predictors are log-transformed and standardized within animals. Gray points show animal estimates where plotted. Black marks show combined odds ratios and 95% confidence intervals. Panel c uses animal-level random-effects summaries with Knapp-Hartung inference. Model-specific populations and multiple-testing corrections are reported in Supplementary Table 3.

**Supplementary Figure 10**

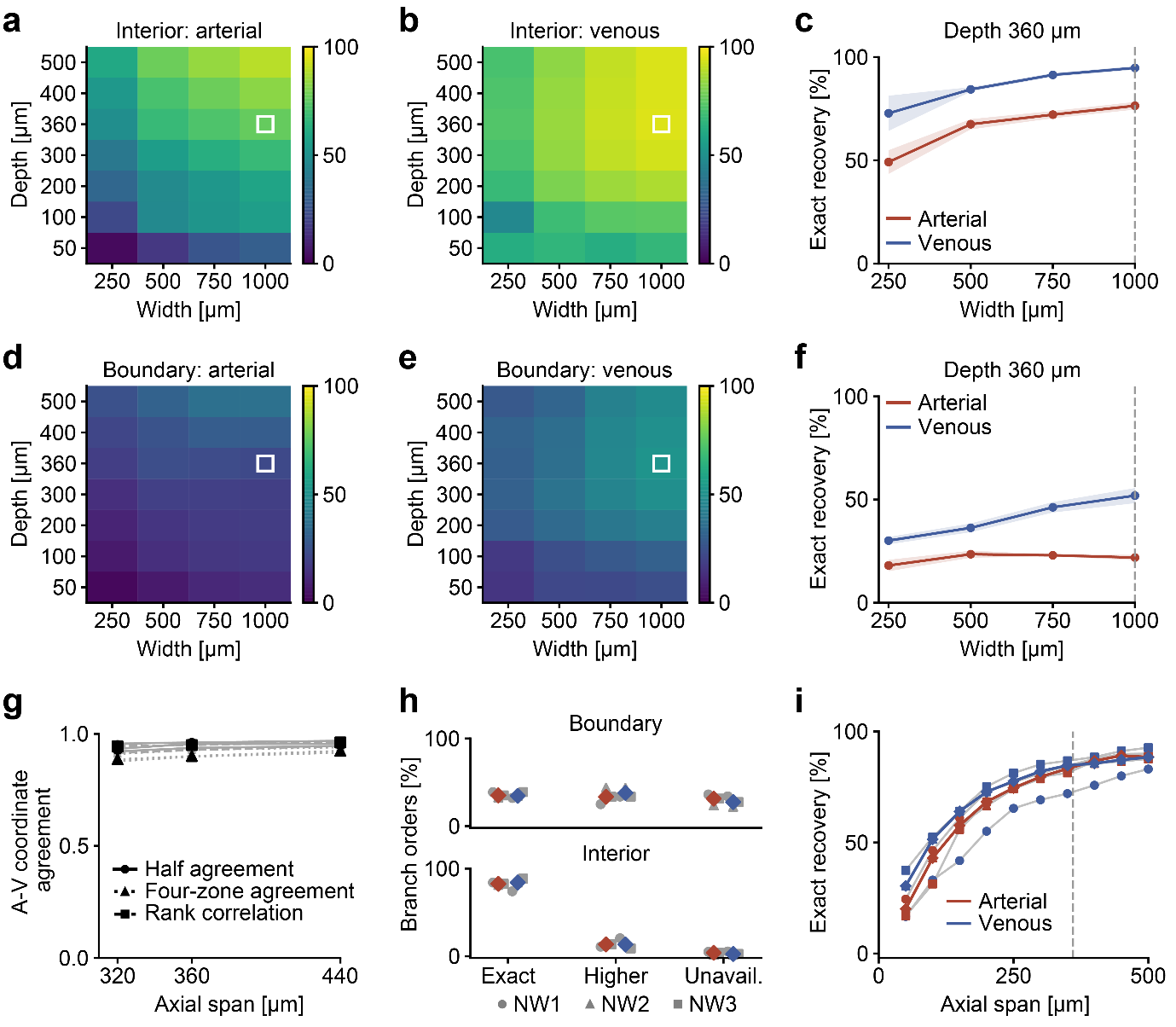

Figure S 10: Effects of imaging volume on network-position measurements from ex vivo reference networks (Blinder et al. 2013, Schmid et al. 2017). (a,b,d,e) Exact arterial (a,d) and venous (b,e) branch order recovery among interior (a,b) and boundary-crossing (d,e) capillaries in surface-anchored crops. Color scales show recovery percentages. White squares mark the experimental 1000 × 1000 × 360 µm field. Heatmap rows represent equally spaced tested depths. (c,f) Recovery versus lateral width at 360 µm depth for interior (c) and boundary-crossing (f) capillaries. Lines and shading show equal-network means ± SD across three reference networks and 42 overlapping crop placements. (g) Relative arterial-venous position recovery in freely positioned crops. Gray lines show individual networks and black symbols their equal-weight medians. (h) Exact, overestimated and unavailable branch orders in nominal freely positioned crops, separated by boundary-crossing and interior status. Gray symbols identify networks. Colored diamonds show equal-network summaries. (i) Exact-order recovery among interior capillaries across paired axial crop depths. Gray trajectories show network medians and colored lines their equal-network summaries. The dashed line at 360 µm. Red denotes arterial and blue venous measurements in c,f,h,i. Network symbols in h also apply to i.

**Supplementary Figure 11**

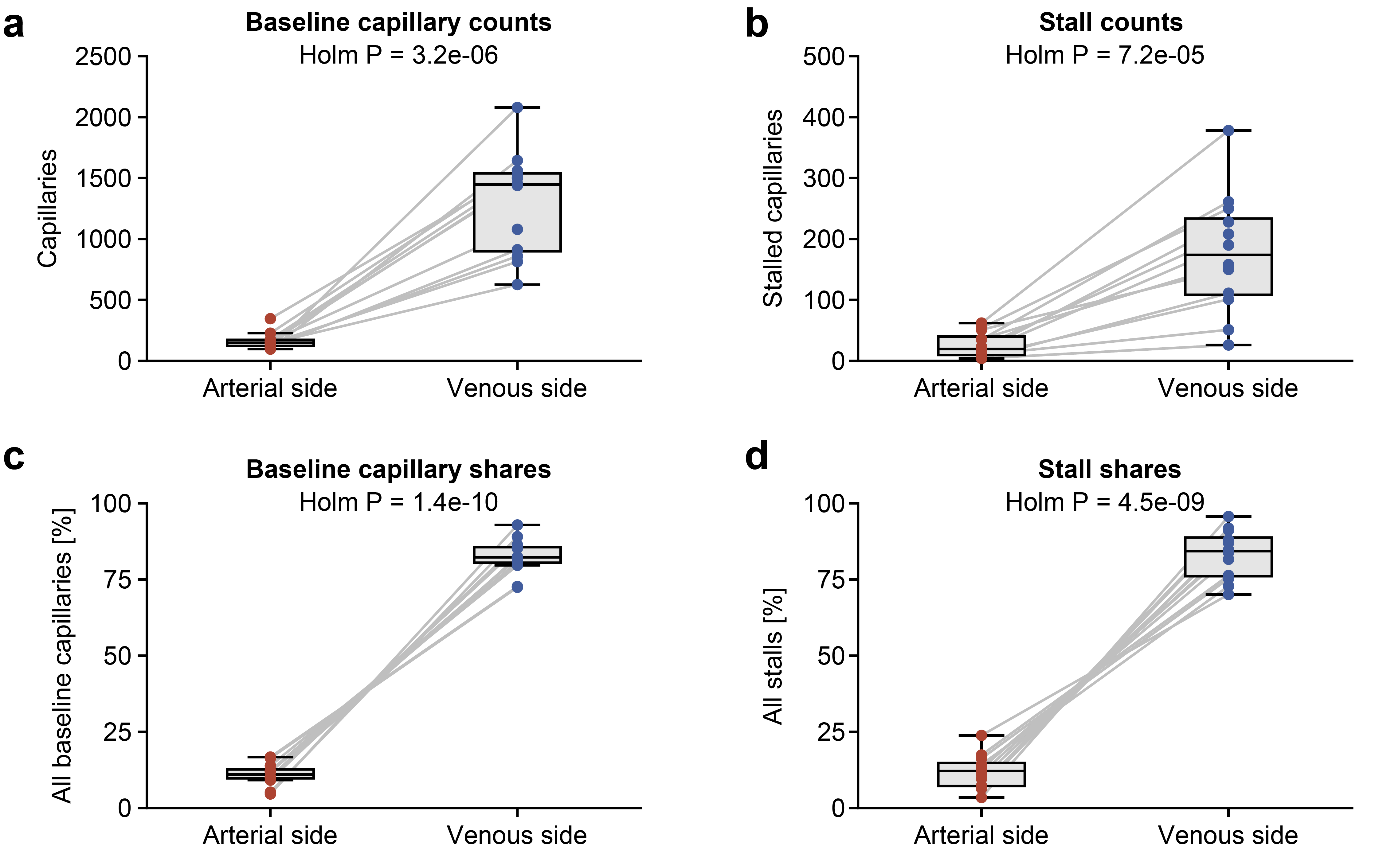

Figure S 11: Arterial-venous distribution of capillaries and stalls. (a,b) Baseline capillary counts (a) and acute stall counts (b) on arterial and venous sides, defined by relative branch order below and above 0.5, respectively. (c,d) Corresponding percentages of all baseline capillaries (c) and all stalls (d). Capillaries at exactly 0.5 remain in the percentage denominators. Points represent 12 mice. Boxes show medians and interquartile ranges, with whiskers extending to the most extreme values within 1.5 interquartile ranges. P values are from two-sided paired t tests with Holm correction across the four comparisons. The population is shared with Fig. 5h. Counts and percentages describe abundance, rather than individual capillary stall risk.

**Supplementary Figure 12**

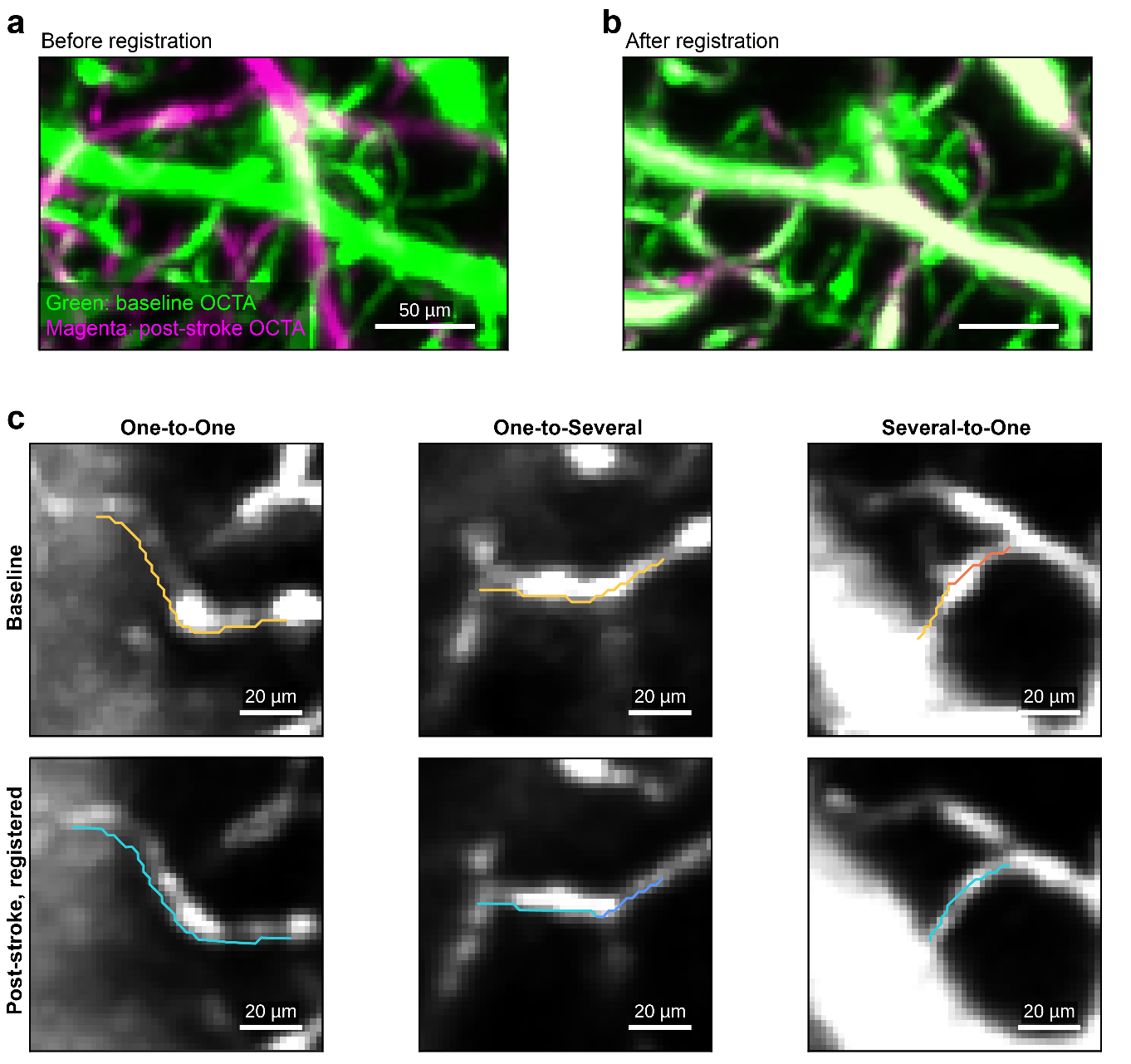

Figure S 12: Registration and vessel correspondence. (a,b) Baseline/acute OCTA overlays before lateral and deformable alignment and after registration. Both use the same depth projection and display settings, with baseline green and acute magenta. (c) Baseline and registered acute crops with centerlines showing one-to-one, one-to-several and several-to-one correspondence, respectively. Colored lines are the skeleton voxels of individual capillary segments.

**Supplementary Figure 13**

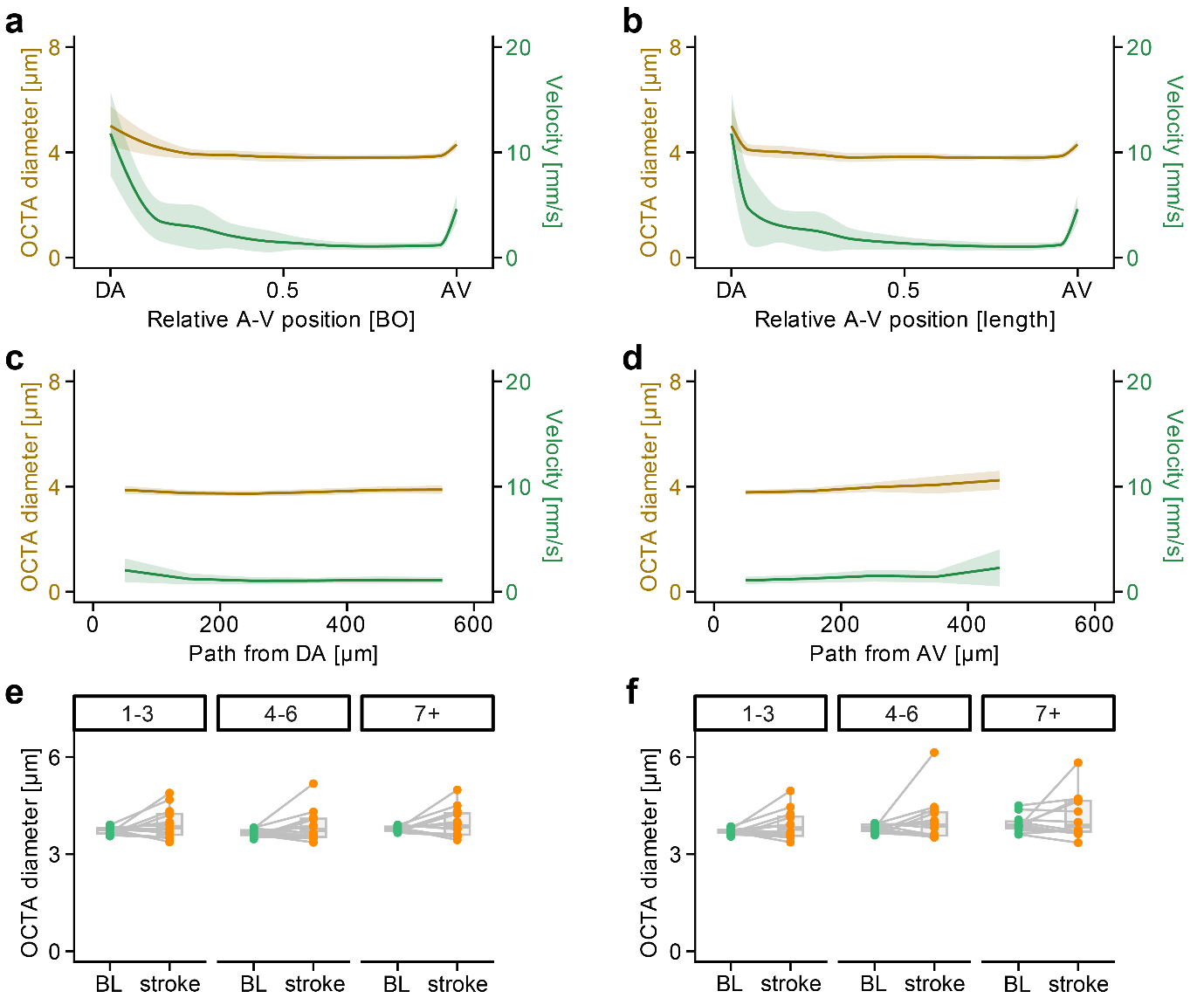

Figure S 13: Additional network coordinates and diameter profiles. (a,b) Diameter and velocity in capillaries along relative arterial-venous position defined by branch order (a) or shortest physical-path distance (b). (c,d) Corresponding profiles by shortest physical-path distance from descending arterioles (c) and ascending venules (d). Yellow curves show diameter on the left axes. Green curves show velocity on the right axes. Curves and shading show animal means ± SD, with 12 diameter-eligible and 11 velocity-eligible mice. (e,f) Baseline (BL) and 1-h post-stroke (stroke) diameter among paired capillaries, grouped by baseline arterial (e) or venous (f) branch order. Points show animal summaries and boxes show their median and interquartile range.

**Supplementary Figure 14**

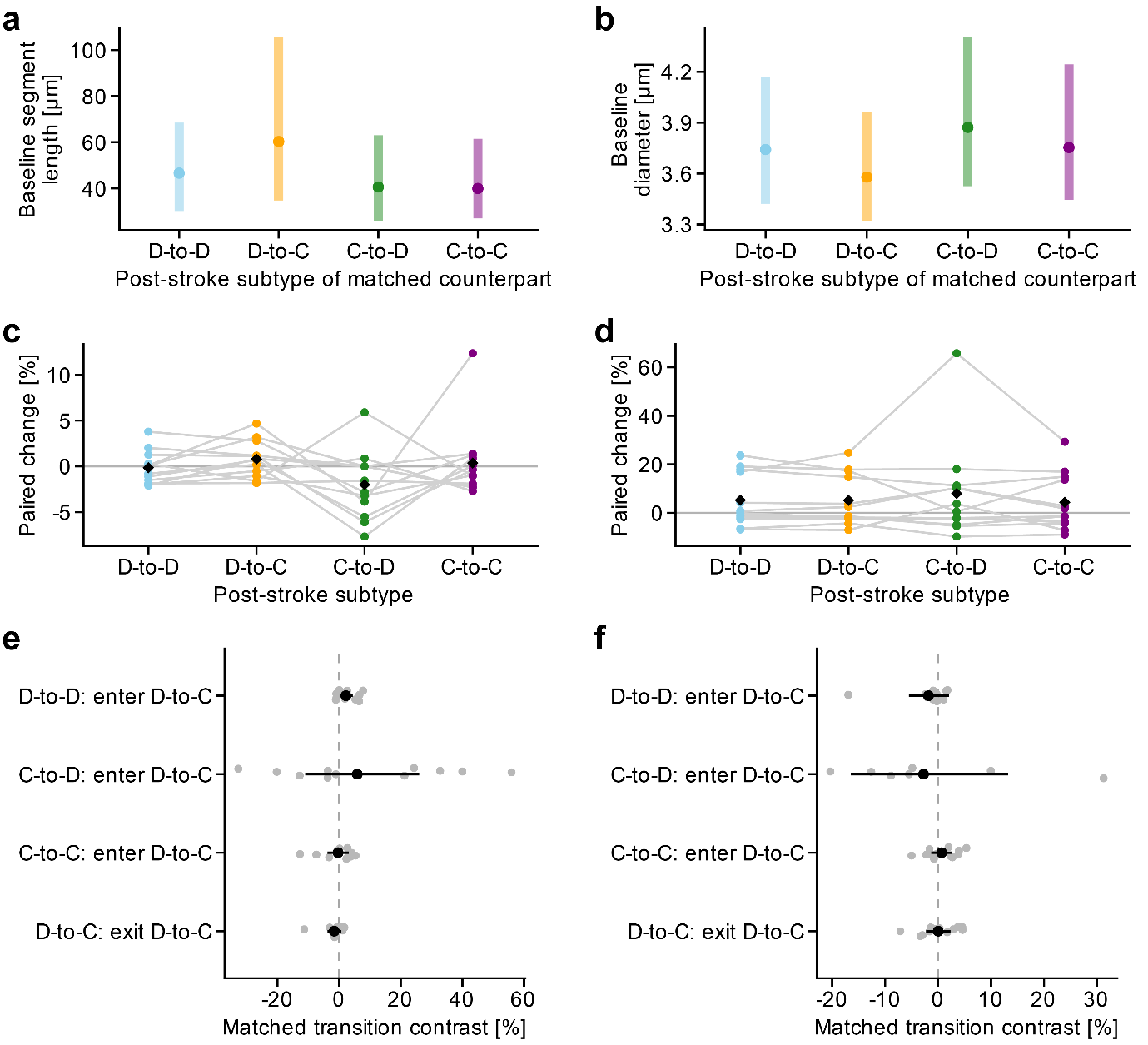

Figure S 14: Morphology of matched capillaries grouped by acute subtype. (a,b) Baseline segment length (a) and diameter (b) among one-to-one matched capillaries grouped by their 1-h post-stroke subtype. Points show equal-animal means of animal medians. Ranges show mean animal quartiles. (c,d) Paired length (c) and diameter (d) changes, calculated as animal median log ratios and transformed to percentages. Lines connect animal estimates and black diamonds show their means. (e,f) Length (e) and diameter (f) changes associated with entering or leaving the divergent-to-convergent (D-to-C) subtype, compared with retaining the starting subtype. Gray points show within-animal contrasts. Black marks show equal-animal estimates and 95% confidence intervals. Holm correction was applied within each measurement.

**Supplementary Figure 15**

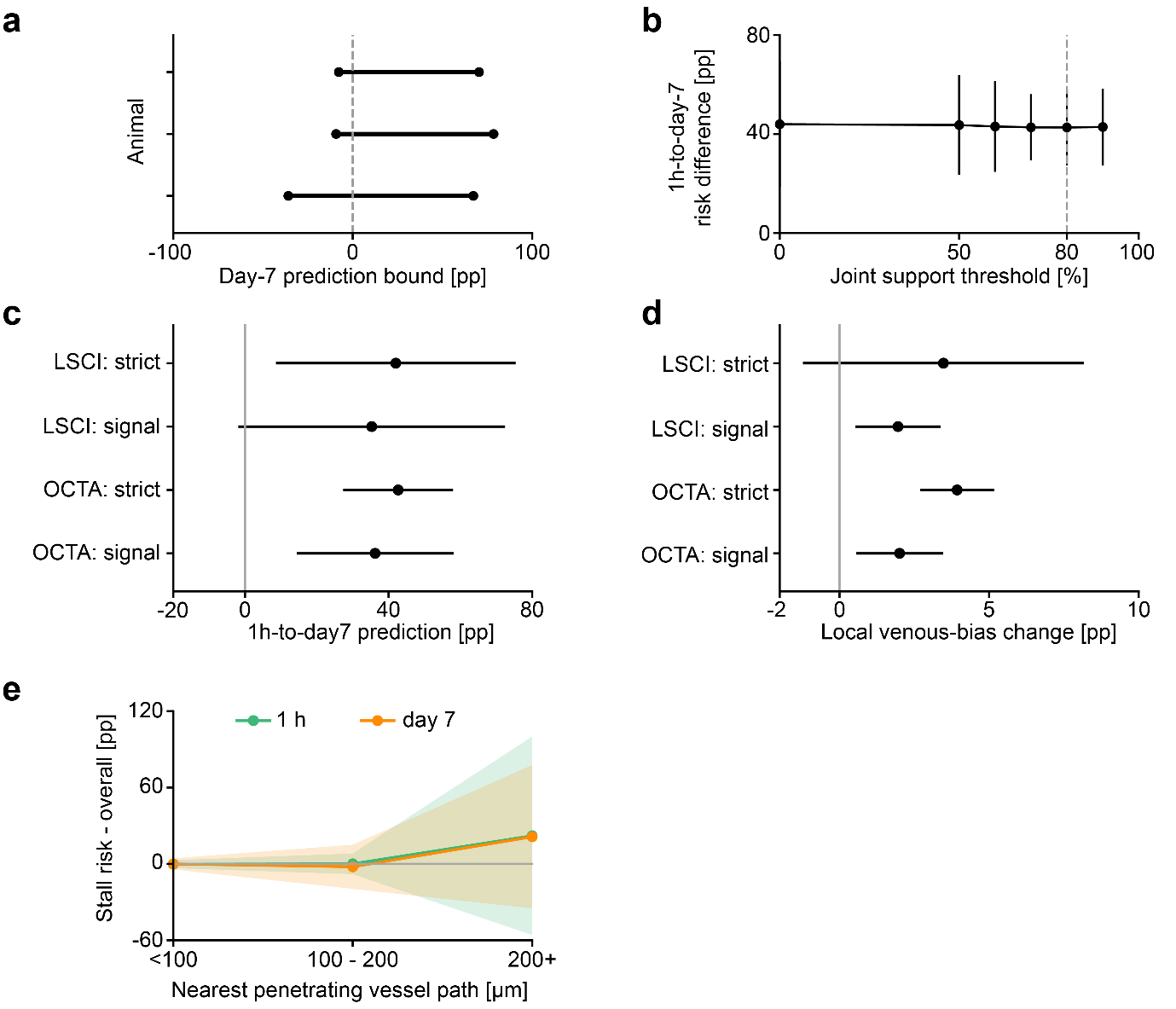

Figure S 15: Sensitivity of longitudinal stall patterns to measurement support, regional definitions and network distance. (a) Animal-specific worst-case bounds on the association between acute and day-7 stall status when indeterminate outcomes are allowed to vary. These are identification bounds, not confidence intervals. (b) Acute-to-day-7 stall-risk difference across joint imaging-support thresholds. The dashed line marks the selected 80% threshold. (c,d) Sensitivity of acute-to-day-7 prediction (c) and the change in local venous bias (d) to LSCI- or OCTA-based regional definitions and strict or strong-signal endpoint rules. (e) Stall risk relative to the animal’s overall risk at 1 h and day 7, grouped by baseline shortest network-path distance to a penetrating arterial or venous interface. Panels b-e show equal-animal means with 95% confidence intervals across three mice. Panel e uses the selected strict OCTA population with at least 80% joint support. Stall denotes loss of detectable OCTA perfusion. pp: percentage points.

**Supplementary Figure 16**

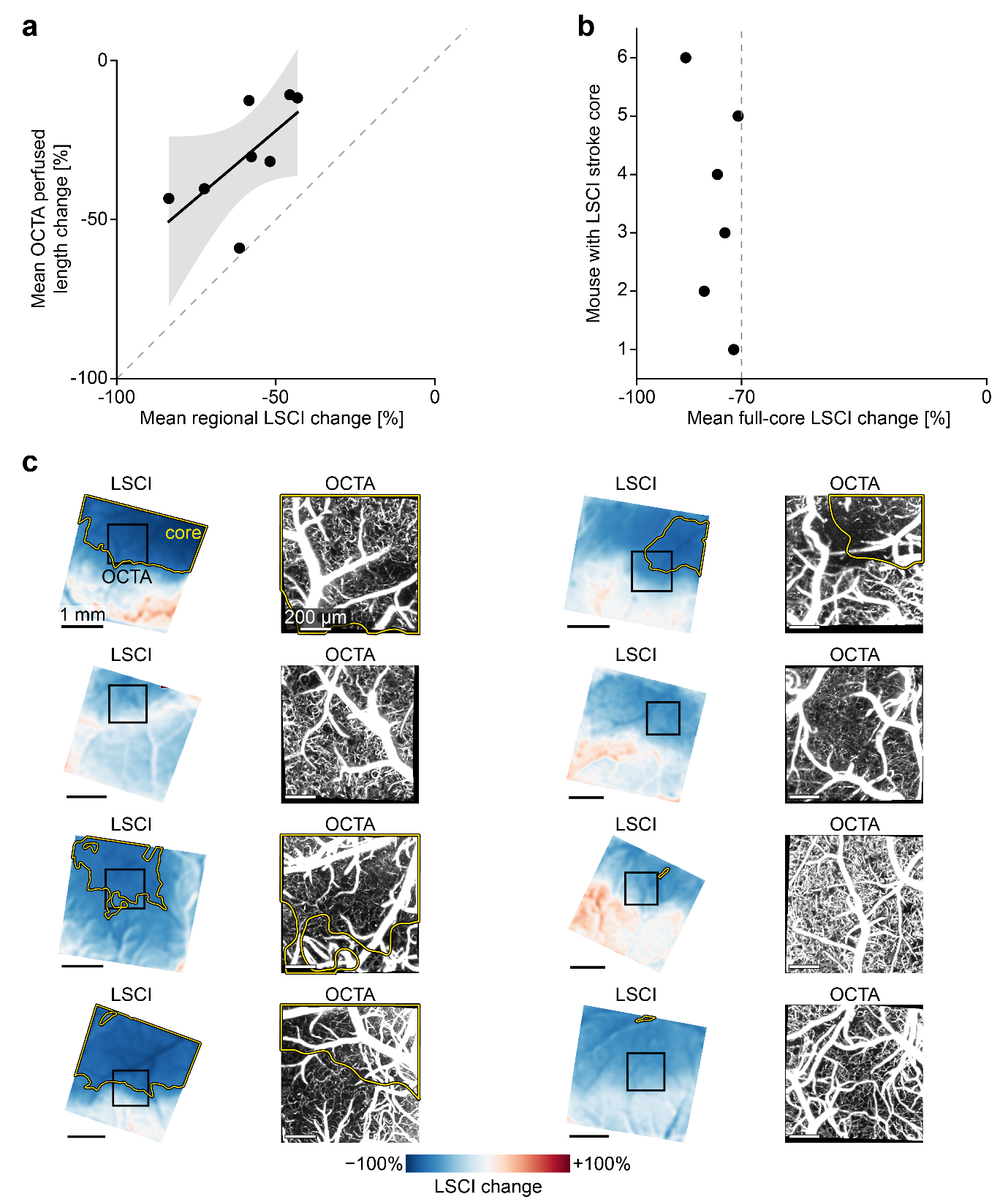

Figure S 16: Regional perfusion and imaging fields. a) Animal mean change in OCTA-detectable capillary length versus regional laser speckle contrast imaging (LSCI) change in the acute post-stroke assessment (n = 8 mice). The black line and gray band show the linear fit and pointwise 95% confidence interval; the dashed line indicates equality. (b) Mean LSCI change within the largest full-field perfusion core in six of these mice. Each point represents one animal; two mice had no core at the declared threshold. The dashed line marks a 70% reduction from baseline. (c) Paired full-field LSCI maps and registered acute OCTA projections for all eight mice. Black squares mark the OCTA field. Yellow outlines show the largest contiguous LSCI region with at least 70% perfusion reduction. Images without an outline have no defined core or a core outside the OCTA field. LSCI and OCTA were acquired at different times and measure different aspects of perfusion Scale bars 1 mm for LSCI and 200 µm for OCTA.

**Supplementary Figure 17**

Example of an OCTA maximum-intensity projection (MIP) illustrating the stroke core

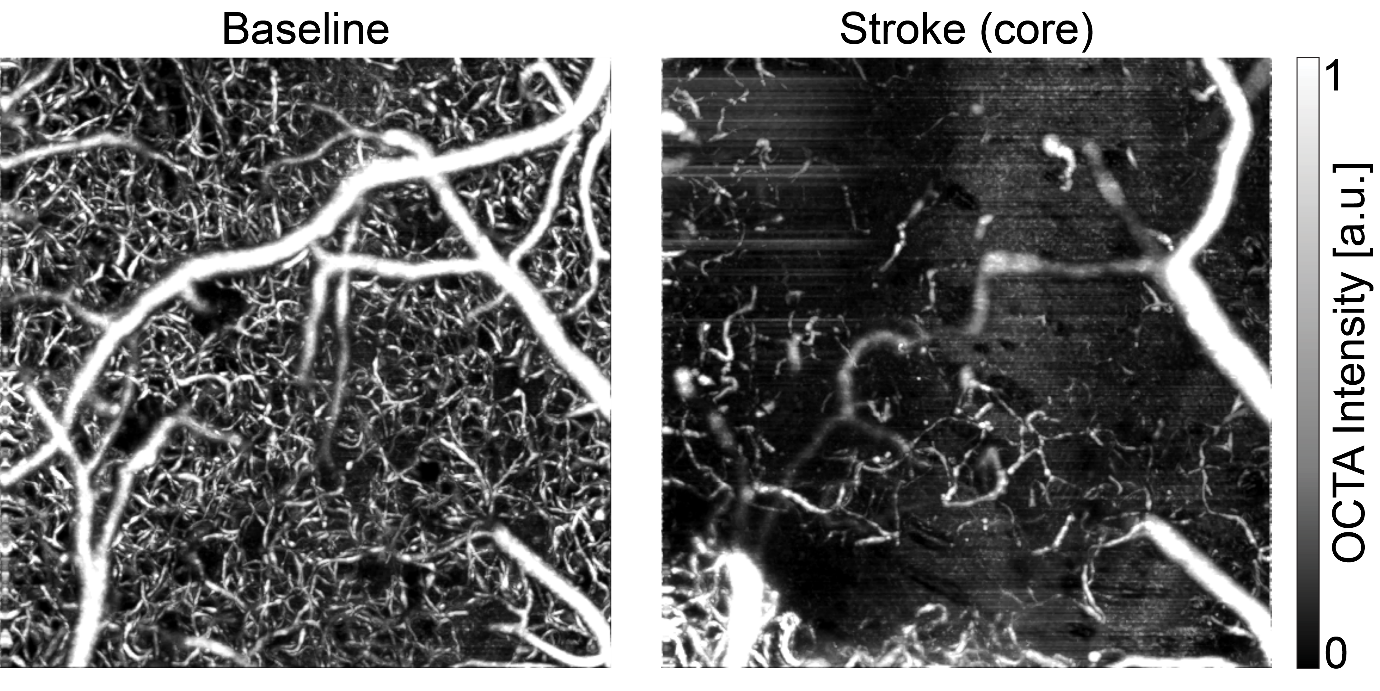

Figure S 17: Example of an OCTA maximum-intensity projection (MIP) illustrating the stroke core.
(left) OCTA MIP at baseline, displaying the feeding artery, capillary bed, and draining venous tree under normal perfusion. (right) OCTA MIP from the same region post-stroke, showing an almost complete absence of flow in the feeding artery and capillary bed, characteristic of the stroke core. The venous tree remains partially perfused and hence visible, with some vessels appearing dilated.

**Supplementary Tables 1 - 4**

**Supplementary Table 1. Animal metadata and analysis eligibility by acquisition group**

| **Group** | **Acquisition track** | **N** | **Age (weeks)** | **Sex** | **Strain** | **Acquired OCTA time points (n)** | **Animals included by analysis (n)** | **Exclusions and scope** |
| --- | --- | --- | --- | --- | --- | --- | --- | --- |
| 1 | Acute | 4 | 8-12 | Mixed  (2/2) | BALB/c | Baseline: 4 Acute: 4 | Acute: 4 Velocity: 4 Longitudinal: 0 LSCI regional: 0 | All four animals contributed to acute analyses. |
| 2 | Acute | 5 | 8-12 | male | BALB/c | Baseline: 5 Acute: 4 | Acute: 4 Velocity: 3 Longitudinal: 0 LSCI regional: 4 | One animal was excluded because the full field-of-view lost vessel contrast after stroke. A different animal was excluded due to velocity signal corruption at baseline |
| 3 | Longitudinal | 5 | 8-12 | female | BALB/c | Baseline: 5 Acute: 5 Day 7: 5 | Acute: 4 Velocity: 4 Longitudinal: 3 LSCI regional: 4 | One animal was excluded from paired analyses after registration failure. One further animal was excluded from day-7 tracking. |

**Supplementary Table 2. OMNIMap baseline measurements and literature comparisons**

| **Measurement** | **OMNIMap baseline measurement** | **Literature/reference value and citation** | **Reference preparation, method and sampling** | **Comparability** |
| --- | --- | --- | --- | --- |
| Capillary diameter | 3.85 ± 0.10 µm 12 animals, 26467 vessels/components Mean ± SD of animal medians. This study, Fig. 1 | 4.2 ± 0.4 µm Steinman et al. (2017) | In vivo two-photon fluorescence width, C57BL/6 mouse cortex under isoflurane. capillaries <8 µm and not penetrating vessels 4 mice. reported mean ± value. | Fluorescent vessel width and OCTA signal width use different contrast and fitting definitions. |
|  |  | 3.5 ± 0.2 µm Gould et al. (2017) | Ex vivo fluorescent-gel reconstruction of mouse somatosensory cortex, through cortical depth. 4 reconstructed cortical specimens, capillary distribution mean ± reported value. | Anatomical reconstruction. Preparation and caliber definition differ from in vivo OCTA. |
| Capillary velocity | 1.17 ± 0.36 mm/s 11 animals, 27724 vessels/components Mean ± SD of animal medians. This study, Fig. 1 | 1.0 ± 0.2 mm/s Pian et al. (2023) | DLS-OCT, isoflurane-anesthetized female C57BL/6 mouse barrel cortex, to 250 µm. strong-signal edges with axial tilt ≤35° 3 animal means, Table 1. mean ± reported value. | Centerline velocity, with angle/signal selection. Mean of animal means differs from OMNIMap animal medians and its moving-scatterer estimator. |
|  |  | 1.5 ± 1.2 mm/s Santisakultarm et al. (2012) | In vivo two-photon RBC centerline line scans, isoflurane-anesthetized C57BL/6 mouse cortex, imaging to 350 µm 156 vessels in 17 mice, reported pooled mean ± value. | Capillaries <10 µm. RBC centerline velocity and a pooled vessel summary differ from the OMNIMap moving-scatterer estimator and animal summaries. |
| AV:DA ratio | 3.16 ratio 12 animals, 731 AV / 231 DA components Pooled component ratio. This study, Fig. 2 | 3.0 ± 0.1 Blinder et al. (2013) | Ex vivo mouse vibrissa somatosensory cortex Mean ± SEM. 8 mice used for the penetrating-vessel spatial analysis. The ratio paragraph does not separately specify its animal n. | Blinder counts anatomical penetrations. OMNIMap counts labeled connected components within a finite field. |
|  |  | 2.75 ratio, pooled 151 AV / 55 DA components Our reanalysis of Schmid et al. (2017) | Three reconstructed mouse cortical vascular graphs. All full-network same-sign penetrating components, using the same connected-component counting approach. | Same component-counting approach, but full reference networks differ in lateral area and extent from the OMNIMap field.. |
| Nearest-vessel distances (k=1) | Artery→vein: 79.30 ± 13.28 µm Artery→artery: 110.95 ± 19.21 µm Vein→vein: 56.48 ± 5.87 µm 12 animals, mean ± SD of animal medians, k = 1. This study, Fig. 2 | Artery→vein: 97.25 ± 9.25 µm  Artery→artery: 79.08 ± 17.98 µm  Vein→vein: 55.46 ± 3.31 µm  Our reanalysis of Schmid et al. (2017) | 3 reference networks; 42 overlapping, field-matched crops (1000 × 1000 × 360 µm). Crop medians were averaged within each network, then summarized as mean ± SD across networks. | Reanalysis of Schmid et al. data using the same planar nearest-neighbor definition. Distances use the shallowest endpoint of each observed penetrating-vessel component; same-class comparisons exclude self. Artery→vein is directional. Differences in anatomical versus flow-based visibility and vessel labeling remain. |

**Notes**

DA: descending arteriole, AV: ascending venule, RBC: red blood cell, SD: standard deviation, SEM: standard error of the mean.

**Supplementary Table 3. Baseline predictors of subsequent capillary stalling without subtype**

| **Predictor** | **Animals** | **Capillaries** | **Stalls** | **Odds ratio [95% CI]** | **Raw P** | **Holm P** |
| --- | --- | --- | --- | --- | --- | --- |
| Baseline segment length | 12 | 18629 | 2564 | 1.689 [1.453, 1.964] | 9.9e-06 | 4.95e-05 |
| Baseline velocity | 11 | 16352 | 2112 | 0.782 [0.663, 0.923] | 0.00778 | 0.0156 |
| Baseline direct OCTA diameter | 12 | 16260 | 2180 | 1.061 [0.954, 1.18] | 0.243 | 0.243 |

**Notes**

Outcome: acute loss of detectable OCTA perfusion. Each row is a separate adjusted logistic model without subtype. Odds ratios are per within-animal SD of the log predictor.

All models adjust for depth, relative A-V position and boundary distance. Velocity and diameter models additionally adjust for log segment length. Continuous terms are standardized within animal.

Animal coefficients are combined by random-effects REML meta-analysis with Knapp–Hartung 95% intervals. Animals are independent replicates. Capillary and stall counts describe contributing cases.

Holm adjustment retains the original five-predictor family for length and two-predictor family for velocity/diameter

**Supplementary Table 4. Jointly adjusted capillary subtype and baseline predictors of stalling**

| **Model term** | **Odds ratio [95% CI]** | **Raw P** | **Holm P (eight terms)** |
| --- | --- | --- | --- |
| Velocity | 0.741 [0.587, 0.935] | 0.0171 | 0.0998 |
| Diameter | 1.219 [0.996, 1.491] | 0.0535 | 0.214 |
| Segment length | 1.705 [1.434, 2.027] | 6.5e-05 | 0.00052 |
| Arterial BO | 1.247 [1.097, 1.418] | 0.00368 | 0.0257 |
| Venous BO | 1.172 [0.934, 1.472] | 0.149 | 0.446 |
| D-to-C vs other | 1.392 [1.079, 1.797] | 0.0166 | 0.0998 |
| Tissue depth | 1.101 [0.83, 1.462] | 0.46 | 0.92 |
| Boundary distance | 0.918 [0.705, 1.195] | 0.481 | 0.92 |

**Notes**

All coefficients use the same population: 10 animals, 6,061 capillaries and 694 acute stalls.

Outcome: acute loss of detectable OCTA perfusion. The joint logistic model includes all eight listed terms and an intercept. Diameter, segment length and velocity are log-transformed. All continuous terms are standardized within animal. D-to-C is compared with all other subtypes. BO denotes branch order.

Animal coefficients are combined by random-effects REML meta-analysis with Knapp–Hartung 95% intervals. Animals are independent replicates.

The displayed Holm P values adjust across all eight terms. The separate prespecified two-model D-to-C family (branch order versus physical-path adjustment) gives Holm P = 0.0281. These correction families differ.
